# Positive and negative feedback regulation of the TGF-β1–SMAD4 axis explains two equilibrium states in human skin aging

**DOI:** 10.1101/2023.07.18.546970

**Authors:** Masatoshi Haga, Keita Iida, Mariko Okada

## Abstract

Skin homeostasis during aging is critical not only for the appearance but also biological defense of the human body. In this study, we identified thrombospondin-1 (THBS1) and fibromodulin (FMOD) as positive and negative regulators, respectively, of the TGF-β1–SMAD4 axis in human skin aging based on *in vitro* and *in vivo* omics analyses and mathematical modeling. Transcriptomic and epigenetic analyses of senescent dermal fibroblasts identified TGF-β1 as the key upstream regulator. Bifurcation analysis identified a binary senescent/non-senescent switch, with THBS1 as the main controller. Sensitivity analysis of the TGF-β1 signaling pathway indicated that THBS1 expression was sensitively regulated while FMOD was robustly regulated, suggesting that THBS1 is a controllable factor. Inhibition of SMAD4 complex formation was experimentally validated as a promising manner to control THBS1 production and senescence. This study demonstrates the potential of a data-driven mathematical approach in determining the mechanisms of skin aging.

**Graphical abstract:** 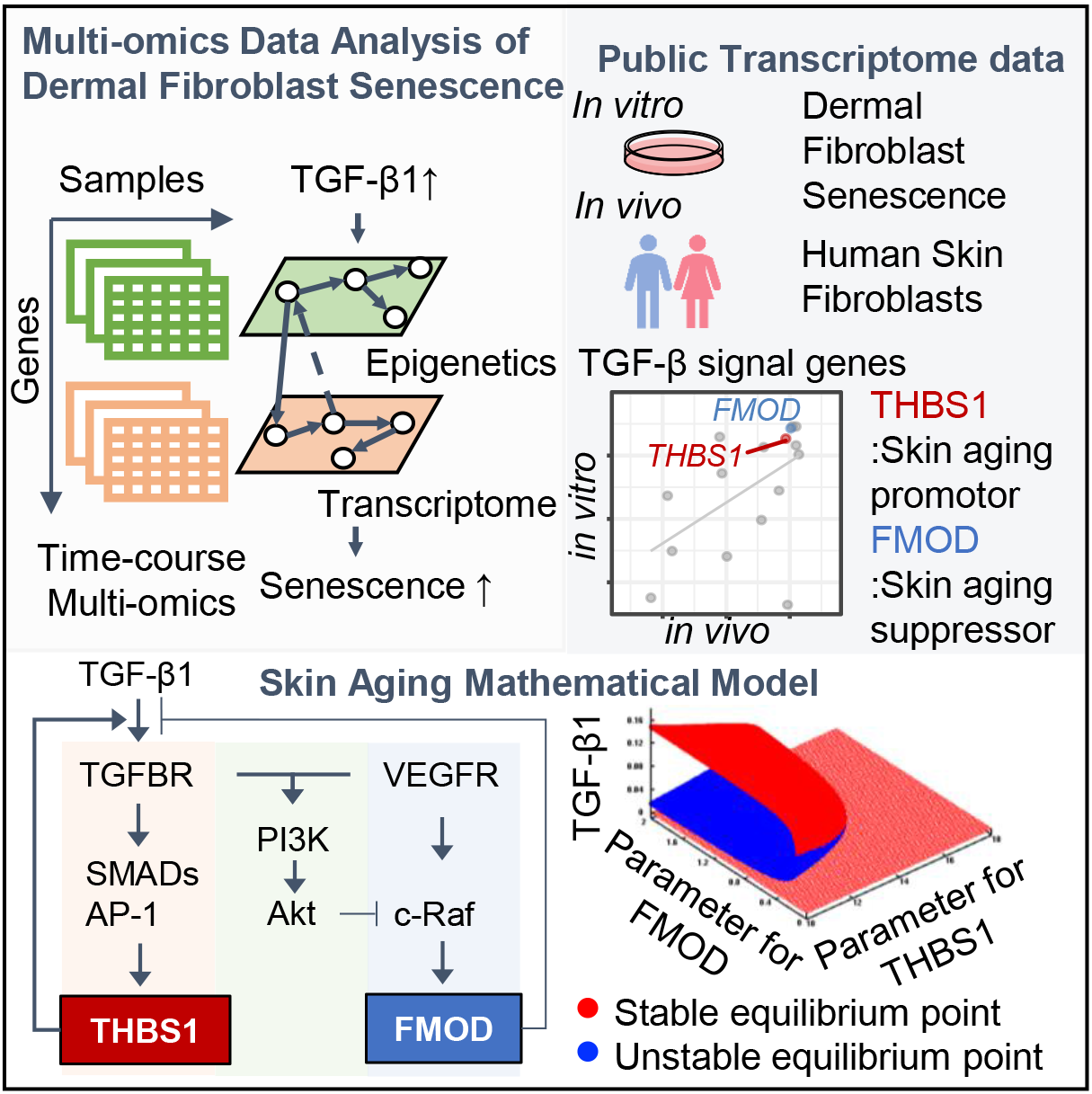

**eTOC Blurb:** Haga, Iida and Okada revealed that the balance between FMOD and THBS1 determines the two equilibrium states of skin homeostasis and THBS1 is a controllable factor in skin aging

**Highlights:**

- Multi-omics analysis identified TGF-β1–SMAD4 as key regulators in human skin aging
- THBS1 and FMOD promoted and suppressed factors in skin aging, respectively
- THBS1 controlled the senescent and non-senescent states of the aging switch
- SMAD4 is a potential target for inhibiting THBS1 expression and cell senescence

## INTRODUCTION

Aging is an inevitable and irreversible physiological phenomenon that disrupts the homeostasis of the human body and rapidly increases the risk of diseases, such as skin cancer^1, 2^. The skin is the largest organ of the human body that acts as a barrier to the external environment and protects against unwanted external stimuli^3^. Maintaining skin homeostasis during aging is important not only for physical appearance but also biological defense, as the skin becomes thinner and more fragile, and the development of skin diseases is accelerated^4^.

In general, skin aging is caused by both external (e.g., UV exposure, smoking, environmental pollution)^5^ and internal factors (e.g., impaired skin repair, hormones)^6^. Cellular senescence is a critical internal factor of skin aging^7, 8^. Skin tissue consists mainly of the epidermis and dermis. Keratinocytes, which make up the epidermis, have less influence on senescence because they have a higher turnover than fibroblasts in the dermis^9^. In the dermis, skin aging results in the accumulation of senescent fibroblasts^10–13^. Fibroblasts orchestrate the development of a functional skin barrier through crosstalk between the dermis and epidermis^14^. Incorporation of senescent fibroblasts into a 3D equivalent model of human skin can reproduce typical changes associated with skin aging^15, 16^. Early or late passage fibroblasts could mimic young and aged skin, respectively, and the late passage fibroblasts thinned the dermis and induced epidermal differentiation^15^. Another 3D skin model using oxidative stress-induced premature senescent fibroblasts caused progressive thinning of the epidermis with an increase in premature senescent cells in the dermis^16^. These studies strongly suggest that regulation of fibroblast senescence is fundamental to maintaining the skin barrier and aging.

Senescent cells exhibit irreversible terminal growth arrest with flattened large-cell morphology, shortened telomeres, increased levels of the cyclin-dependent kinase inhibitors p16 or p53/p21, and secretion of the senescence-associated secretory phenotype (SASP) consisting of inflammatory cytokines, such as interleukin (IL)-6 and IL-8^9, 17^. Senescence-associated β-galactosidase (SA-β-Gal) activity is another well-known biomarker of cellular senescence. Various stressors (e.g., replication stress, DNA double-strand breaks, telomere shortening, oncogene activation, reactive oxygen species) induce cellular senescence^18^. However, cellular senescence has not yet been fully elucidated at the systems level.

Data-driven multi-omics approaches have been used in various studies to elucidate the complex mechanisms of skin aging^19, 20^. RNA-seq, DNA methylation, histone methylation (histone H3 lysine 4 tri-methylation [H3K4me3] and histone H3 lysine 27 tri-methylation [H3K27me3]) from healthy skin fibroblasts of donors aged 35 to 75 years revealed an age-dependent decrease in the expression of genes involved in translation and ribosomal function^19^. An assay for transposase-accessible chromatin with high-throughput sequencing (ATAC-seq) and RNA-seq analyses of healthy and Hutchinson–Gilford progeria syndrome (HGPS, a premature aging syndrome) skin fibroblasts revealed altered chromatin accessibility enriched in lamina-associated domains (LAD) and HGPS-specific gene expression^20^. Although multi-omics approaches can capture the landscape of gene regulation, it remains difficult to identify key factors and regulatory structures among the thousands of genes involved in skin aging. Mathematical modeling combined with omics analysis provides a potential solution to enhance our understanding of the key mechanisms of skin aging controlled by genetic and environmental factors^21^. Mathematical models have been used to represent human disease as a gene regulatory network and stratify patients, identify key regulators, and predict drug targets^22–24^. As the use of animals in skin cosmetic research is prohibited in the industry^25^, insights from simulations with mathematical models are important to study skin aging.

In this study, the mechanisms of human skin aging were identified through multi-omics analysis of *in vitro* and *in vivo* data and mathematical modeling. Replication stress was introduced into human dermal fibroblasts, HFF-1 cells—the senescence cell model. Analyses of RNA-seq, chromatin immunoprecipitation sequencing (ChIP-seq) for histone H3 lysine 27 (H3K27Ac) modification, and ATAC-seq of different passages of HFF-1 cells and two independent *in vitro* and *in vivo* RNA-seqs of fibroblasts^26, 27^ identified transforming growth factor β1 (TGF-β1) as a key regulator of internal and external aging. The levels of thrombospondin-1 (THBS1) and fibromodulin (FMOD) were highly correlated with population doubling level (PDL) or age both *in vitro* and *in vivo*. THBS1 is known to activate the latent form of TGF-β1^28^ while FMOD binds to TGF-β1 and inhibits its binding to the TGF-BR receptor^29–31^. Inhibitor and knockdown (KD) assays implicated the signaling network for THBS1 and FMOD production. Finally, mathematical modeling and bifunctional analysis revealed a regulatory network mechanism by TGF-β1, THBS1, and FMOD in skin aging. This study provides important insights into the network mechanisms that drive and control skin aging and senescence.

## RESULTS

### Multi-omics analysis identifies TGF-β1 as potential regulator of skin aging

To identify global transcriptional signatures and their regulatory mechanisms in dermal fibroblast senescence, we performed RNA-seq, ChIP-seq for H3K27Ac modification, and ATAC-seq of replicatively stressed HFF-1 cells (Figure 1A). First, HFF-1 and BJ, a different cell line of human dermal fibroblasts, were maintained from early (HFF-1: PDL 13, BJ: PDL 23) to late (HFF-1: PDL 53, BJ: PDL 61) PDL to induce replication stress (Figure 1B). We examined the activities of p53, p21, and SA-β-gal, molecular markers known to increase with senescence, for different PDLs of fibroblast cells (HFF-1: Figure 1C, 1D; BJ: Figure S1A, S1B). Replication stress-induced cellular senescence was promoted by increased PDL in both cell lines. To identify transcription factors (TFs) that regulate senescence in HFF-1, the TF enrichment score was calculated using DoRothEA^32^ for the top 20 TFs (Figure 1E). SMAD3 and SMAD4, downstream TFs of the TGF-β pathway, were enriched with increasing PDL. We also identified known senescence-associated TFs, including JUNB^33^, GATA6^34^, TP53^35^, TEAD1^33^, and E2F4^33^. To confirm whether SMAD motifs are indeed enriched with cellular senescence, we performed a TF motif enrichment analysis using HOMER^36^, with which we compared the gained ATAC peaks—the open chromatin regions specifically enriched for late PDL (Figure 1F). SMAD2 (–log[adj-p value]: 15.8) and SMAD4 (–log[adj-p value]: 10.3) were associated with the gained ATAC peaks. We analyzed whether TGF-β is the upstream regulator of cellular senescence in dermal fibroblasts from 93 overlapping differentially expressed genes (DEGs) identified in the RNA-seq, differential ATAC peaks, and differential H3K27Ac peaks (Figure 1G) between PDL 24, PDL 36, and PDL 47. TGF-β1 was identified as the top regulator among the overlapping DEGs. From the 215 common genes shared between upregulated DEGs and genes annotated from differentially gained peaks of H3K27Ac ChIP-seq, indicating active enhancer regions, TGF-β1, TGF-β3, and SMAD3 were the top three upstream regulators (Figure S2A). These results indicate that TGF-β1 plays a critical role in the senescence of dermal fibroblasts.

**Figure 1.**
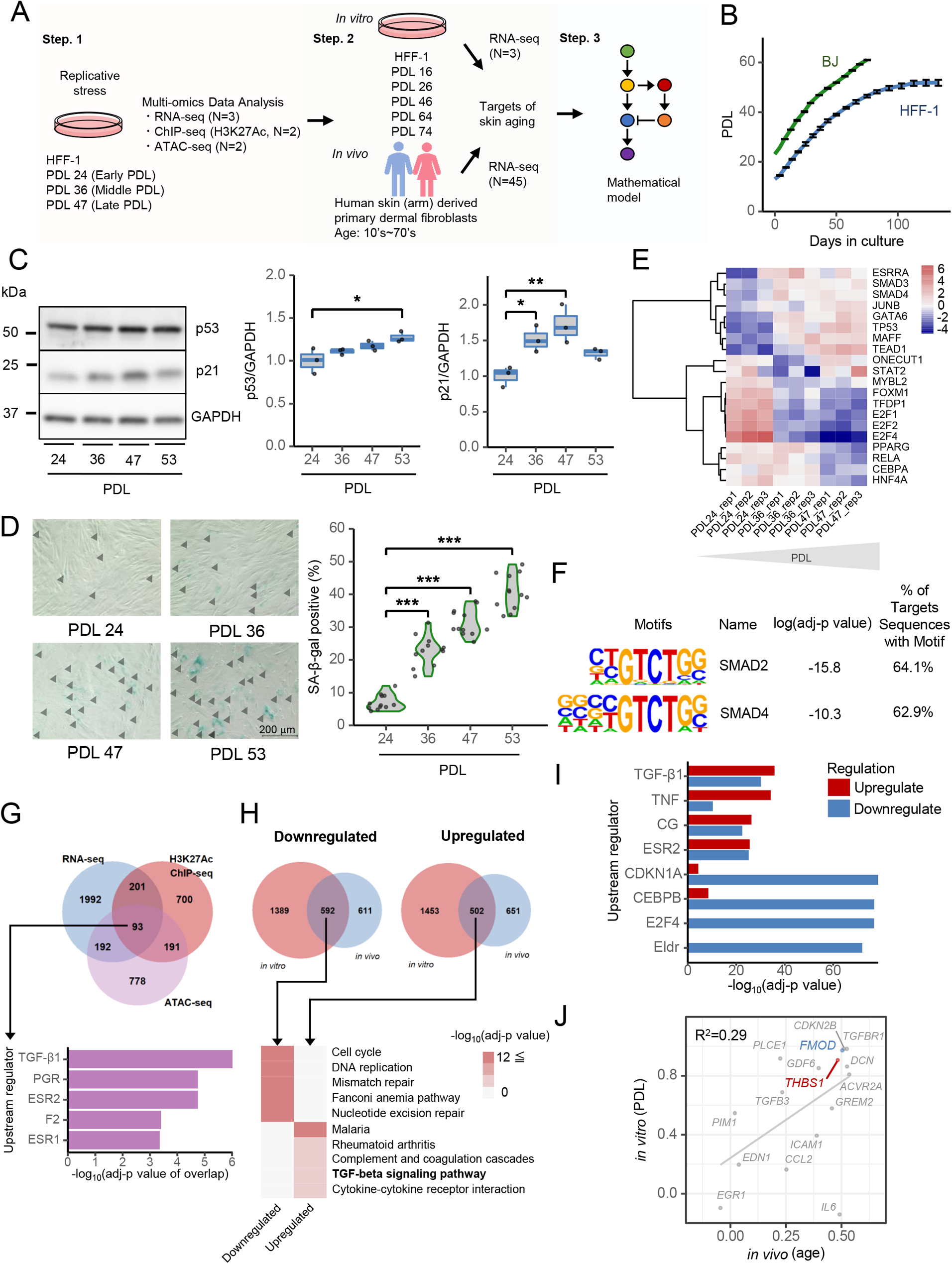
Integrated multi-omics analysis identifies TGF-β1 as central modulator of skin aging and senescence. A) Workflow of skin aging and senescence analyses. In step one, replication stress was induced by passage culturing using human foreskin fibroblasts, HFF-1, to prepare each PDL (see STAR Methods), and generate RNA-seq, ChIP-seq with H3K27Ac antibody, and ATAC-seq data. In the second step, two independent public RNA-seq datasets were analyzed, one *in vitro*^27^ and another *in vivo*^26^. Finally, in the third step, we constructed a mathematical model based on the multi-omics analysis and *in vitro* experimental results. B) Growth curve of HFF-1 (blue line) and BJ cells (green line). N = 3, mean (SD). C) Western blot of p53 and p21 in replication-stress-induced HFF-1 cells. (Left panel) Representative image. (Middle panel) Quantification of p53 expression. N = 3, * p < 0.05 (Dunnett’s test). (Right panel) Quantification of p21 expression. N = 3, * p < 0.05, ** p < 0.01 (Dunnett’s test). D) (Left panel) SA-β-gal staining of replication-stress-induced HFF-1 cells. SA-β-gal-positive cells are indicated by an arrowhead (black). (Right panel) Quantification of SA-β-gal: SA-β-gal-positive rate (%) = number of SA-β-gal positive cells / total number of cells × 100, N = 3 (4 points/well), *** p < 0.001 (Dunnett’s test). E) Transcription factor (TF) enrichment analysis of RNA-seq data derived from replication-stress-induced HFF-1 cells. The heatmap shows the normalized TF enrichment scores calculated using the DoRothEA. F) Enriched motifs in the gained ATAC-seq peaks with increase of PDL. log(adj-p value) and proportion of target sequences with motif was calculated using the “findMotifsGenome.pl” function of HOMER. G) (Upper panel) Venn diagram showing RNA-seq differentially expressed genes (DEGs; |fold change (FC)| > 1.2, adj-p < 0.05, blue shade), genes annotated from ATAC differential peaks (|log_2_FC| > 0, adj-p < 0.05, purple shade), and genes annotated from H3K27Ac differential peaks (|log_2_FC| > 0, adj-p value < 0.05, red shade) between PDL 24, PDL 36, and PD47. The number of genes in each condition is shown in the Venn diagram. (Lower panel) The top five upstream regulators using Ingenuity Pathway Analysis (IPA) are shown with epigenetic-linked DEGs (93 genes). A right-tailed Fisher’s exact test was used to calculate –log(adj-p) of overlap. H) (Upper panel) Venn diagram showing the public *in vitro* DEGs (blue shade) and *in vivo* DEGs (red shade). (Lower panel) Heat map showing top five pathways annotated in the pathway enrichment analysis (Kyoto Encyclopedia of Genes and Genomes [KEGG]). The adj-p value was calculated using “compareCluster” of clusterProfiler. I) Upstream regulators using common genes between the public *in vitro* and *in vivo* datasets. The top four upstream regulators of 592 downregulated and 502 upregulated genes are shown with either a red or blue bar. A right-tailed Fisher’s exact test was used to calculate the adj-p value of overlap. J) Correlation analysis between the public *in vitro* (PDL), the public *in vivo* (age), and gene expression data. Spearman correlations were calculated using the “cor” function of R software; *THBS1* (red) and *FMOD* (blue). R^2^ was calculated using the “ggpmisc::stat_poly_eq” function.

To further confirm the involvement of TGF-β1 in skin aging and senescence, two additional time-course-independent public RNA-seq datasets were analyzed: *in vivo* data for primary human dermal fibroblasts^26^ representing a wide age range (11–71 years of age, see donor list in Table S1), to analyze external factors, and *in vitro* data for HFF-1 cells over a long-term passage^27^, to analyze internal factors (PDL 16–74, five PDL points). The number of overlapping DEGs in both datasets corresponded to 592 downregulated and 502 upregulated genes (Figure 1H). Functional analysis of overlapping gene sets using the Kyoto Encyclopedia of Genes and Genomes (KEGG) database^37^ revealed that the cell cycle and TGF-β signaling pathways were enriched in both datasets. In addition, TGF-β1 was identified as the top regulator for the upregulated gene set by upstream analysis of the overlapping gene sets (Figure 1I). Comparisons between gene expression and *in vivo* age or *in vitro* PDL revealed that *THBS1* and *FMOD* expression were strongly correlated with skin aging and senescence (Figure 1J). In addition, the expression of the TGF-β receptor, *TGFBR1*, was also correlated with gene expression levels both *in vivo* and *in vitro*, indicating that TGF-β signaling is more likely to be activated in skin aging.

The percentage of TGF-β1-positive fibroblasts in the human dermis increases with age^38^. Higher PDL corresponded with higher TGF-β1 expression in HFF-1 cells (Figure 2A). Western blot (Figure 2B), qPCR (Figure S2F), and enzyme-linked immunosorbent assay (ELISA, Figure S2G) analyses revealed an increased and decreased expression of THBS1 and FMOD, respectively, in higher cell passages of dermal fibroblasts. The expression of THBS1 was confirmed to increase with age in human dermal tissue isolated from female donors (23–63 years of age; Figure 2C, see Table S2 for donor list). From the same donors, p21 expression was also confirmed to increase with age, suggesting that our model of cellular senescence induced by long-term passaged culturing accurately captured changes in human skin-factors associated with aging. Note that THBS1 was not detected in the epidermis, suggesting that THBS1 functions in the dermis during skin aging. Interestingly, using publicly available transcriptome data^26^ (see Table S3 for donor list), *THBS1* expression was found to be increased in patients with HGPS (2–8 years of age) compared to that in healthy controls (1–9 years of age) of the same age (Figure S2H). The results from our data-driven analysis identified the TGF-β1–SMAD signaling pathway along with THBS1 and FMOD expression as critical factors regulating skin aging and dermal fibroblast senescence.

**Figure 2.**
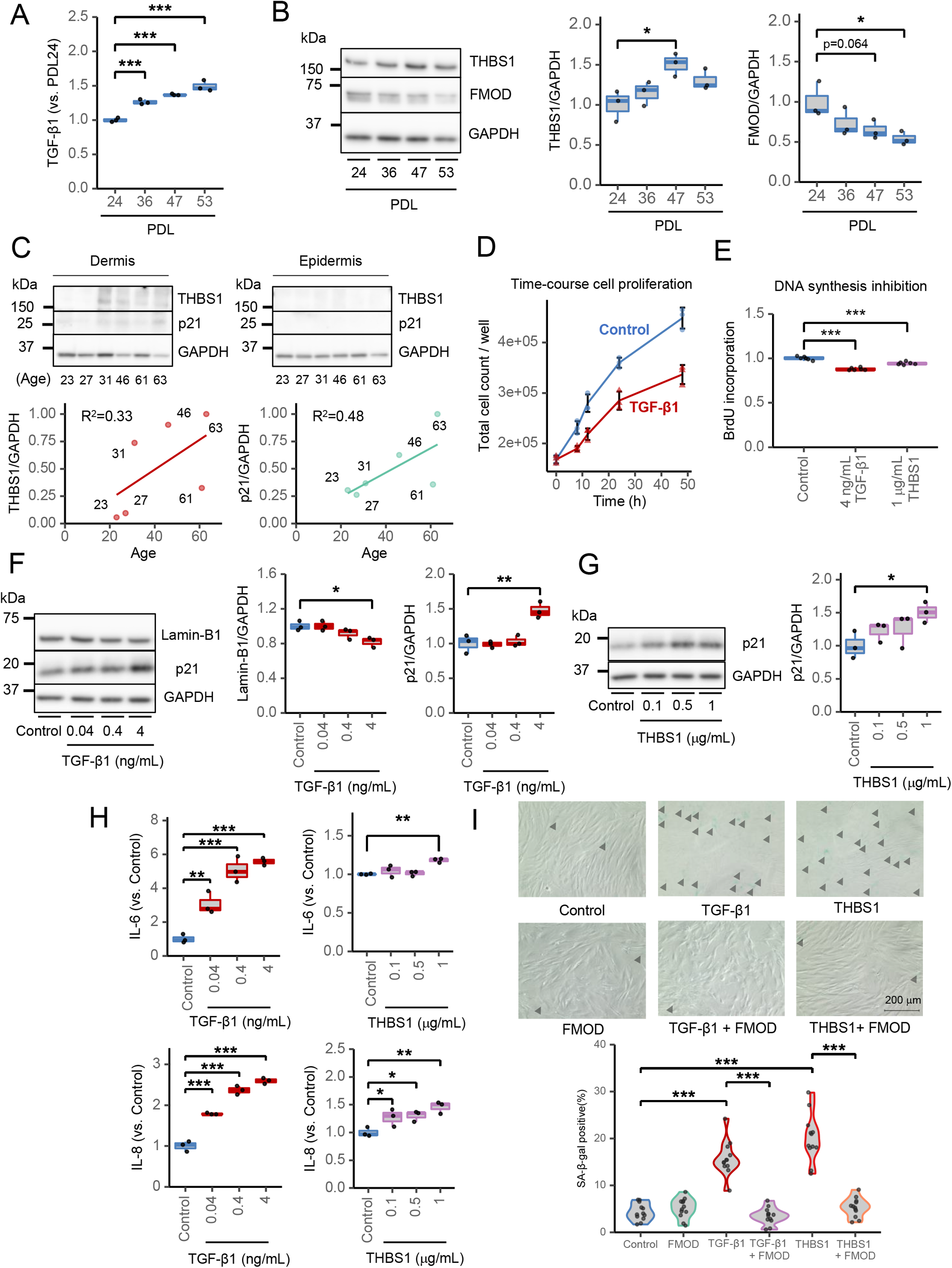
TGF-β1 and THBS1 act as senescence-inducing factors, while FMOD acts as a senescence inhibitor. A) TGF-β1 ELISA in replication-stress-induced HFF-1 cell supernatant. N = 3, *** p < 0.001 (Dunnett’s test). B) Western blot analysis of THBS1 and FMOD in replication-stress-induced HFF-1 cells. (Left panel) Representative image. (Middle panel) Quantification of THBS1 expression. N = 3, * p < 0.05 (Dunnett’s test). (Right panel) Quantification of FMOD expression. N = 3, * p < 0.05 (Dunnett’s test). C) Western blot analysis of THBS1 and p21 in human dermal and epidermal tissues (N = 6). The data were normalized to the maximum (1) value. R^2^ was calculated using the “stat_poly_eq” function of R. (Upper panel) The age of each donor is displayed in the bottom of the image (see donor list in Table S2). (Lower left panel) Quantification of THBS1 expression in dermis. (Lower right panel) Quantification of p21 expression in dermis. D) Time-course cell proliferation analysis of control (blue-circled dots) or 4 ng/mL TGF-β1-treated (red triangular dots) cells stained with trypan blue. N = 3, mean (SD). E) BrdU incorporation assay. Control (blue), 4 ng/mL TGF-β1 (red), or 1 μg/mL THBS1 (purple). F) Western blot of Lamin-B1 and p21 in TGF-β1-stimulated HFF-1 cells. Cell lysates were collected 48 h after control (blue) or TGF-β1 (red) treatment. (Left panel) Representative image. (Middle panel) Quantification of Lamin-B1. N = 3, * p < 0.05 (vs. Control, Dunnett’s test). (Right panel) Quantification of p21. N = 3, *** p < 0.001 (Dunnett’s test). G) Western blot of p21 in THBS1-stimulated HFF-1 cells. Cell lysates were collected 48 h after control (blue) or THBS1 (purple) treatment. (Left panel) Representative image. (Right panel) Quantification of p21. N = 3, * p < 0.05 (Dunnett’s test). H) IL-6 and IL-8 ELISA in TGF-β1- or THBS1-stimulated HFF-1 cells. Cell supernatants were collected 48 h after TGF-β1 (red) or THBS1 (purple) treatment. (Upper left panel) Quantification of IL-6 ELISA with TGF-β1 treatment. N = 3, ** p < 0.01, *** p < 0.001 (Dunnett’s test). (Upper right panel) Quantification of IL-6 ELISA with THBS1 treatment. N = 3, ** p < 0.01 (Dunnett’s test). (Bottom left panel) Quantification of IL-8 ELISA with TGF-β1 treatment. N = 3, ** p < 0.01, *** p < 0.001 (Dunnett’s test). (Bottom right panel) Quantification of IL-8 ELISA with THBS1 treatment. N = 3, ** p < 0.01 (Dunnett’s test). I) Effect of TGF-β1 or THBS1 treatment and inhibition by FMOD on SA-β-gal activity. HFF-1 cells were treated with control, TGF-β1 (4 ng/mL), THBS1 (0.5 μg/mL), or FMOD (8 ng/mL). Combinations of TGF-β1 (4 ng/mL) and FMOD (8 ng/mL) as well as THBS1 (0.5 μg/mL) and FMOD (8 ng/mL) were performed alongside stand-alone treatments. (Left panel) Representative images. SA-β-gal-positive cells are shown with an arrowhead (black). Scale bars, 200 μm. (Right panel) Quantification of SA-β-gal: SA-β-gal-positive rate (%) = number of SA-β-gal-positive cells / total number of cells × 100, N = 3 (4 points/well), *** p < 0.001 (Tukey’s multiple comparisons).

### TGF-β1 and THBS1 induce senescence of human dermal fibroblasts

We investigated the biological functions of TGF-β1 and THBS1 in terms of the senescence of human dermal fibroblasts. We first assessed the proliferation of HFF-1 cells (Figure 2D) and found that TGF-β1 treatment suppressed HFF-1 growth over time (8 h, 12 h, 24 h, and 48 h) compared to that of untreated cells. As with TGF-β1, THBS1 treatment resulted in a concentration-dependent decrease in cell viability (Figure S2J). TGF-β1 and THBS1 treatments were confirmed to inhibit DNA synthesis by BrdU incorporation experiments (Figure 2E). The levels of the senescence markers Lamin-B1 decreased and p21 increased in a TGF-β1 dose-dependent manner (Figure 2F). Similar to the TGF-β1 treatment, THBS1 enhanced p21 expression (Figure 2G), confirming that TGF-β1- or THBS1-dependent suppression of cell proliferation is caused by cellular senescence. TGF-β1 or THBS1 treatment of HFF-1 cells induced the production of the pro-inflammatory SASPs IL-6 and IL-8, which are known to be secreted with cellular senescence^39, 40^ (Figure 2H). TGF-β1 treatment also decreased Lamin-B1 expression (Figure S1D) and increased IL-6, IL-8 (Figure S1E), and SA-β-gal production (Figure S1F) in BJ cells. In addition, treatment with TGF-β1 or THBS1 significantly increased the number of SA-β-gal-positive cells (Figure 2I). Interestingly, FMOD alone did not affect SA-β-gal activity; however, in combination treatments, FMOD suppressed the effects of TGF-β1 or THBS1 on SA-β-gal activation. These results strongly suggest that TGF-β1 and THBS1 promote the senescence of skin fibroblasts and this effect is suppressed by FMOD.

### THBS1 and FMOD expression is controlled by the TGF-β1 signaling pathway

THBS1 is known to activate latent TGF-β1^28^, while FMOD reportedly binds to TGF-β1 to inhibit its binding to the TGF-BR receptor^29–31^. THBS1 is known to be induced by TGF-β1 stimulation in human dermal fibroblasts^41^. Consistently, we found that THBS1 expression was induced by TGF-β1 in HFF-1 cells (Figure 3A). However, the same TGF-β1 treatment decreased FMOD expression. These results were also confirmed in BJ cells (Figure S1C, S1D). Since THBS1 is positively regulated by TGF-β1, the effect of FMOD on THBS1 expression was examined in HFF-1 cells (Figure 3B). While TGF-β1 alone increased THBS1 expression, the addition of FMOD completely suppressed THBS1 levels. In contrast, THBS1 treatment downregulated FMOD expression in a dose-dependent manner (Figure 3C). These findings suggest a mutual inhibitory role for THBS1 and FMOD. Interestingly, stimulation of BJ cells with THBS1 enhanced its own expression (Figure 3D), suggesting the positive-feedback regulation of the TGF-β pathway by THBS1 in dermal fibroblasts.

**Figure 3.**
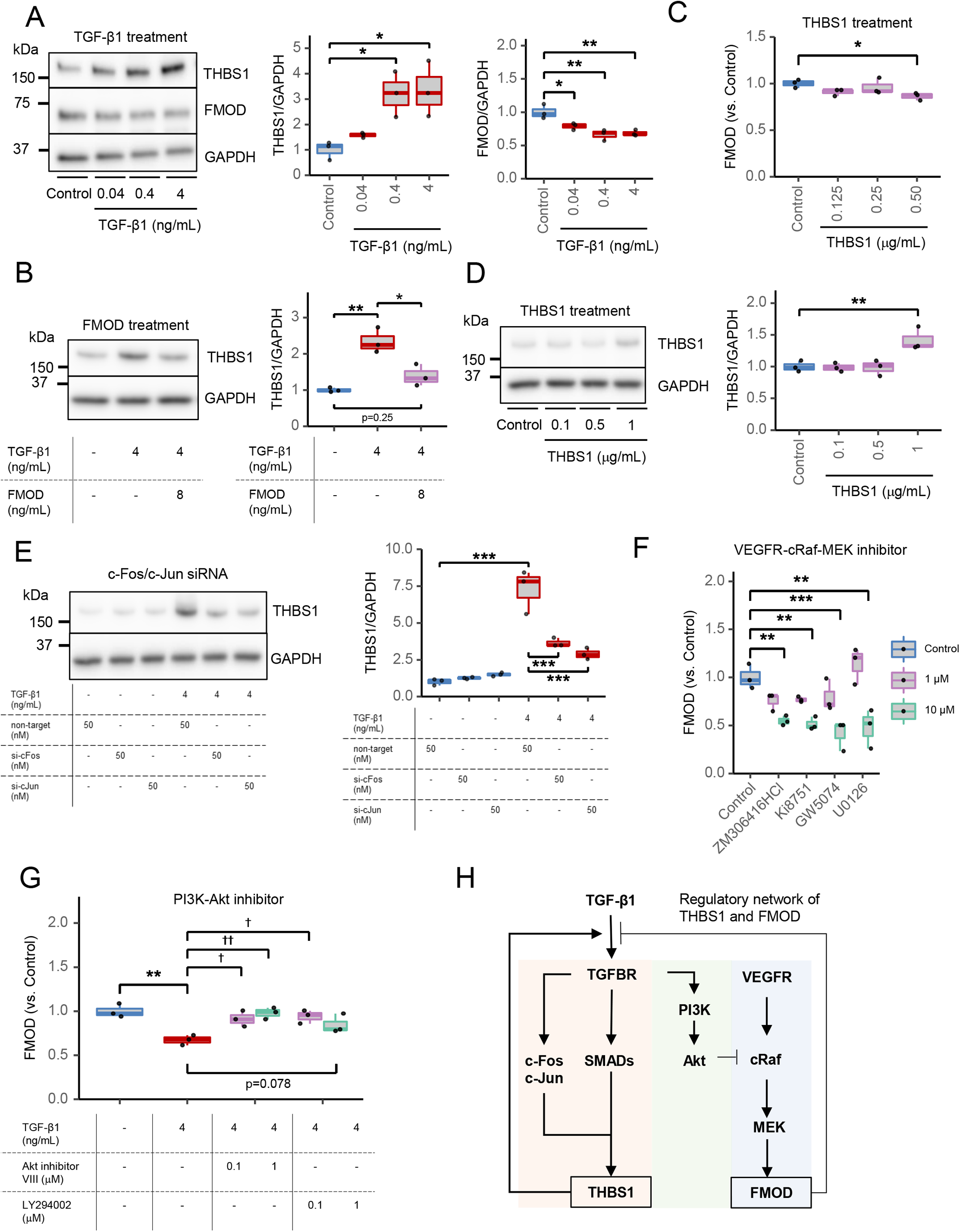
Regulatory network of THBS1 and FMOD in skin aging. A) Western blot of THBS1 and FMOD in TGF-β1-stimulated HFF-1 cells. Cell lysates were collected 48 h after control (blue) or TGF-β1 (red) treatment. (Left panel) Representative image. (Middle panel) Quantification of THBS1. N = 3, * p < 0.05 (vs. Control, Dunnett’s test). (Right panel) Quantification of FMOD. N = 3, * p < 0.05, ** p < 0.01 (Dunnett’s test). B) Western blot analysis of THBS1 in TGF-β1- and FMOD-stimulated HFF-1 cells. Cell lysates were collected 48 h after treatment with control (blue), 4 ng/mL TGF-β1 (red), or a combination of 4 ng/mL TGF-β1 and 8 ng/mL FMOD (purple). (Left panel) Representative image. (Right panel) Quantification of THBS1. N = 3, * p < 0.05, ** p < 0.01 (Tukey’s multiple comparisons). C) FMOD ELISA of THBS1-stimulated HFF-1 cells. Cells were treated with THBS1 (purple) for 48 h and their supernatants were analyzed. N = 3, * p < 0.01 (Dunnett’s test). D) Western blot analysis of THBS1 in THBS1-stimulated BJ cells. Cell lysates were collected 48 h after control (blue) or THBS1 (purple) treatment. (Left panel) Representative image. (Right panel) Quantification of THBS1. N = 3, ** p < 0.01 (Dunnett’s test). E) Western blot analysis of THBS1 with c-Fos/c-Jun knockdown (KD). Cell lysates were collected 48 h after control (blue) or TGF-β1 (red) treatment. (Left panel) Representative image. (Right panel) Quantification of THBS1. N = 3, ** p < 0.01 (Dunnett’s test). F) FMOD ELISA of kinase inhibitor-treated HFF-1 cells. Cells were treated with ZM306416HCl, Ki8751, GW5074, or U0126 for 48 h and their supernatants were analyzed. N = 3, ** p < 0.01, *** p < 0.001 (Dunnett’s test). G) FMOD ELISA in kinase inhibitor- and TGF-β1-treated HFF-1 cells. Cells were treated with Akt inhibitor VIII or LY294002 combined with TGF-β1 for 48 h and their supernatants were analyzed. N = 3, ** p < 0.01 (Student’s *t*-test), † p < 0.05, †† p < 0.01 (Dunnett’s test). H) Regulatory network of THBS1 and FMOD. In human dermal fibroblasts, TGF-β1 induced THBS1 production via TGF-βR–SMAD activation (red shade). FMOD was regulated via activation of the VEGFR–cRaf–MEK pathway (blue shade). Crosstalk between these pathways occurred with the TGF-β pathway inhibiting the VEGF pathway via the PI3K-Akt pathway (green shade).

We investigated the regulation of THBS1 and FMOD by TGF-β1 and related signaling pathways using small molecule inhibitors and small interfering (si) RNA knockdown (KD) experiments. TGF-β1-dependent induction of THBS1 and phosphorylation of SMAD2 and SMAD3 were inhibited by LY364947 (TGF-βRI/TGF-βRII inhibitor) and SB431542 (TGF-βRI inhibitor; Figure S3A). Since SMAD3 and SMAD4 cooperate with c-Fos/c-Jun of the activator protein-1 (AP1) family to mediate TGF-β-induced transcription^42^, we tested the involvement of c-Fos/c-Jun in the regulation of THBS1 using siRNA. c-Fos/c-Jun KD significantly reduced the TGF-β1-induced expression of THBS1 compared to that with non-targeted KD (Figure 3E). The AP-1 DNA binding inhibitor, T-5224, was also tested for its effect on THBS1 regulation in HFF-1 (Figure S3B). While T-5224 alone had no effect on THBS1 expression compared to TGF-β1 alone, the combination of TGF-β1 and T-5224 reduced THBS1 expression. We also confirmed that T-5224 and KD of c-Fos/c-Jun did not affect the activation of SMAD2, SMAD3, and SMAD4 (Figure S3C, S4B). These findings suggest that both SMAD activation and c-Fos/c-Jun binding to DNA, which is activated by TGF-β stimulation, are required for AND-gate network formation and THBS1 regulation.

FMOD expression was significantly inhibited by ZM306416 (VEGFRI inhibitor), Ki8751 (VEGFRII inhibitor), GW5074 (c-Raf1 inhibitor), and U0126 (MEK1/2 inhibitor; Figure 3F). To determine the mechanism of FMOD suppression by TGF-β1, we tested whether the combination of kinase inhibitor and TGF-β1 would restore FMOD expression (Figure 3G). Akt inhibitor VIII (Akt inhibitor) or LY294002 (PI3K inhibitor) significantly restored FMOD expression in the presence of TGF-β1. These results indicate that THBS1 and FMOD expression is regulated by the TGF-β and VEGF-Raf-ERK signaling pathway, respectively (Figure 3H), and that the TGF-β1-dependent suppression of FMOD is mediated by the PI3K-Akt pathway.

### Mechanistic mathematical model of skin aging

A nonlinear ordinary differential equation (ODE) model of the core transcription factor network consisting of TGF-β1, THBS1, and FMOD was constructed to elucidate the regulatory network at the systems level (Figure 4A, see details in the Methods section). Fitting against real THBS1 and FMOD levels at different TGF-β1 concentrations yielded parameters that reproduced the experimental results (Figure S5A). Experimental evidence suggests that THBS1 and FMOD induce and suppress senescence, respectively; therefore, it is likely that FMOD expression is suppressed in senescent cells through the formation of a positive feedback of THBS1 production. Accordingly, we examined the steady state solutions by controlling the THBS1 regulation parameter *K*_1_ and FMOD regulation parameter *K*_2_, while the TGF-β1 regulation parameter *c*/*d*_3_ was arbitrarily fixed to 1. When *K*_1_ is low and *K*_2_ is high, TGF-β1 expression is promoted via a saddle-node bifurcation, which exhibits a bistable switch (Figure 4B). Since the expression of TGF-β1 was upregulated in the senescent state (Figure 2A), changes in these parameters simulate the transition from a non-senescent to senescent state. Importantly, the bistable region was quite asymmetric with respect to *K*_1_ and *K*_2_ (Figure 4C). For any fixed *K*_2_, changes in *K*_1_ will induce a transition from the current state. On the contrary, *K*_2_ will induce the state transition for a small parameter range of *K*_1_. This indicates that *K*_1_ has more influence on state transitioning.

**Figure 4.**
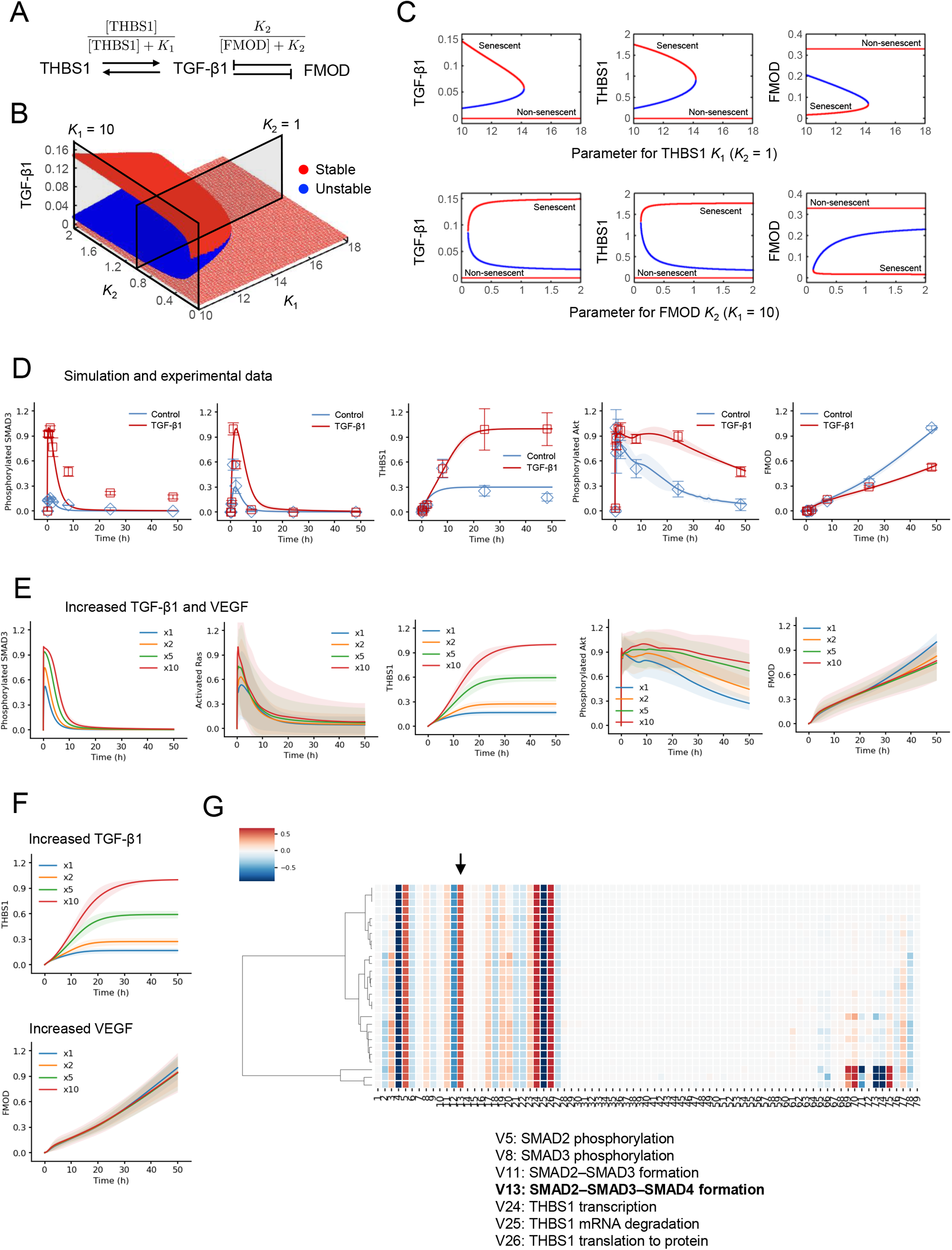
Experimental data-based mathematical modeling and simulation of skin aging. A) Core transcription factor network consisting of TGF-β1, THBS1, and FMOD. Arrows and barred lines represent positive and negative regulations, respectively. Bifurcation diagram of the steady state expression of TGF-β1 with respect to *K*_1_ and *K*_2_. Note that [TGFβ1] ≡ 0 is a stable solution for *K*_1_ >0 and *K*_2_ > 0. B) Bifurcation diagrams of the steady state expressions of TGF-β1, THBS1, and FMOD, with respect to *K*_1_ (top panel) and *K*_2_ (bottom panel), which correspond to the cut surfaces shown in Figure 4B. Red and blue points represent stable and unstable points, respectively. Note that solutions with high TGF-β1 expression levels indicate a senescent state, while solutions with low TGF-β1 expression levels indicate a non-senescent state. C) Model parameters for the integrated TGF-β–VEGF model were trained on time-course phosphorylated SMAD3, c-Fos, THBS1, phosphorylated Akt, and FMOD protein levels obtained with or without TGF-β1 treatment. The points (Control: blue squares, TGF-β1: red squares) indicate experimental data, solid lines indicate the average simulation of 30 parameter sets, and shade areas indicate SD. The error bar represents SD. N = 3, mean (SD). Western blot images can be found in Figure S6A. D) Simulation of phosphorylated SMAD3, activated Ras (Ras-GTP), THBS1, FMOD, and phosphorylated Akt levels using integrated TGF-β–VEGF mathematical model. Initial values of both TGF-β1 and VEGF were increased, as indicated by the color code (from ×1 to ×10). Solid lines indicate the average simulation of 30 parameter sets, and shade areas indicate SD. E) Simulation of THBS1 and FMOD using developed mathematical model. The initial value of TGF-β1 for THBS1 or VEGF for FMOD were increased, as indicated by the color code (From ×1 to ×10). Solid lines indicate the average simulation of 30 parameter sets, and shade areas indicate SD. F) Sensitivity analysis of TGF-β1-induced THBS1. Negative coefficients (blue) indicate that the quantity of the response metric decreases as species increase, while positive coefficients (red) indicate that the quantity of the metric increases.

We constructed a comprehensive ODE model of the TGF-β and VEGF receptor signaling network to identify the mechanism of skin aging that is transiently regulated by THBS1 and FMOD (Figure S5B). The integrated model included latent TGF-β1 activation by THBS1^28^, inhibition by FMOD^29–31^, SMAD-AP1 complex formation^42^, a negative feedback by SMAD7^43^, positive feedback of THBS1^28, 44–46^, VEGFR and Raf-ERK cascade^24, 47^, and PI3K-Akt crosstalk between the TGF-β and VEGF pathways (identified in “THBS1 and FMOD expression is controlled by the TGF-β1 signaling pathway”). The model was constructed using parameters estimated from time-course data (i.e., 0 min, 15 min, 30 min, 60 min, 120 min, 8 h, 24 h, and 48 h) of protein phosphorylation or expression (i.e., phosphorylated SMAD3, c-Fos, THBS1, phosphorylated Akt, and FMOD) in HFF-1 cells stimulated with or without TGF-β1 (Figure 4D, image: S6A). We obtained 30 well-fitting parameter sets (Figure S6B) that reproduced the experimental data (see details in the Methods section). The resulting model had 79 rate equations, 83 species, and 194 parameters. To validate the model and assess the reproducibility of the model dynamics, we used phosphor-SMAD2 data, which was not used as training data (Figure S6D). These results indicated that the generated model could successfully reproduce most of the experimental results for HFF-1 cells.

### Late inhibition of PI3K-Akt is crucial for FMOD downregulation by TGF-β1

Since TGF-β1 and VEGF expression is increased during cellular senescence^48, 49^ as well as in HFF-1 cells with higher PDL (Figure 2A, S2I), numerical simulations were performed with the generated comprehensive ODE model to identify the key regulators of senescence. Accordingly, the initial concentrations of TGF-β1 and VEGF were increased to represent the state of senescence (Fig 4E). An increase in input gradually increased the levels of phosphorylated SMAD3 and activated Ras (Ras-GTP). Interestingly, as senescence progressed, the maximum stable state of THBS1 increased around 24 h, but that of FMOD decreased compared to the control around 24 h later. We also found that, while the basal condition (blue line) induced transient activation of phosphorylated Akt, reactions with elevated initial values induced sustained Akt activation. This is likely because Akt activation changes from transient to sustained activation as cellular senescence progresses, and the sustained activation suppresses FMOD expression in the later stages of dynamics.

To verify the simulation, FMOD recovery in TGF-β1-treated HFF-1 cells was observed by adding LY294002 at specific time points to suppress Akt phosphorylation (Figure 5A). While FMOD expression was significantly reduced by the TGF-β1 treatment alone, it was restored by co-treatment with LY294002. FMOD recovered to the same level observed at 0 h when the inhibitor was initially added up to 24 h, whereas no effect was observed after 32 h. This indicates that sustained Akt activity is important for the negative regulation of FMOD by TGF-β1. The same trend was also observed with Akt inhibitor VIII (Figure S6F). Thus, from the model simulations and their validation, the dynamics of Akt activity were found to be critical for the downregulation of FMOD expression in the progression of cellular senescence.

**Figure 5.**
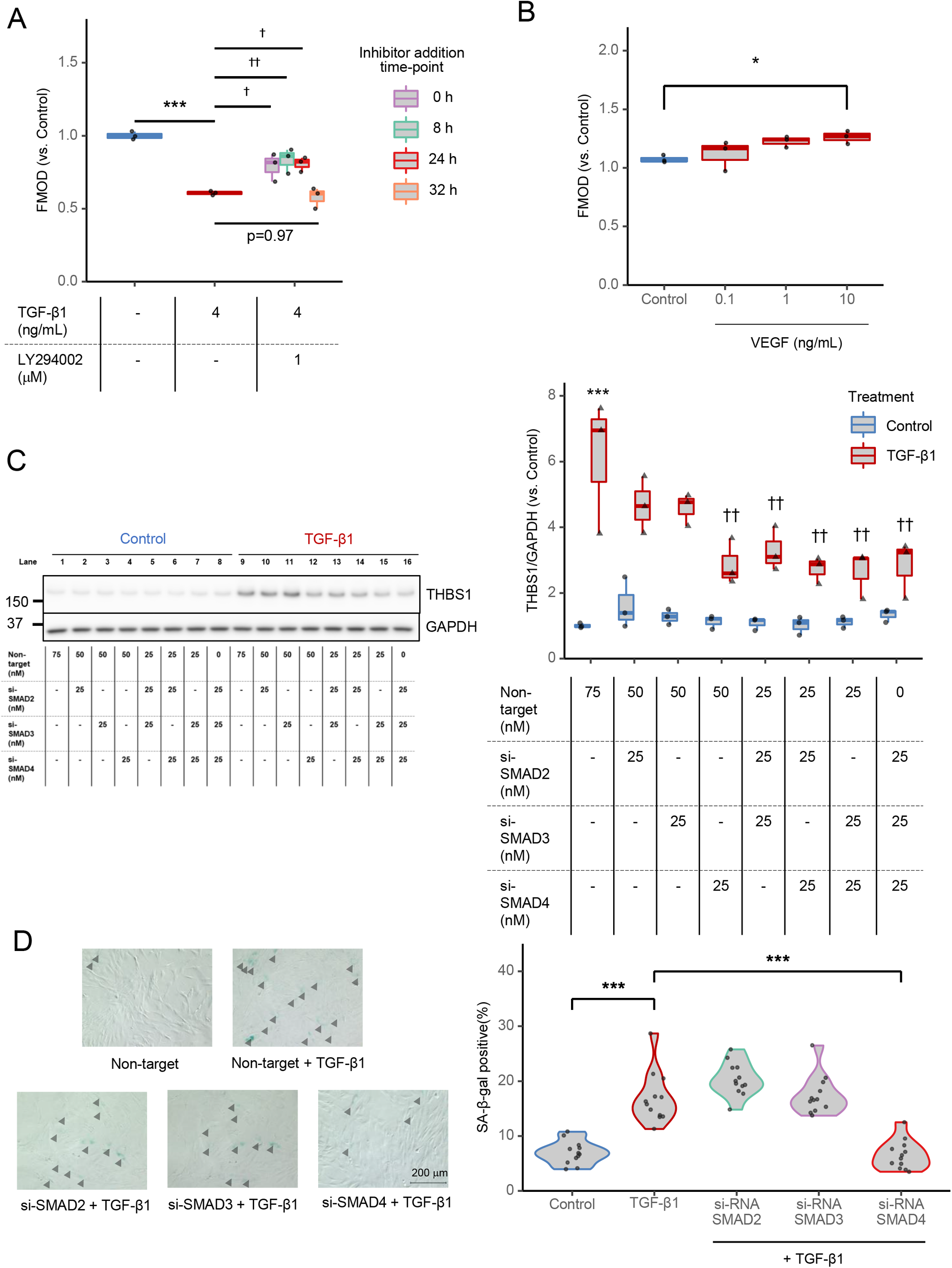
Inhibiting SMAD complex formation as a potential target for downregulating THBS1 expression. A) Quantification of FMOD in LY294002-treated HFF-1 cells by ELISA. HFF-1 cells were initially treated with TGF-β1 and LY294002 was added at certain time points (0 h, purple; 8 h, green; 24 h, bright red; 32 h, orange) and supernatants were collected at 48 h. The relative expression to the Control (DMEM supplemented with 2% FBS) is shown. N = 3, *** p < 0.01 (vs. Control, Student’s *t*-test), † p < 0.05, †† p < 0.01 (vs. TGF-β1, Dunnett’s test). B) Experimental validation of simulated FMOD expression in HFF-1 cells treated with VEGF165 by ELISA. HFF-1 cells were treated with VEGF165 for 48 h and the supernatants were analyzed. N = 3, * p < 0.05 (Dunnett’s test). C) Validation of the sensitivity analysis by western blot analysis of THBS1 following siRNA KD of SMADs in HFF-1 cells. Lysates were collected 48 h after pretreatment with each SMAD siRNA alone or in combination for 24 h prior to stimulation with or without TGF-β1 (4 ng/mL). (Left panel) Representative image. (Right panel) Quantification of THBS1 with SMAD KD. N = 3, *** p < 0.001 (vs. Non-target Control treatment, Student’s *t*-test), †† p < 0.01 (vs. Non-target TGF-β1 treatment, Dunnett’s test). D) Effect of siRNA KD of SMADs on SA-β-gal activity. HFF-1 cells were pretreated with each siRNA (25 nM) and stimulated with TGF-β1 (4 ng/mL). (Left panel) Representative images. SA-β-gal-positive cells are indicated with the arrowhead (black). Scale bars, 200 μm. (Right panel) Quantification of SA-β-gal: SA-β-gal-positive rate (%) = SA-β-gal-positive cells / total number of cells × 100, N = 3 (4 points/well), *** p < 0.001 (Tukey’s multiple comparisons).

### THBS1 is a sensitive factor, FMOD is a robust factor, and SMAD2/3/4 formation is effective in suppressing THBS1 expression

We aimed to identify the target molecule regulating the rate of cellular senescence. Our findings thus far point to two possible targets: THBS1 can be downregulated or FMOD can be upregulated. We determined which target is suitable for manipulation by simulating THBS1 and FMOD expression as model outputs and increasing the input of TGF-β1 and VEGF, respectively (Figure 4F). Interestingly, the simulation showed that THBS1 expression was sensitive to TGF-β1 input, whereas FMOD was hardly changed by VEGF input. This prediction for VEGF was confirmed in an *in vitro* experiment with HFF-1 cells (Figure 5B). Indeed, the expression of FMOD was significantly increased with increasing VEGF concentrations, but the rate of increase was much lower than that for THBS1 induced by TGF-β1 (Figure 3A). These results suggest that THBS1 is a sensitive factor and FMOD is a robust factor in cell senescence.

To further identify the molecular factors regulating THBS1 activity, we performed a sensitivity analysis^50^ to determine the bottleneck of the TGF-β signaling network with THBS1 as output (Figure 4G). A reaction involving SMAD2–SMAD3–SMAD4 complex formation (V13) was shown to be more sensitive than reactions involving SMAD2 and SMAD3, which suggests that inhibition of the SMAD4 reaction can effectively suppress THBS1 production. This simulation was validated by monitoring TGF-β1-induced THBS1 expression after siRNA KD of SMAD2, SMAD3, and SMAD4 in HFF-1 cells (Figure 5C). As suggested by the model, SMAD4 KD had the greatest effect on THBS1 expression compared to that of SMAD2 and SMAD3. Combinations of SMAD2–SMAD4, SMAD3–SMAD4, or SMAD2–SMAD3–SMAD4 KDs did not show synergistic or additive effects compared to SMAD4 KD alone. After KD of each SMAD, we found that SMAD4 KD significantly suppressed the induction of TGF-β1-induced SA-β-gal positivity compared to that by SMAD2 and SMAD3 KD (Figure 5D). These results suggest that SMAD4 is important for THBS1 induction and is a suitable target for controlling cell senescence.

## DISCUSSION

This study used multi-omics and mechanistic approaches to investigate the regulatory network in skin aging. Our integrative analysis of RNA-seq, H3K27Ac ChIP-seq, and ATAC-seq using replication-stressed HFF-1 cells and public *in vitro* and *in vivo* data of human skin fibroblasts identified TGF-β1 as a critical upstream regulator. Since public RNA-seq data on internal and external skin aging support the enrichment of the TGF-β signaling pathway, we consider the HFF-1 model to be suitable for studying skin aging. A previous study showed that inflammation and fibrosis define senescence *in vivo*^51^. NF-κB signaling, which regulates inflammation, has been associated with cellular senescence and aging^21^. In contrast, the TGF-β signaling pathway, which regulates fibrosis, has not been extensively studied in senescence. Nevertheless, we know that senescent cells can induce senescence in surrounding cells via TGF-β1^49, 52^; the transcription of p21, a critical inducer of cellular senescence, can be activated by the TGF-β signaling pathway^53^; and a transcriptome analysis of aging mouse skin implicated the TGF-β pathway as a regulator^54^. These results support TGF-β1 as a key upstream regulator of human skin aging. In addition to TGF-β1, our analysis identified an age-related enrichment of estrogen receptor 2 (ESR2). Knockout of the human *ESR2* gene reduced the expression of human *THBS1* mRNA^55^ and its ligand 17β-estradiol induced *THBS1* mRNA expression^56^. 17β-estradiol decreased *Fmod* mRNA expression in the frontal cortex of rats^57^. These results suggest that ESR2 may also be involved in the regulation of THBS1 and FMOD along with TGF-β1 in dermal senescence.

Our study showed that the expression of THBS1 increased with the senescence of dermal fibroblasts, aging of dermal tissue, as well as in HGPS patients, and THBS1 itself was found to induce senescence in dermal fibroblasts. In earlier reports, analysis of human dermal fibroblast mRNA suggested that *THBS1* expression increased with age^58^. Among 998 proteins that showed an age-dependent secretion pattern, THBS1 was upregulated with skin aging^59^. Earlier reports implicated THBS1 as a biomarker of skin aging but could not directly demonstrate its function. Collectively, the current and previous findings suggest that THBS1 expression is a driver and universal phenotype in skin aging. In addition, THBS1 was reported to promote senescence in endothelial cells^60, 61^. Given the strong relationship between THBS1 and age-associated diseases^62^, our current findings on THBS1 have potential applications beyond skin aging. In contrast, the expression of FMOD seems to be heterogeneous among reports of skin aging. We found that the expression of FMOD decreased with the senescence of dermal fibroblasts. Proteomic analysis of human skin punch biopsies showed that FMOD protein expression decreased with age^63^. In contrast, FMOD expression was upregulated in the public RNA-seq dataset with increasing age and senescence (Figure S2D, S2E). An earlier analysis of skin-fibroblast mRNA suggested that FMOD increased with age^58^. These inconsistent reports on the regulation of FMOD may be due to differences between proteomic and transcriptomic data in aging, suggesting that their expressions are poorly correlated and that mRNA profiling alone does not provide the complete picture^64^.

We elucidated the regulatory network of THBS1 and FMOD in dermal fibroblasts using small molecule inhibitors and siRNA KD. In earlier studies, THBS1 was regulated through the SMAD3 binding site of the THBS1 promoter^46^; c-Jun, but not c-Fos, was involved in AP-1 activity at the AP-1 binding site of the THBS1 promoter in human hepatocarcinoma cell lines^65^. Our findings suggest that both SMAD activation and c-Fos/c-Jun binding to DNA are required to regulate THBS1 expression in dermal fibroblasts. As various TFs (e.g., NF-κB, USF, E2F1, AP-1, EGR1, and SP1) have been reported to bind to the promoter of THBS1^66^, our results suggest that, while SMAD activation is essential for THBS1 expression, the TFs associated with THBS1 may vary among cell lines. Meanwhile, FMOD expression in dermal fibroblasts was found to be regulated by the VEGF–Raf–ERK pathway. Previous reports indicated that FMOD was regulated by the Wnt/β-catenin pathway in human breast cancer cell lines^67^, MAPK/AP-1 pathway in human pancreatic stellate cells^68^, and TGF-β2 in rat pericytes^69^. These results suggest that the regulation of FMOD expression is tissue and species specific.

From the bifurcation analysis, THBS1 was identified as the main controller of the TGF-β1, THBS1, and FMOD network. Because this network was described in terms of bistable states, i.e., senescent and non-senescent, it becomes difficult to return to the original state once a transition occurs. Thus, this model is also representative of the robustness and irreversibility of the senescent state. Earlier studies identified a bistable switch that regulates TGF-β1 activation in liver fibrosis^70^ and asthmatic airways^71^. Our analysis of TGF-β1 regulation indicated that, in addition to these diseases, the bifurcation analysis may be useful in qualitatively capturing other phenomena of senescence.

Several mathematical models for the TGF-β signaling pathway have been reported^72–74^. Nevertheless, our model is invaluable because it reflects unbiased data-driven insights into the gene regulatory network of skin aging. The reported models do not include activation of latent TGF-β by THBS1 or binding of FMOD to TGF-β and are not specific to skin aging. We integrated our TGF-β and VEGF signaling pathways to comprehensively elucidate the regulatory network between THBS1 and FMOD. Our mathematical model of skin aging showed that THBS1 responded sensitively while FMOD was robustly regulated with any input. These results suggest that the downregulation of THBS1 is a more promising drug target than the upregulation of FMOD. The sensitivity analysis further confirmed that the model-predicted sensitive response, i.e., the complex formation by SMAD4, was conserved across 30 independent parameter sets, suggesting that parameter identifiability does not affect the uncertainty of model outputs. The approach of targeting THBS1 by inhibiting SMAD4 complex formation will open new avenues for skin aging research.

To the best of our knowledge, this is the first study to integrate multi-omics analysis, *in vitro* experimental data, and mechanistic modeling to elucidate the THBS1 and FMOD regulatory network of skin aging. There is a growing demand for representative simulations of real-world disorders and diseases, with our work representing a considerable advance in the modeling of skin aging.

### Limitations of the study

The mechanisms regulating FMOD expression in dermal fibroblasts remain largely unclear. Therefore, we used an anonymous transcription factor for FMOD in our mathematical model. Further research is needed to elucidate the factors regulating FMOD in skin aging, especially as the regulation of FMOD may vary in different cells and tissues or environments.

## Supporting information

Key_Resource_Table

## ACKNOWLEDGEMENTS

The authors would like to thank all the members of the Laboratory of Cell Systems, Osaka University, for their kind advice with this study. We especially thank Dr. S. Tabata for his fruitful discussions about the project. We are also grateful to Mr. J.N. Wibisana for his guidance in the bioinformatics analysis, and Dr. H. Imoto and Mr. K. Murakami for their help in the mathematical analysis. M.O. received funding for this study from the JSPS KAKENHI (18H04031), JST CREST Program (JPMJCR21N3), and Uehara Memorial Foundation. K.I. received funding for this study from the JSPS KAKENHI (20K14361) and JST Moonshot (J225201004).

## AUTHOR CONTRIBUTIONS

M.H. and M.O. planned the experimental and study design. M.H. preformed all data curation, preparation of resources for sequence analysis, and data analysis. K.I. performed the bifurcation analysis. M.H. and M.O. wrote and reviewed the manuscript. M.H. and M.O. interpreted the results of the analysis. M.O. provided overall supervision.

## STAR METHODS

### RESOURCE AVAILABILITY

#### Lead Contact

Requests for raw data and code should be directed to and will be fulfilled by the Lead Contact, Mariko Okada.

#### Materials Availability

This study did not generate new unique reagents.

#### Data and Code Availability

All sequence data have been deposited in the DNA Data Bank of Japan (DDBJ) under DRA016119. The code for the bioinformatics analysis and mathematical modeling is available from GitHub (https://github.com/okadalabipr/Haga2023). Upon reasonable request, all other data will be made available.

### EXPERIMENTAL MODEL AND SUBJECT DETAILS

#### Cell lines

HFF-1 and BJ cells were purchased from the American Type Culture Collection (ATCC). HFF-1 and BJ cells were maintained using Dulbecco’s modified Eagle’s medium (DMEM, ATCC) supplemented with 10% fetal bovine serum (FBS, Corning) and 1% antibiotic–antimycotic (Thermo Fisher Scientific). Unless otherwise noted, the PDL of HFF-1 cells in the following experiments was kept at approximately 24, and that for BJ cells was approximately 36, which correspond with young cells. In the experiments, the cells were serum starved with 0.1% FBS and tested under uniform 2% FBS conditions during stimulation. All cell lines were maintained at 37 °C in a humidified atmosphere at 5% CO_2_.

#### Human dermal tissue

Human full-thickness dermal tissues were purchased from Biopredic International via KAC and stored frozen at –20 °C. Skin samples were anonymized, and all participants provided informed consent (see Table S2 for donor list). This study using human tissue samples was approved by the Ethics Committee of the Institute for Protein Research, Osaka University (clearance no. 2021-2).

### METHODOLOGICAL DETAILS

#### Induction of replication stress in dermal fibroblasts

Replication stress was induced by passage culturing of HFF-1 and BJ cells. Briefly, cells were seeded in a cell culture flask and grown on DMEM supplemented with 10% FBS until reaching a sub-confluent condition. The experiment was performed in three separate flasks. After removing the medium, washing with PBS, and detaching using trypsin/EDTA (ATCC), cells were frozen in BAMBANKER^®^ solution (NIPPON Genetics) at –80 °C in a BICELL container (Nihon Freezer) and then stored under liquid nitrogen. The PDL for each collection was calculated as follows: n = 3.32 (log A – log B) + X (n: final PDL of cell line, A: yield of harvested cells, B: count of seeded cells, X: initial PDL of the seeded cell population).

#### Sample preparation for RNA-seq and genomic alignment

HFF-1 cells (PDL 24, PDL 36, and PDL 47) were used to prepare RNA-seq libraries. Briefly, each PDL was seeded in 6-well plates at 200,000 cells/well and total RNA (three wells per sample) was collected after 48 h using a NucleoSpin RNA kit (Macherey-Nagel GmbH & Co.). The quality of the total RNA was evaluated using a 2100 Bioanalyzer (Agilent) and RNA samples with RNA integrity > 9.0 were used for library preparation. cDNA libraries were prepared using a NEBNext^®^ Poly(A) mRNA Magnetic Isolation Module (New England Biolabs) for PolyA selection and NEBNext^®^ Ultra™ ll Directional RNA Library Prep Kit (New England Biolabs). Samples were prepared according to the manufacturer’s protocol. The RNA-seq data were generated as paired-end 150 base reads on a NovaSeq 6000 (Illumina). The expression of specific genes was validated by qRT-PCR.

All RNA-seq data, including public RNA-seq data, were trimmed using Trim Galore! version

0.6.6 and aligned to human reference genome GRCh38 using hisat2 version 2.2.1. Mapped reads were extracted using samtools version 1.9 and a read count matrix was created using gene annotation (GRCh38.p13) with Subread version 2.0.1 for downstream analysis.

#### Sample preparation for ChIP-seq and genomic alignment

HFF-1 cells (PDL 24, PDL 36, and PDL 47) were used to prepare ChIP-seq libraries. Briefly, cells for each PDL were seeded in a 145 mm dish at 4,000,000 cells per dish. After 48 h of incubation, the cells were collected using a SimpleChIP Enzymatic Chromatin IP kit (Cell Signaling Technology) to obtain sheared chromatin. Briefly, two dishes per sample were fixed with fresh 1% formaldehyde (Thermo Fisher Scientific) for 5 min. The cells were subjected to a micrococcal nuclease treatment at 37 °C for 20 min and M220 Focused-ultrasonicator (Covaris) for 10 min to obtain chromatin in 150–900 bp DNA/protein fragments. Next, H3K27Ac antibody (Abcam) and an iDeal ChIP-seq kit for Histones (Diagenode) were used to obtain ChIP-DNA. Using an SX-8G (Diagenode) platform, 2 μg anti-H3K27Ac antibody was incubated with 10 μL DiaMag ProteinA-coated magnetic beads for 3 h and an IP reaction was performed for 12 h. Decrosslinking was performed using NaCl and Proteinase K (Thermo Fisher Scientific) at 60 °C for 4 h. The purification of ChIP-DNA was conducted using a MinElute^®^ PCR Purification Kit (Qiagen). The ChIP-seq libraries were prepared using a NEBNext^®^ Ultra II DNA Library Prep Kit for Illumina (New England Biolabs). All samples were prepared according to the manufacturer’s protocol. ChIP-seq data were generated as paired-end 150 base reads on the NovaSeq 6000 (Illumina).

All ChIP-seq data were analyzed using the nfcore/chipseq pipeline version 1.2.2 with “nextflow run nf-core/chipseq -r 1.2.2 -profile singularity –input samplesheet_ChIP.csv --genome GRCh38 --save_reference --max_cpus 16 --max_memory 256.GB.” Briefly, reads were trimmed using Trim Galore! and aligned against the GRCh38 reference genome using BWA. MACS2 was used for broadPeak calling. Consensus peak sets across all samples were created using BEDTools version 2.30.0. The counts for consensus peaks were generated using FeatureCounts and differential chromatin accessibility was analyzed using DESeq2 version 1.36.0. For more details, see the nf-core/chipseq pipeline (https://nf-co.re/chipseq/1.2.2).

#### Sample preparation for ATAC-seq and genomic alignment

HFF-1 cells (PDL 24, PDL 36, and PDL 47) were used to prepare ATAC-seq libraries. Cells for each PDL were seeded in 6-well plates at 200,000 cells/well. After 48 h of incubation, the cells were washed with PBS and detached using trypsin/EDTA. Using 200,000 cells of each PDL, ATAC-seq libraries were constructed using an ATAC-Seq Kit (Active Motif) according to the manufacturer’s protocol. The ATAC-seq data were generated as paired-end 150 base reads on the NovaSeq 6000 (Illumina).

All ATAC-seq data were analyzed using the nfcore/atacseq pipeline version 1.2.1 with default parameters using the “nextflow run nf-core/atacseq -r 1.2.1 -profile singularity –input samplesheet_ATAC.csv --genome GRCh38 --save_reference --max_cpus 16 --max_memory 256.GB” arguments. The downstream procedure is similar to that for the ChIP-seq analysis (for more detail, see the nf-core/atacseq pipeline [https://nf-co.re/atacseq/1.2.1]).

#### Transcription factor enrichment

The activity of the top 20 TFs was determined for the DEGs according to the FC value for RNA-seq data between PDLs and normalized transcripts per million (TPM) value for RNA-seq data using DoRothEA version 1.8.0. First, to identify the DEGs in replication-stress induced HFF-1 cells, the FC and adj-p values were calculated for PDL 24 vs. PDL 36, PDL 24 vs. PDL 36, and PDL 24 vs. PDL 36 (DEGs: |FC| > 1.2, adj-p < 0.05) using DESeq2. DEGs and FC values were used as input to identify the top 20 TFs regulating the DEGs using DoRothEA (confidence level A, B, and C). The top 20 TFs were identified by extracting the top 10 TFs for PDL 24 vs. PDL 36, PDL 24 vs. PDL 36, and PDL 24 vs. PDL 36, excluding duplicate TFs. Finally, a normalized enrichment score (based on DoRothEA) was calculated for the top 20 TFs using normalized TPM values for all genes greater than 5.

#### Motif analysis

Motif analysis was performed using HOMER version 4.11. Briefly, we first calculated the gain peak region of ATAC-seq from the counts at the consensus peak using DESeq2. The region with Log_2_FC > 0 and adj-p < 0.05 was defined as the gain peak that significantly increases with increasing PDL. Next, the gained peaks for PDL 24 vs. PDL 36, PDL 24 vs. PDL 36, and PDL 24 vs. PDL 36 were concatenated using the “cat” command, sorted and merged using “sortBed” and “mergeBed,” respectively, in Bedtools version 2.30.0. The concatenated gained peak bed file was analyzed with “findMotifsGenome.pl hg38 -size given -p 8” in HOMER to output log(adj-p value) and percent of target sequences with motif from knownResults.html.

#### Gene annotation of peaks for ATAC-seq and ChIP-seq and intersection with DEGs

Peak annotation for ATAC-seq and ChIP-seq was performed using “annotatePeak” in the R package ChIPseeker version 1.32.1. First, the region with |Log2FC| > 0 and adj-p < 0.05 was defined as the differential peak that changes significantly for PDL 24 vs. PDL 36, PDL 24 vs. PDL 36, and PDL 24 vs. PDL 36 using DESeq2. Next, differential peaks were concatenated using the “cat” command, sorted and merged using “sortBed” and “mergeBed” in Bedtools. The differential peaks were annotated as the nearest neighboring gene with the closest distance from the peak to the transcription start site (TSS). The TSS region occurred from –3kb to +3kb. The annotation package for hg38 (TxDb.Hsapiens.UCSC.hg38.knownGene)^87^ was used as the TxDb object. Peak annotation was conducted with the “tssRegion = c (−3000, 3000), TxDb = TxDb.Hsapiens.UCSC.hg38.knownGene, annoDb = ‘org.Hs.eg.db’” option.

DEGs from the RNA-seq data were identified for PDL 24 vs. PDL 36, PDL 24 vs. PDL 36, and PDL 24 vs. PDL 36 using DESeq2 (DEGs: |FC| > 1.2, adj-p < 0.05). Venn diagrams were created using DEGs and annotated genes from differential peaks in both the ATAC-seq and ChIP-seq data.

#### Ingenuity Pathway Analysis (IPA)

IPA was performed for upstream analysis using IPA tool version 81348237 software. The gene set was mapped to Ingenuity Knowledge Base using “core analysis.” Molecule type, genes, RNAs, and proteins were used for the upstream analysis. A right-tailed Fisher’s exact test was used to calculate a p-value of overlap determining the probability that the association between the gene set and the upstream regulators is explained by chance alone. The Benjamini–Hochberg (BH) method for multiple testing was used to calculate the adjusted p-value.

#### Analysis of public *in vivo* and *in vitro* RNA-seq data

All public RNA-seq data used the same genomic alignment and read count matrix pipeline as described in the “Sample preparation for RNA-seq and genomic alignment”.

*In vivo* RNA-seq data (GSE113957) of human arm skin was analyzed after filtering outliers and clustering. Since multiple ethnic and tissue sources were included, data for 55 samples were manually selected for skin fibroblasts derived from Caucasian individuals (11–71 years of age, see donor list in Table S1). The samples were manually divided into three clusters according to age (young: 10s to 20s; middle: 30s to 50s; aged: 60s to 70s). For each cluster, a robust principal component analysis (PCA) was performed using “rrcov” version 1.5.2 to detect outliers, resulting in a final dataset with 45 samples. Spearman’s correlation coefficient was calculated to evaluate the distance between each sample using normalized TPM values: distance =1 – Spearman’s correlation coefficient. Using the distance calculated, samples were clustered using the “hclust” function in R with “distance, method = ‘ward.D2’” (Figure S2A, S2B). Finally, DEGs were identified by comparing each cluster using DESeq2 (DEGs: |FC| > 1.5, adj-p < 0.05). The BH method was used to calculate the adjusted p-value.

*In vitro* RNA-seq data (GSE63577) for the human dermal fibroblast HFF-1 cell line was compared for each PDL (PDL 16, PDL 26, PDL 46, PDL 64, and PDL 74). DEGs were identified for PDL 16 vs. PDL 26, PDL 16 vs. PDL 46, PDL 16 vs. PDL 64, PDL 16 vs. PDL 74, PDL 26 vs. PDL 46, PDL 26 vs. PDL 64, PDL 26 vs. PDL 74, PDL 46 vs. PDL 64, PDL 46 vs. PDL 74, and PDL 64 vs. PDL 74 using DESeq2 (DEGs: |FC| > 2.0, adj-p < 0.01) and intersected. The BH method was used to calculate the adjusted p-value.

RNA-seq data for skin fibroblasts from healthy subjects (age: 1–9 years, N = 12) and Hutchinson–Gilford progeria syndrome (HGPS) patients (age: 2–8 years, N = 10) were downloaded from GSE113957 (see donor list in Table 1).

#### Kyoto Encyclopedia of Genes and Genomes (KEGG) analysis of public RNA-seq

KEGG enrichment was compared using the “comparecluster” function in the R package clusterProfiler version 4.4.4 with “fun” = enrichKEGG, “organism” = hsa, “keyType” = kegg, “pAdjustMethod” = BH, “minGSSize” = 10, “maxGSSize” = 500, “pvalueCutoff” = 0.05, “qvalueCutoff” = 0.05, and “use_internal_data” = “FALSE”. The BH method was used to calculate the adjusted p-value.

#### Correlation analysis of TGF-β pathway-enriched genes using public RNA-seq data

Spearman’s correlation between age and gene expression was calculated for the *in vivo* data or PDL and gene expression for the *in vitro* data using “cor” (method = “spearman”) in R.

#### Extraction of human skin tissue protein

Skin tissue proteins were extracted after dividing full thickness skin into dermis and epidermis as previously reported^88^. Briefly, a biopsy punch (tip diameter: 3 mm ø) was used to collect a full-layer skin sample, from which subcutaneous tissue was physically removed. After washing in 70% ethanol and HBSS (Thermo Fisher Scientific), samples were treated with 25 UI/mL Dispase II (Roche) HBSS solution for 15 h at 4 °C. Enzymatic digestion was inactivated with HBSS supplemented with 10% FBS and the dermis and epidermis were separated with tweezers. Each dermis and epidermis were separately homogenized in RIPA buffer (Thermo Fisher Scientific) supplemented with Halt™ Protease and Phosphatase Inhibitor (Thermo Fisher Scientific) for 10 min using a Powermasher II (Nippi) and centrifuged (13 000 rpm, 20 min, 4 °C) to generate protein extracts for western blot analysis.

#### Time-course cell proliferation analysis (trypan blue)

HFF-1 cells were seeded in 6-well plates with DMEM at 200,000 cells/well. After serum starvation for 16 h, cells were treated with either control (DMEM with vehicle supplemented with 2% FBS) or 4 ng/mL TGF-β1 and collected at 0 h, 8 h, 12 h, 24 h, and 48 h. Reconstitution buffer (0.1% bovine serum albumin [BSA] in 4 mM HCl PBS; R&D Systems) was used as vehicle and to dissolve TGF-β1. At each time point, cells were washed with PBS and detached using trypsin/EDTA to count cells positive for trypan blue (Thermo Fisher Scientific) using a cell counter (WakenBtech) according to the manufacturer’s protocol.

#### Cell viability analysis (WST-8)

HFF-1 cells were seeded in 96-well plates with DMEM at 10,000 cells/well. After serum starvation for 16 h, cells were treated with either control (DMEM with vehicle supplemented with 2% FBS), 4 ng/mL TGF-β1, or 1 μg/mL THBS1. After 24 h, 10 μL Cell Counting Kit-8 solution (DOJINDO) was added, incubated for 1 h, and cell viability was calculated at an absorbance wavelength of 450 nm using a Multiskan FC system (Thermo Fisher Scientific).

#### 5-Bromo-2-deoxyuridine (BrdU) incorporation assay

The inhibition of DNA synthesis was measured using a CycLex Cellular BrdU ELISA Kit Ver.2 (Medical & Biological Laboratories) according to the manufacturer’s instructions. Briefly, HFF-1 cells were seeded in 96-well plates with DMEM at 10,000 cells/well. After 16 h of serum starvation, cells were treated with either control (DMEM with vehicle supplemented with 2% FBS), 4 ng/mL TGF-β1, or 1 μg/mL THBS1. After 8 h, the cells were incubated with anti-BrdU antibody and substrate for 16 h, and absorbance was measured at 450 nm with 540 nm as a reference using the Multiskan FC system (Thermo Fisher Scientific).

#### SA-β-gal staining

The rate of positive SA-β-gal staining was calculated using a SA-β-gal Detection Kit (BioVision) according to the manufacturer’s protocol. Briefly, cells were seeded in 12-well plates at 100,000 cells/well, treated with each stimulant, and fixed for 10 min with fixative solution after 48 h. The cells were stained with Staining Solution Mix at 37 °C overnight. After washing with PBS, cells were treated with Hoechst^®^ 33342 (DOJINDO) for 10 min and washed with PBS before observation (BZ-9000, KEYENCE). SA-β-gal-positive rates were calculated as follows: SA-β-gal-positive rate (%) = number of SA-β-gal-positive cells per image / total number of cells per image × 100. For each condition, four images were analyzed per well (N = 3). The number of SA-β-gal-positive cells was counted manually and the total number of cells per image was counted as Hoechst^®^ 33342-positive cells using the ImageJ Fiji package (The National Institutes of Health).

#### siRNA transfection

Cells were transfected with siRNA oligomer (Dharmacon/Horizon Discovery) mixed with Lipofectamine RNAiMAX reagent (Thermo Fisher Scientific) in Opti-MEM™ I (Thermo Fisher Scientific) according to the manufacturer’s instructions. Briefly, cells were seeded in 6-well plates with DMEM at 200,000 cells/well. After transfection for 24 h in serum-starved conditions, cells were treated with either control (DMEM with vehicle supplemented with 2% FBS) or 4 ng/mL TGF-β1 and the lysates were collected. The efficacy of SMAD2, SMAD3, and SMAD4 KD at 25 nM siRNA was verified in Fig. S4A. See Fig. S4B for KD efficacy of c-Fos and c-Jun at 50 nM siRNA.

#### Kinase inhibitor treatment for FMOD regulatory pathway

Cells were seeded in 96-well plates with DMEM at 10,000 cells/well. Cells were treated with either control (DMEM with 0.1% DMSO supplemented with 2% FBS) or each inhibitor and the supernatants were collected for FMOD ELISA. The following inhibitors were used: LY364947 (123-05981, Fujifilm), Akt inhibitor VIII (14870, Cayman Chemical), LY294002 (440202, Calbiochem), and T-5224 (S28966, Selleck). The following inhibitors were dispensed from a Tocriscreen Kinase Inhibitor Toolbox (3514, Tocris Bioscience): ZM306416HCl (2499), Ki8751 (2542), GW5074 (1381), U0126 (1144), and SB 431542 (1614).

#### Bifurcation model analysis

Based on the experimental expression of THBS1 and FMOD upon treatment with different concentrations of TGF-β1 (Figure 3A), we constructed a core transcription factor network consisting of TGF-β1, THBS1, and FMOD. A double positive feedback loop (between THBS1 and TGF-β1) and double negative feedback loop (between FMOD and TGF-β1) can be described by the following nonlinear ordinary differential equations:

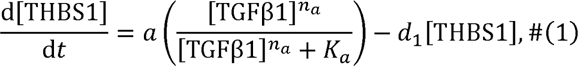

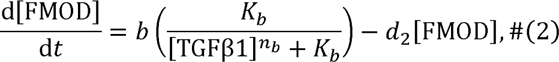

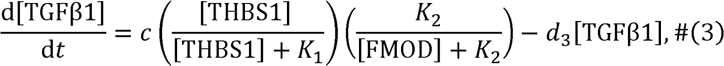

where *a*/*d*_1_, *b*/*d*_2_, and, *c*/*d*_3_ are maximal values of [THBS1], [FMO], and [TGFβ1], respectively; *K_a_*, *K_b_*, *K_1_*, and *K*_2_ are the half saturation constants; and *n_a_* and *n_b_* are Hill coefficients. Setting the left-hand sides of Eqs. (1)–(3) to zero, the steady state solutions can be obtained as follows:

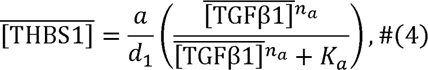

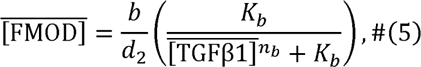

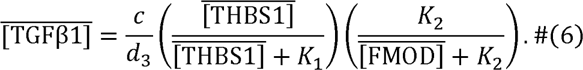

Substituting (4) and (5) into (6), 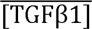 can be computed using Newton’s method and stability can be determined by the sign of the derivative. Note that 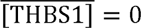, 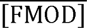 ≡ *b*/*d*_2_ and 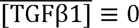 are stable steady state solutions for *K* >0 and *K*_2_ > 0. By fitting (4) and (5) to the experimental data (Figure 3A), the parameters were inferred using a nonlinear least-squares method with Gnuplot (version 5.4); *a*/*d*_1_ = 2.36, *b*/*d*_2_ = 0.33, *K_a_* = 0.016, *K_b_* = 0.002, *n_a_* = 1.6, and *n_b_* = 1.7 (Figure S5A).

#### Time-course datasets for mathematical model training

We used the time-course data for phosphorylated SMAD3, c-Fos, THBS1, phosphorylated Akt, and FMOD activity with or without 4 ng/mL TGF-β1 (eight time points up to 48 h) treatment in HFF-1 cells for data-fitting of the comprehensive model. Briefly, HFF-1 cells were seeded in 6-well plates at 200,000 cells/well and maintained in DMEM supplemented with 10% FBS. After serum starvation for 16 h, the cells were treated with either control (DMEM with vehicle supplemented with 2% FBS) or 4 ng/mL TGF-β1 and the lysate and supernatant were collected at 0 min, 15 min, 30 min, 60 min, 120 min, 8 h, 24 h, and 48 h. Reconstitution buffer (0.1% BSA in 4 mM HCl PBS, R&D Systems) was used as vehicle. The cells were lysed with RIPA buffer (Thermo Fisher Scientific) supplemented with Halt™ Protease and Phosphatase Inhibitor (Thermo Fisher Scientific) and used in the western blot analysis for analyzing anti-phosphorylated SMAD3 (Ser423/425; 9520, Cell Signaling Technology), anti-c-Fos (2524, Cell Signaling Technology), anti-THBS1 (37879, Cell Signaling Technology); and anti-phosphorylated Akt (Ser473; 9271, Cell Signaling Technology) expression. Supernatants were centrifuged (13 000 rpm, 15 min, 4 °C) to remove cell debris and used for the FMOD ELISA (ab275895, Abcam). In addition, anti-phosphorylated SMAD2 data (Ser465/467; 3108, Cell Signaling Technology) were obtained for model validation and not used in data-fitting. The data were normalized between the minimum (0) and maximum (1) values.

#### Model simulation and parameter estimation

We constructed the comprehensive mathematical model linking the TGF-β and VEGF signaling pathways (Figure S5B) by integrating two mathematical models using a Python framework for Modeling and Analysis of Signaling Systems (BioMASS)^47^.

Parameter estimation was conducted in two steps. In step one, parameters related to the TGF-β signaling network, modified based on the Lucarelli model^73^, were trained using normalized phosphorylated SMAD3, c-Fos, and THBS1 time-course expression. The resulting model had 27 rate equations, 22 species, and 42 parameters, of which 34 were to be estimated. By minimizing the objective function, i.e., the sum of residual squares between simulation and experimental values, 30 fitting parameter sets were obtained that reproduce the experimental results with/without TGF-β1 stimulation in HFF-1 cells. In step two, an additional 30 parameter sets for the rest of the model, including the Raf-ERK cascade and PI3K-Akt pathway, were trained using normalized phosphorylated SMAD3, c-Fos, THBS1, phosphorylated Akt, and FMOD time-course expression (Figure 4D). The best fitting parameters in the TGF-β signaling network were adapted from step one. The resulting integrated model had 79 rate equations, 83 species, and 194 parameters. An additional 30 fitting parameter sets were obtained that reproduce the experimental results of TGF-β1 stimulation in HFF-1 cells (parameter range: Figure S6B; objective function trace: Figure S6C).

For the TGF-β pathway, we developed an original model of TGF-βR activation, SMAD phosphorylation, and SMAD complex formation. This model activates the transcription of c-Fos at the same time and forms a logical AND gate with SMAD complexes to regulate THBS1 expression. THBS1 forms positive feedback and activates latent TGF-β1. We modified the Imoto model^47^ to describe the process from VEGFR receptor activation to ERK phosphorylation. ERK phosphorylation results in the activation of TFs for FMOD transcription. FMOD then forms a complex with activated TGF-β1 and inhibits its binding to TGF-βR. The TGF-β and VEGF signaling pathway models crosstalk via the PI3K-Akt pathway.

The mathematical formulas describing the TGF-β and integrated models, and TGF-β VEGF model can be found in Method S1 and Method S2, respectively. All related files to execute BioMASS can be found at https://github.com/okadalabipr/Haga2023. We described each biochemical reaction using ODEs. To train model parameters, we used time-series expression data for HFF-1 cells. As previously described^24, 47^, we minimized the sum of squared differences between the experimental observations and simulated values using the global parameter estimation method Differential Evolution^89^.

#### Initial value for mathematical model

Values measured by ELISA and RNA-seq were used to determine the initial values of the model.

The initial value for SMAD2, SMAD3, and SMAD4 were quantified using the respective ELISA kits (Figure S6E). Briefly, HFF-1 cells were seeded in 6-well plates with DMEM at 200,000 cells/well. Cell lysates were collected using the lysate buffer provided with the ELISA kits after serum starvation for 16 h, centrifuged (13 000 rpm, 15 min, 4 °C), and used for SMAD2 (ab260065, Abcam), SMAD3 (ab264624, Abcam), and SMAD4 ELISAs (ab253211, Abcam).

For other nonzero species, we used RNA-seq data obtained from non-stimulated HFF-1 cells to determine the initial value. Briefly, HFF-1 cells were seeded in 6-well plates with DMEM at 200,000 cells/well. Total RNA (three wells per sample) was collected after serum starvation for 16 h. RNA purification, sequencing library preparation, and downstream analysis was conducted as described in the “Sample preparation for RNA-seq and genomic alignment” section. Initial protein levels of the model species (total of 37 genes) were inferred from RNA-seq data as previously described^24, 47^, where the maximal transcription rate or translated protein level were estimated from the mRNA level. We estimated protein amounts from one or more genes belonging to the same gene family (isoforms) and employed weighting factors to convert the TPM value of the gene corresponding to the protein to the appropriate initial protein value. The TPM values are shown in Table S4.

#### Sensitivity analysis

The sensitivity coefficient Sy was calculated as previously described^24^ using the following equation:

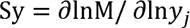

where M is the signaling metric, i.e., the integral expression level of THBS1 with TGF-β1 stimulation, and *y_j_* is each nonzero species in the mechanistic model^50^. The sensitivity coefficients were calculated by finite difference approximations with 1% changes in the biochemical reactions. To calculate the sensitivity coefficients, we used BioMASS with run_analysis = (target = “reaction”, metric = ‘‘integral’’, style = ‘‘heatmap’’).

#### Time-course PI3K-Akt inhibitor treatment

We validated the model results using HFF-1 cell experiments with additional inhibitors at each timepoint (four timepoints, up to 48 h). Briefly, HFF-1 cells were seeded in 96-well plates with DMEM at 10,000 cells/well. The cells were treated with either control (DMEM with 0.1% DMSO supplemented with 2% FBS) or 4 ng/mL TGF-β1. Inhibitors for TGF-β1-treated samples were added at 0 h, 8 h, 24 h, and 32 h. Supernatants were collected at 48 h for quantification. Reconstitution buffer (0.1% BSA in 4 mM HCl PBS, R&D Systems) was used as vehicle. Supernatants were centrifuged (13 000 rpm, 15 min, 4 °C) to remove cell debris and used for FMOD ELISA (ab275895, Abcam).

#### Western blot analysis

All immunoblots are representative images of three biological replicates. Attached cells were washed with PBS and lysed in RIPA buffer (Thermo Fisher Scientific) supplemented with Halt™ Protease and Phosphatase Inhibitor (Thermo Fisher Scientific). Cell lysates were centrifuged (13000 rpm, 15 min, 4 °C) to unify protein concentrations between samples, and a BCA protein assay kit (Thermo Fisher Scientific) was used according to the manufacturer’s protocol. Proteins were separated by SDS-PAGE and transferred to nitrocellulose membranes using iBlot2 (Thermo Fisher Scientific). After blocking with EveryBlot blocking buffer (Bio-Rad), the following antibodies were used for blotting: anti-THBS1 (37879, Cell Signaling Technology), anti-FMOD (60108-1-Ig, ProteinTech), anti-LAMIN-B1 (12987-1-AP, ProteinTech), anti-p53 (2524, Cell Signaling Technology), anti-p21 (2946, Cell Signaling Technology), anti-c-Fos (2524, Cell Signaling Technology), anti-c-Jun (9165, Cell Signaling Technology), anti-SMAD2 (5339, Cell Signaling Technology), anti-phosphorylated SMAD2 (Ser465/467; 3108, Cell Signaling Technology), anti-SMAD3 (9523, Cell Signaling Technology), anti-phosphorylated SMAD3 (Ser423/425; 9520, Cell Signaling Technology), anti-SMAD4 (46535, Cell Signaling Technology), anti-phosphorylated Akt (Ser473; 9271, Cell Signaling Technology), anti-GAPDH (M171-3, Medical & Biological Laboratories), and Anti-GAPDH (10494-1-Ap, ProteinTech). Primary antibodies were reacted at 4 °C overnight, while secondary antibodies were reacted at room temperature for 1 h. For protein detection, Clarity Western ECL Substrate (Bio-Rad) or Clarity Max Western ECL Substrate (Bio-Rad) was used with an Amersham Imager 680 (GE Healthcare). Relative protein quantification was performed with ImageJ Fiji. Expression values (N = 3) were normalized using GAPDH as loading control and the ratio against the control was calculated. Molecular weights are indicated on the left of each image.

#### Enzyme-Linked Immunosorbent Assay (ELISA)

All ELISAs were conducted according to the manufacturer’s protocol. Briefly, supernatant samples were centrifuged (1 000 rpm, 4 min, 4 °C) to remove cell debris and used for measurement. Lysate samples were collected using lysis buffer included in the kits, cell lysates were centrifuged (13 000 rpm, 15 min, 4 °C) and used for each ELISA. Relative expression levels are displayed with the control (1) value. The following ELISA kits were used for quantification: THBS1 (DTSP10-1, R&D Systems), FMOD (ab275895, Abcam), TGF-β1 (DB100B, R&D Systems), VEGF (DVE00, R&D Systems), IL-6 (D6050, R&D Systems), IL-8 (ab214030, Abcam), SMAD2 (ab260065, Abcam), SMAD3 (ab264624, Abcam), and SMAD4 (ab253211, Abcam).

#### Quantitative RT-PCR (qRT-PCR) analysis

Total RNA from HFF-1 cells was prepared using the NucleoSpin RNA kit (Macherey-Nagel GmbH & Co.) and subjected to complementary DNA (cDNA) synthesis using ReverTra Ace^®^ qPCR RT Master Mix (Toyobo Life Science) as follows: 15 min at 37 °C, 5 min at 50 °C, and 5 min at 98 °C. Quantitative PCR using cDNA was conducted using a KOD SYBR qPCR kit (Toyobo Life Science) with a CFX96 Real-Time PCR System (Bio-Rad) according to the manufacturer’s protocol. PCR cycling conditions were as follows: 40 cycles of 10 s at 98 °C, 10 s at 60 °C, and 30 s at 68 °C. The primers used for qRT-PCR were as follows: THBS1 (5′-TCCCCATCCAAAGCGTCTTC-3′ and 5′-ACCACGTTGTTGTCAAGGGT-3′); FMOD (5′-GGACGTGGTCACTCTCTGAA-3′ and 5′-GGCTCGTAGGTCTCATACGG-3′); GAPDH (5′-GTCTCCTCTGACTTCAACAGCG-3′ and 5′-ACCACCCTGTTGCTGTAGCCAA-3′). Gene expression was quantified using the ΔΔCq method. Expression values (N = 3) were normalized using GAPDH and the ratio against the PDL 24 value was calculated.

### QUANTIFICATION AND STATISTICAL ANALYSIS

Statistical data are presented as the mean with standard deviation (SD) calculated using the “sd” function of R. For detection of upstream regulators, the right-tailed Fisher’s exact test was used in the IPA. Comparisons of more than two groups were made using a one-way Dunnett’s or Tukey’s multiple comparisons test. Comparisons of two groups were evaluated by Student’s *t*-test, Welch’s *t*-test, or Wilcoxon rank sum test. We considered p < 0.05 to be statistically significant. Unless otherwise noted, p-values were calculated using the multcomp package in R. The statical details of each experiment can be found in the figure legends.

## REFERENCES

1. Niccoli, T., and Partridge, L. (2012). Ageing as a risk factor for disease. Curr. Biol. 22, R741–R752. 10.1016/j.cub.2012.07.024.

2. Albert, A., Knoll, M.A., Conti, J.A., and Zbar, R.I.S. (2019). Non-melanoma skin cancers in the older patient. Curr. Oncol. Rep. 21, 79. 10.1007/s11912-019-0828-9.

3. Mosteller, R.D. (1987). Simplified calculation of body-surface area. N. Engl. J. Med. 317, 1098. 10.1056/NEJM198710223171717.

4. de Bengy, A.F., Lamartine, J., Sigaudo-Roussel, D., and Fromy, B. (2022). Newborn and elderly skin: Two fragile skins at higher risk of pressure injury. Biol. Rev. 97, 874–895. 10.1111/brv.12827.

5. Krutmann, J., Bouloc, A., Sore, G., Bernard, B.A., and Passeron, T. (2017). The skin aging exposome. J. Dermatol. Sci. 85, 152–161. 10.1016/j.jdermsci.2016.09.015.

6. Costello, L., Dicolandrea, T., Tasseff, R., Isfort, R., Bascom, C., von Zglinicki, T., and Przyborski, S. (2022). Tissue engineering strategies to bioengineer the ageing skin phenotype in vitro. Aging Cell 21, e13550. 10.1111/acel.13550.

7. Shin, S.H., Lee, Y.H., Rho, N.K., and Park, K.Y. (2023). Skin aging from mechanisms to interventions: Focusing on dermal aging. Front. Physiol. 14, 1–10. 10.3389/fphys.2023.1195272.

8. Low, E., Alimohammadiha, G., Smith, L.A., Costello, L.F., Przyborski, S.A., von Zglinicki, T., and Miwa, S. (2021). How good is the evidence that cellular senescence causes skin ageing? Ageing Res. Rev. 71, 101456. 10.1016/j.arr.2021.101456.

9. Lee, Y.I., Choi, S., Roh, W.S., Lee, J.H., and Kim, T.G. (2021). Cellular senescence and inflammaging in the skin microenvironment. Int. J. Mol. Sci. 22, 3849. 10.3390/ijms22083849.

10. Ressler, S., Bartkova, J., Niederegger, H., Bartek, J., Scharffetter-Kochanek, K., Jansen-Dürr, P., and Wlaschek, M. (2006). p16INK4A is a robust in vivo biomarker of cellular aging in human skin. Aging Cell 5, 379–389. 10.1111/j.1474-9726.2006.00231.x.

11. Dimri, G.P., Lee, X., Basile, G., Acosta, M., Scott, G., Roskelley, C., Medrano, E.E., Linskens, M., Rubelj, I., Pereira-Smith, O., et al. (1995). A biomarker that identifies senescent human cells in culture and in aging skin in vivo. Proc. Natl. Acad. Sci. U.S.A. 92.

12. Waaijer, M.E.C., Gunn, D.A., Adams, P.D., Pawlikowski, J.S., Griffiths, C.E.M., Van Heemst, D., Slagboom, P.E., Westendorp, R.G.J., and Maier, A.B. (2016). P16INK4a positive cells in human skin are indicative of local elastic fiber morphology, facial wrinkling, and perceived age. Journals Gerontol. - Ser. A Biol. Sci. Med. Sci. 71, 1022–1028. 10.1093/gerona/glv114.

13. Ogata, Y., Yamada, T., Hasegawa, S., Sugiura, K., and Akamatsu, H. (2023). Changes of senescent cell accumulation and removal in skin tissue with ageing. Exp. Dermatol. 10.1111/exd.14818.

14. Jevtić, M., Löwa, A., Nováčková, A., Kováčik, A., Kaessmeyer, S., Erdmann, G., Vávrová, K., and Hedtrich, S. (2020). Impact of intercellular crosstalk between epidermal keratinocytes and dermal fibroblasts on skin homeostasis. Biochim. Biophys. Acta - Mol. Cell Res. 1867, 118722. 10.1016/j.bbamcr.2020.118722.

15. Janson, D., Rietveld, M., Willemze, R., and El Ghalbzouri, A. (2013). Effects of serially passaged fibroblasts on dermal and epidermal morphogenesis in human skin equivalents. Biogerontology 14, 131–140. 10.1007/s10522-013-9416-9.

16. Weinmüllner, R., Zbiral, B., Becirovic, A., Stelzer, E.M., Nagelreiter, F., Schosserer, M., Lämmermann, I., Liendl, L., Lang, M., Terlecki-Zaniewicz, L., et al. (2020). Organotypic human skin culture models constructed with senescent fibroblasts show hallmarks of skin aging. npj Aging Mech. Dis. 6. 10.1038/s41514-020-0042-x.

17. Ho, C.Y., and Dreesen, O. (2021). Faces of cellular senescence in skin aging. Mech. Ageing Dev. 198. 10.1016/j.mad.2021.111525.

18. Petrova, N. V., Velichko, A.K., Razin, S. V., and Kantidze, O.L. (2016). Small molecule compounds that induce cellular senescence. Aging Cell 15, 999–1017. 10.1111/acel.12518.

19. Jung, M., Jin, S.G., Zhang, X., Xiong, W., Gogoshin, G., Rodin, A.S., and Pfeifer, G.P. (2015). Longitudinal epigenetic and gene expression profiles analyzed by three-component analysis reveal down-regulation of genes involved in protein translation in human aging. Nucleic Acids Res. 43, 1–14. 10.1093/nar/gkv473.

20. Köhler, F., Bormann, F., Raddatz, G., Gutekunst, J., Corless, S., Musch, T., Lonsdorf, A.S., Erhardt, S., Lyko, F., and Rodríguez-Paredes, M. (2020). Epigenetic deregulation of lamina-associated domains in Hutchinson-Gilford progeria syndrome. Genome Med. 12. 10.1186/s13073-020-00749-y.

21. Haga, M., and Okada, M. (2022). Systems approaches to investigate the role of NF-κB signaling in aging. Biochem. J. 479, 161–183. 10.1042/BCJ20210547.

22. Domínguez-Hüttinger, E., Christodoulides, P., Miyauchi, K., Irvine, A.D., Okada-Hatakeyama, M., Kubo, M., and Tanaka, R.J. (2017). Mathematical modeling of atopic dermatitis reveals “double-switch” mechanisms underlying 4 common disease phenotypes. J. Allergy Clin. Immunol. 139, 1861–1872.e7. 10.1016/j.jaci.2016.10.026.

23. Fey, D., Halasz, M., Dreidax, D., Kennedy, S.P., Hastings, J.F., Rauch, N., Munoz, A.G., Pilkington, R., Fischer, M., Westermann, F., et al. (2015). Signaling pathway models as biomarkers: Patient-specific simulations of JNK activity predict the survival of neuroblastoma patients. Sci. Signal. 8, 1–16. 10.1126/scisignal.aab0990.

24. Imoto, H., Yamashiro, S., and Okada, M. (2022). A text-based computational framework for patient-specific modeling for classification of cancers. iScience 25, 103944. 10.1016/j.isci.2022.103944.

25. Union, P. (2009). Regulation (EC) No 1223/2009 of the European parliament and of the council. Off. J. Eur. Union L 342, 59.

26. Fleischer, J.G., Schulte, R., Tsai, H.H., Tyagi, S., Ibarra, A., Shokhirev, M.N., Huang, L., Hetzer, M.W., and Navlakha, S. (2018). Predicting age from the transcriptome of human dermal fibroblasts. Genome Biol. 19, 1–8. 10.1186/s13059-018-1599-6.

27. Marthandan, S., Menzel, U., Priebe, S., Groth, M., Guthke, R., Platzer, M., Hemmerich, P., Kaether, C., and Diekmann, S. (2016). Conserved genes and pathways in primary human fibroblast strains undergoing replicative and radiation induced senescence. Biol. Res. 49, 1–16. 10.1186/s40659-016-0095-2.

28. Murphy-Ullrich, J.E., and Suto, M.J. (2018). Thrombospondin-1 regulation of latent TGF-β activation: A therapeutic target for fibrotic disease. Matrix Biol. 68–69, 28–43. 10.1016/j.matbio.2017.12.009.

29. Hildebrand, A., Romaris, M., Rasmussen, M., Heinegard, D., Twardzik, D.R., Border, W.A., and Ruoslahti, E. (1994). Interaction of the small interstitial proteoglycans biglycan, decorin and fibromodulin with transforming growth factor β. Biochem. J. 302, 527–534. 10.1042/bj3020527.

30. Embree, M.C., Kilts, T.M., Ono, M., Inkson, C.A., Syed-Picard, F., Karsdal, M.A., Oldberg, Å., Bi, Y., and Young, M.F. (2010). Biglycan and fibromodulin have essential roles in regulating chondrogenesis and extracellular matrix turnover in temporomandibular joint osteoarthritis. Am. J. Pathol. 176, 812–826. 10.2353/ajpath.2010.090450.

31. Pang, X., Dong, N., and Zheng, Z. (2020). Small leucine-rich proteoglycans in skin wound healing. Front. Pharmacol. 10, 1–18. 10.3389/fphar.2019.01649.

32. Garcia-Alonso, L., Holland, C.H., Ibrahim, M.M., Turei, D., and Saez-Rodriguez, J. (2019). Benchmark and integration of resources for the estimation of human transcription factor activities. Genome Res. 29, 1363–1375. 10.1101/gr.240663.118.

33. Wang, Y., Liu, L., Song, Y., Yu, X., and Deng, H. (2022). Unveiling E2F4, TEAD1 and AP-1 as regulatory transcription factors of the replicative senescence program by multi-omics analysis. Protein Cell 13, 742–759. 10.1007/s13238-021-00894-z.

34. Jiao, H., Walczak, B.E., Lee, M.S., Lemieux, M.E., and Li, W.J. (2021). GATA6 regulates aging of human mesenchymal stem/stromal cells. Stem Cells 39, 62–77. 10.1002/stem.3297.

35. Mijit, M., Caracciolo, V., Melillo, A., Amicarelli, F., and Giordano, A. (2020). Role of p53 in the regulation of cellular senescence. Biomolecules 10. 10.3390/biom10030420.

36. Heinz, S., Benner, C., Spann, N., Bertolino, E., Lin, Y.C., Laslo, P., Cheng, J.X., Murre, C., Singh, H., and Glass, C.K. (2010). Simple combinations of lineage-determining transcription factors prime *cis*-regulatory elements required for macrophage and B cell identities. Mol. Cell 38, 576–589. 10.1016/j.molcel.2010.05.004.

37. Yi, Y., Fang, Y., Wu, K., Liu, Y., and Zhang, W. (2020). Comprehensive gene and pathway analysis of cervical cancer progression. Oncol. Lett. 19, 3316–3332. 10.3892/ol.2020.11439.

38. Gunin, A.G., and Golubtzova, N.N. (2019). Transforming growth factor-β (TGF-β) in human skin during aging. Adv. Gerontol. 9, 267–273. 10.1134/S2079057019030068.

39. Coppé, J.P., Desprez, P.Y., Krtolica, A., and Campisi, J. (2010). The senescence-associated secretory phenotype: The dark side of tumor suppression. Annu. Rev. Pathol. Mech. Dis. 5, 99–118. 10.1146/annurev-pathol-121808-102144.

40. Lopes-Paciencia, S., Saint-Germain, E., Rowell, M.C., Ruiz, A.F., Kalegari, P., and Ferbeyre, G. (2019). The senescence-associated secretory phenotype and its regulation. Cytokine 117, 15–22. 10.1016/j.cyto.2019.01.013.

41. Guo, F., Hutchenreuther, J., Carter, D.E., and Leask, A. (2013). TAK1 is required for dermal wound healing and homeostasis. J. Invest. Dermatol. 133, 1646–1654. 10.1038/jid.2013.28.

42. Zhang, Y., Feng, X.-H., and Derynck, R. (1998). Smad3 and Smad4 cooperate with c-Jun/c-Fos to mediate TGF-β-induced transcription. Nature 394, 909–913.

43. Hu, H.-H., Chen, D.-Q., Wang, Y.-N., Feng, Y.-L., Cao, G., Vaziri, N.D., and Zhao, Y.-Y. (2018). New insights into TGF-β/Smad signaling in tissue fibrosis. Chem. Biol. Interact. 292, 76–83. 10.1016/j.cbi.2018.07.008.

44. Sobel, K., Menyhart, K., Killer, N., Renault, B., Bauer, Y., Studer, R., Steiner, B., Bolli, M.H., Nayler, O., and Gatfield, J. (2013). Sphingosine 1-Phosphate (S1P) receptor agonists mediate pro-fibrotic responses in normal human lung fibroblasts via S1P2 and S1P3 receptors and Smad-independent signaling. J. Biol. Chem. 288, 14839–14851. 10.1074/jbc.M112.426726.

45. Flügel-Koch, C., Ohlmann, A., Fuchshofer, R., Welge-Lüssen, U., and Tamm, E.R. (2004). Thrombospondin-1 in the trabecular meshwork: Localization in normal and glaucomatous eyes, and induction by TGF-β1 and dexamethasone in vitro. Exp. Eye Res. 79, 649–663. 10.1016/j.exer.2004.07.005.

46. Daubon, T., Léon, C., Clarke, K., Andrique, L., Salabert, L., Darbo, E., Pineau, R., Guérit, S., Maitre, M., Dedieu, S., et al. (2019). Deciphering the complex role of thrombospondin-1 in glioblastoma development. Nat. Commun. 10. 10.1038/s41467-019-08480-y.

47. Imoto, H., Zhang, S., and Okada, M. (2020). A computational framework for prediction and analysis of cancer signaling dynamics from RNA sequencing data—application to the ErbB receptor signaling pathway. Cancers. 12, 1–13. 10.3390/cancers12102878.

48. Coppe, J.P., Kauser, K., Campisi, J., and Beauséjour, C.M. (2006). Secretion of vascular endothelial growth factor by primary human fibroblasts at senescence. J. Biol. Chem. 281, 29568–29574. 10.1074/jbc.M603307200.

49. Acosta, J.C., Banito, A., Wuestefeld, T., Georgilis, A., Janich, P., Morton, J.P., Athineos, D., Kang, T.W., Lasitschka, F., Andrulis, M., et al. (2013). A complex secretory program orchestrated by the inflammasome controls paracrine senescence. Nat. Cell Biol. 15, 978–990. 10.1038/ncb2784.

50. Schoeberl, B., Pace, E.A., Fitzgerald, J.B., Harms, B.D., Xu, L., Nie, L., Linggi, B., Kalra, A., Paragas, V., Bukhalid, R., et al. (2009). Therapeutically targeting ErbB3: A key node in ligand-induced activation of the ErbB receptor-PI3K axis. Sci. Signal. 2, 1–15. 10.1126/scisignal.2000352.

51. Moiseeva, V., Cisneros, A., Sica, V., Deryagin, O., Lai, Y., Jung, S., Andrés, E., An, J., Segalés, J., Ortet, L., et al. (2023). Senescence atlas reveals an aged-like inflamed niche that blunts muscle regeneration. Nature 613, 169–178. 10.1038/s41586-022-05535-x.

52. Minagawa, S., Araya, J., Numata, T., Nojiri, S., Hara, H., Yumino, Y., Kawaishi, M., Odaka, M., Morikawa, T., Nishimura, S.L., et al. (2011). Accelerated epithelial cell senescence in IPF and the inhibitory role of SIRT6 in TGF-β-induced senescence of human bronchial epithelial cells. Am. J. Physiol. - Lung Cell. Mol. Physiol. 300, 391–401. 10.1152/ajplung.00097.2010.

53. Datto, M.B., Li, Y., Panus, J.F., Howe, D.J., Xiong, Y., and Wang, X.F. (1995). Transforming growth factor β induces the cyclin-dependent kinase inhibitor p21 through a p53-independent mechanism. Proc. Natl. Acad. Sci. U. S. A. 92, 5545–5549. 10.1073/pnas.92.12.5545.

54. Zhang, L.-j., Chen, S.X., Guerrero-Juarez, C.F., Li, F., Tong, Y., Liang, Y., Liggins, M., Chen, X., Chen, H., Li, M., et al. (2019). Age-related loss of innate immune antimicrobial function of dermal fat is mediated by transforming growth factor beta. Immunity 50, 121–136.e5. 10.1016/j.immuni.2018.11.003.

55. Thomas, C., Karagounis, I. V., Srivastava, R.K., Vrettos, N., Nikolos, F., Francois, N., Huang, M., Gong, S., Long, Q., Kumar, S., et al. (2021). Estrogen receptor β-mediated inhibition of actin-based cell migration suppresses metastasis of inflammatory breast cancer. Cancer Res. 81, 2399–2414. 10.1158/0008-5472.CAN-20-2743.

56. Mazur, E.C., Vasquez, Y.M., Li, X., Kommagani, R., Jiang, L., Chen, R., Lanz, R.B., Kovanci, E., Gibbons, W.E., and DeMayo, F.J. (2015). Progesterone receptor transcriptome and cistrome in decidualized human endometrial stromal cells. Endocrinology 156, 2239–2253. 10.1210/en.2014-1566.

57. Sárvári, M., Kalló, I., Hrabovszky, E., Solymosi, N., Tóth, K., Likó, I., Molnár, B., Tihanyi, K., and Liposits, Z. (2010). Estradiol replacement alters expression of genes related to neurotransmission and immune surveillance in the frontal cortex of middle-aged, ovariectomized rats. Endocrinology 151, 3847–3862. 10.1210/en.2010-0375.

58. Ly, D.H., Lockhart, D.J., Lerner, R.A., and Schultz, P.G. (2000). Mitotic misregulation and human aging. Science 287, 2486–2492. 10.1126/science.287.5462.2486.

59. Waldera Lupa, D.M., Kalfalah, F., Safferling, K., Boukamp, P., Poschmann, G., Volpi, E., Götz-Rösch, C., Bernerd, F., Haag, L., Huebenthal, U., et al. (2015). Characterization of skin aging-associated secreted proteins (SAASP) produced by dermal fibroblasts isolated from intrinsically aged human skin. J. Invest. Dermatol. 135, 1954–1968. 10.1038/jid.2015.120.

60. Meijles, D.N., Sahoo, S., Al Ghouleh, I., Amaral, J.H., Bienes-Martinez, R., Knupp, H.E., Attaran, S., Sembrat, J.C., Nouraie, S.M., Rojas, M.M., et al. (2017). The matricellular protein TSP1 promotes human and mouse endothelial cell senescence through CD47 and Nox1. Sci. Signal. 10. 10.1126/scisignal.aaj1784.

61. Zhao, W., Shen, B., Cheng, Q., Zhou, Y., and Chen, K. (2023). Roles of TSP1-CD47 signaling pathway in senescence of endothelial cells: Cell cycle, inflammation and metabolism. Mol. Biol. Rep. 50, 4579–4585. 10.1007/s11033-023-08357-w.

62. Isenberg, J.S., and Roberts, D.D. (2020). Thrombospondin-1 in maladaptive aging responses: A concept whose time has come. Am. J. Physiol. - Cell Physiol. 318, C45–C63. 10.1152/ajpcell.00089.2020.

63. McCabe, M.C., Hill, R.C., Calderone, K., Cui, Y., Yan, Y., Quan, T., Fisher, G.J., and Hansen, K.C. (2020). Alterations in extracellular matrix composition during aging and photoaging of the skin. Matrix Biol. Plus 8, 100041. 10.1016/j.mbplus.2020.100041.

64. Takemon, Y., Chick, J.M., Gyuricza, I.G., Skelly, D.A., Devuyst, O., Gygi, S.P., Churchill, G.A., and Korstanje, R. (2021). Proteomic and transcriptomic profiling reveal different aspects of aging in the kidney. Elife 10. 10.7554/eLife.62585.

65. Kim, S.A., Um, S.J., Kang, J.H., and Hong, K.J. (2001). Expression of thrombospondin-1 in human hepatocarcinoma cell lines and its regulation by transcription factor Jun/AP-1. Mol. Cell. Biochem. 216, 21–29. 10.1023/A:1011022822077.

66. Xu, L., Zhang, Y., Chen, J., and Xu, Y. (2020). Thrombospondin-1: A key protein that induces fibrosis in diabetic complications. J. Diabetes Res. 2020. 10.1155/2020/8043135.

67. Khan, F.U., Owusu-Tieku, N.Y.G., Dai, X., Liu, K., Wu, Y., Tsai, H.I., Chen, H., Sun, C., and Huang, L. (2019). Wnt/β-catenin pathway-regulated fibromodulin expression is crucial for breast cancer metastasis and inhibited by aspirin. Front. Pharmacol. 10, 1–17. 10.3389/fphar.2019.01308.

68. An, W., Zhu, J. W., Jiang, F., Jiang, H., Zhao, J.L., Liu, M.Y., Li, G.X., Shi, X.G., Sun, C., and Li, Z.S. (2020). Fibromodulin is upregulated by oxidative stress through the MAPK/AP-1 pathway to promote pancreatic stellate cell activation. Pancreatology 20, 278–287. 10.1016/j.pan.2019.09.011.

69. Syaidah, R., Tsukada, T., Azuma, M., Horiguchi, K., Fujiwara, K., Kikuchi, M., and Yashiro, T. (2016). Fibromodulin expression in folliculostellate cells and pericytes is promoted by TGFβ signaling in rat anterior pituitary gland. Acta Histochem. Cytochem. 49, 171–179. 10.1267/ahc.16021.

70. Li, H., Venkatraman, L., Narmada, B.C., White, J.K., Yu, H., and Tucker-Kellogg, L. (2017). Computational analysis reveals the coupling between bistability and the sign of a feedback loop in a TGF-β1 activation model. BMC Syst. Biol. 11. 10.1186/s12918-017-0508-z.

71. Pybus, H.J., O’dea, R.D., and Brook, B.S. (2023). A dynamical model of TGF-β activation in asthmatic airways. Math. Med. Biol. A J. IMA, dqad004. 10.1093/imammb/dqad004.

72. Zi, Z., Feng, Z., Chapnick, D.A., Dahl, M., Deng, D., Klipp, E., Moustakas, A., and Liu, X. (2011). Quantitative analysis of transient and sustained transforming growth factor-β signaling dynamics. Mol. Syst. Biol. 7, 1–12. 10.1038/msb.2011.22.

73. Lucarelli, P., Schilling, M., Kreutz, C., Vlasov, A., Boehm, M.E., Iwamoto, N., Steiert, B., Lattermann, S., Wäsch, M., Stepath, M., et al. (2018). Resolving the combinatorial complexity of Smad protein complex formation and its link to gene expression. Cell Syst. 6, 75–89.e11. 10.1016/j.cels.2017.11.010.

74. Khatibi, S., Zhu, H.J., Wagner, J., Tan, C.W., Manton, J.H., and Burgess, A.W. (2017). Mathematical model of TGF-β signaling: Feedback coupling is consistent with signal switching. BMC Syst. Biol. 11, 1–15. 10.1186/s12918-017-0421-5.

75. Schindelin, J., Arganda-Carreras, I., Frise, E., Kaynig, V., Longair, M., Pietzsch, T., Preibisch, S., Rueden, C., Saalfeld, S., Schmid, B., et al. (2012). Fiji: An open-source platform for biological-image analysis. Nat. Methods 9, 676–682. 10.1038/nmeth.2019.

76. Krämer, A., Green, J., Pollard, J.J., and Tugendreich, S. (2014). Causal analysis approaches in Ingenuity Pathway Analysis. Bioinformatics 30, 523–530. 10.1093/bioinformatics/btt703.

77. Di Tommaso, P., Chatzou, M., Floden, E.W., Barja, P.P., Palumbo, E., and Notredame, C. (2017). Nextflow enables reproducible computational workflows. Nat. Biotechnol. 35, 316–319. 10.1038/nbt.3820.

78. Ewels, P.A., Peltzer, A., Fillinger, S., Patel, H., Alneberg, J., Wilm, A., Garcia, M.U., Di Tommaso, P., and Nahnsen, S. (2020). The nf-core framework for community-curated bioinformatics pipelines. Nat. Biotechnol. 38, 276–278. 10.1038/s41587-020-0439-x.

79. Kim, D., Paggi, J.M., Park, C., Bennett, C., and Salzberg, S.L. (2019). Graph-based genome alignment and genotyping with HISAT2 and HISAT-genotype. Nat. Biotechnol. 37, 907–915. 10.1038/s41587-019-0201-4.

80. Danecek, P., Bonfield, J.K., Liddle, J., Marshall, J., Ohan, V., Pollard, M.O., Whitwham, A., Keane, T., McCarthy, S.A., Davies, R.M., et al. (2021). Twelve years of SAMtools and BCFtools. Gigascience 10. 10.1093/gigascience/giab008.

81. Liao, Y., Smyth, G.K., and Shi, W. (2014). featureCounts: An efficient general purpose program for assigning sequence reads to genomic features. Bioinformatics 30, 923–930. 10.1093/bioinformatics/btt656.

82. Quinlan, A.R., and Hall, I.M. (2010). BEDTools: A flexible suite of utilities for comparing genomic features. Bioinformatics 26, 841–842. 10.1093/bioinformatics/btq033.

83. Wu, T., Hu, E., Xu, S., Chen, M., Guo, P., Dai, Z., Feng, T., Zhou, L., Tang, W., Zhan, L., et al. (2021). clusterProfiler 4.0: A universal enrichment tool for interpreting omics data. The Innovation 2, 100141. 10.1016/j.xinn.2021.100141.

84. Love, M.I., Huber, W., and Anders, S. (2014). Moderated estimation of fold change and dispersion for RNA-seq data with DESeq2. Genome Biol. 15, 550. 10.1186/s13059-014-0550-8.

85. Yu, G., Wang, L.G., and He, Q.Y. (2015). ChIP seeker: An R/Bioconductor package for ChIP peak annotation, comparison and visualization. Bioinformatics 31, 2382–2383. 10.1093/bioinformatics/btv145.

86. Hubert, M., Rousseeuw, P.J., and Vanden Branden, K. (2005). ROBPCA: A new approach to robust principal component analysis. Technometrics 47, 64–79. 10.1198/004017004000000563.

87. Team BC, M.B. (2019). TxDb.Hsapiens.UCSC.hg38.knownGene: Annotation package for TxDb object(s). R Packag. version 3.4.6. 10.18129/B9.bioc.TxDb.Hsapiens.UCSC.hg38.knownGene.

88. Henrot, P., Laurent, P., Levionnois, E., Leleu, D., Pain, C., Truchetet, M.E., and Cario, M. (2020). A method for isolating and culturing skin cells: Application to endothelial cells, fibroblasts, keratinocytes, and melanocytes from punch biopsies in systemic sclerosis skin. Front. Immunol. 11, 1–12. 10.3389/fimmu.2020.566607.

89. Storn, R., and Price, K. (1997). Differential evolution – A simple and efficient heuristic for global optimization over continuous spaces. J. Glob. Optim. 11, 341–359. 10.1023/A:1008202821328.

