## Supplementary figures and images for "Positive and negative feedback regulation of the TGF-β1–SMAD4 axis explains two equilibrium states in human skin aging"

### Supplemental figure

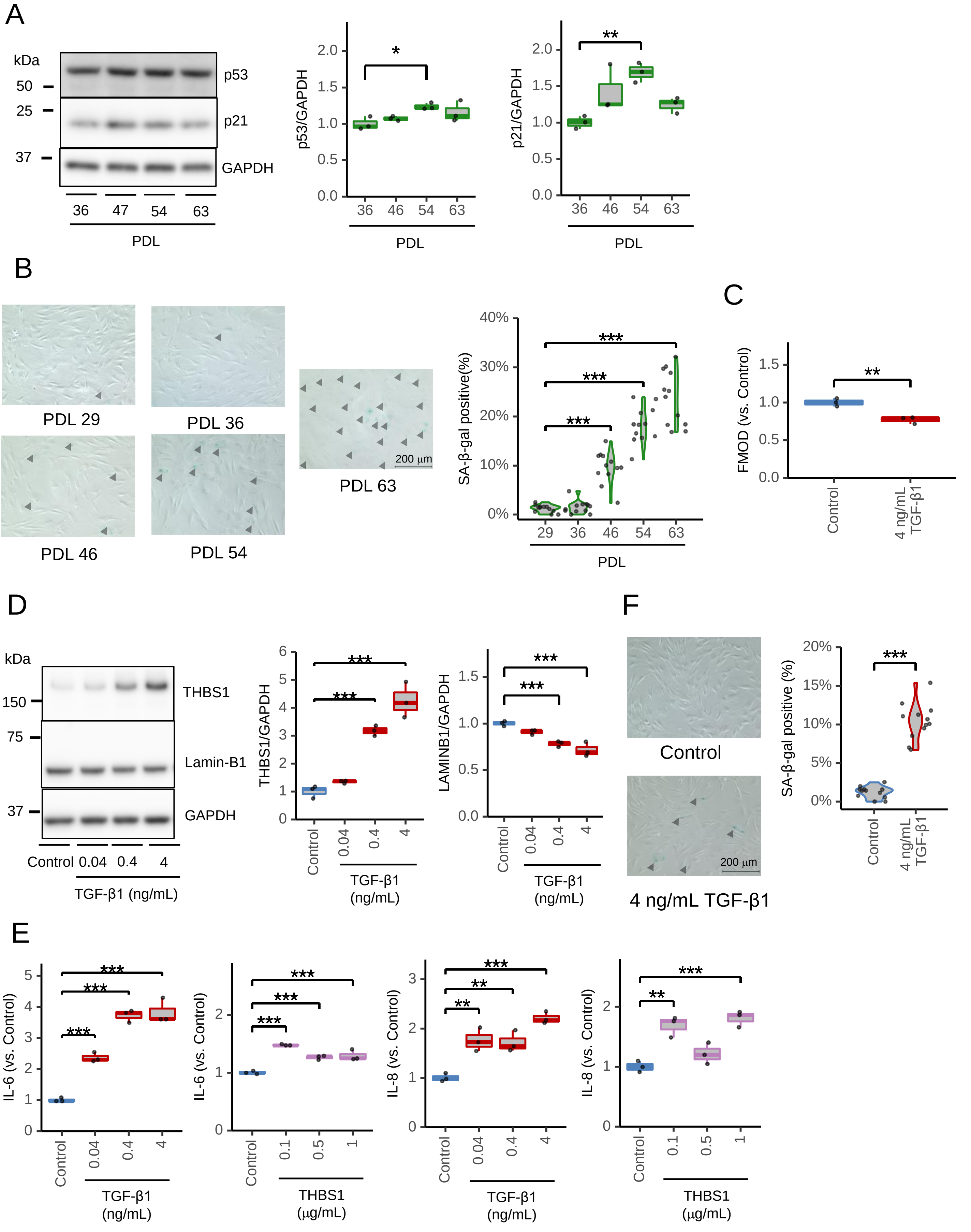

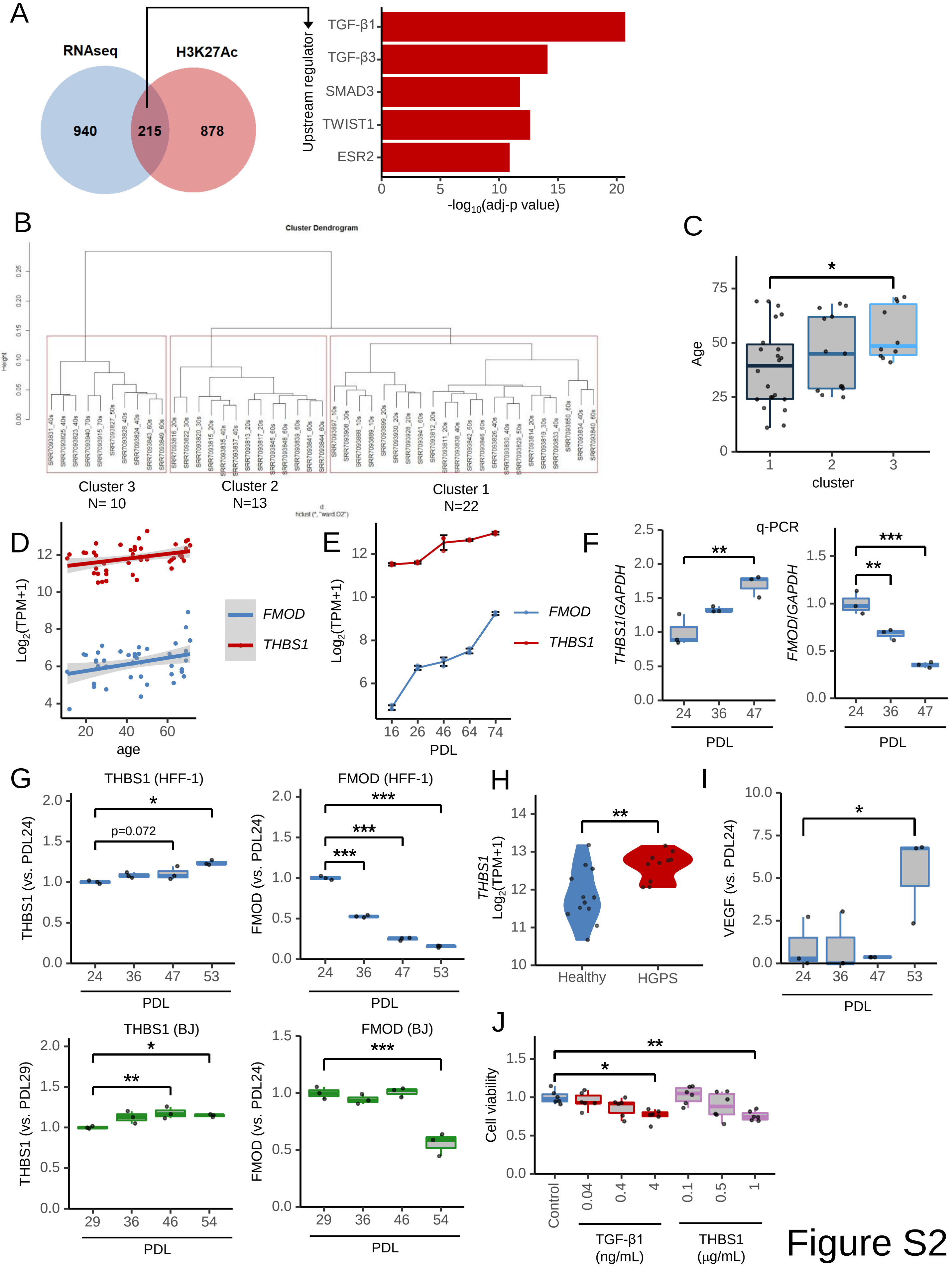

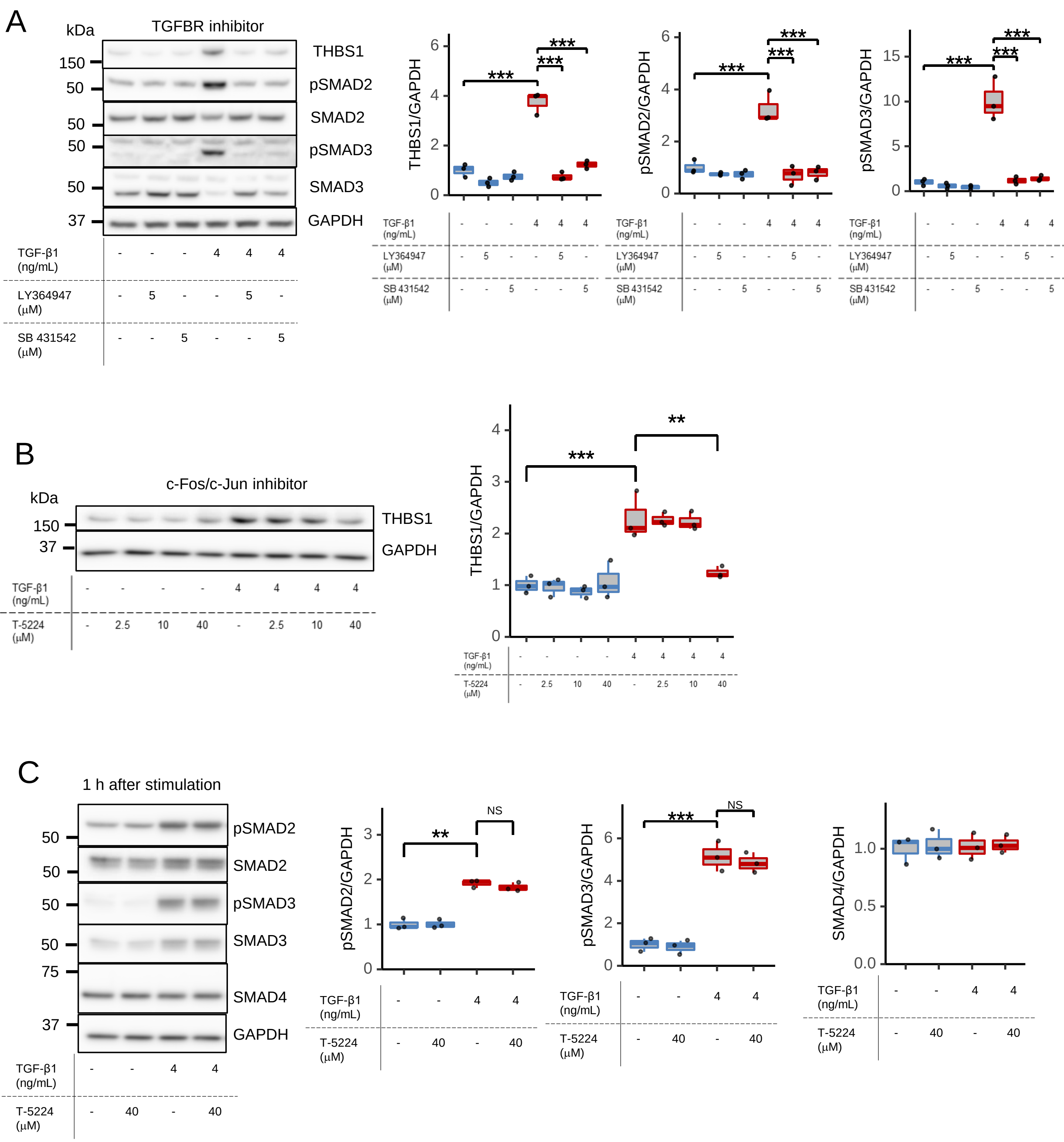

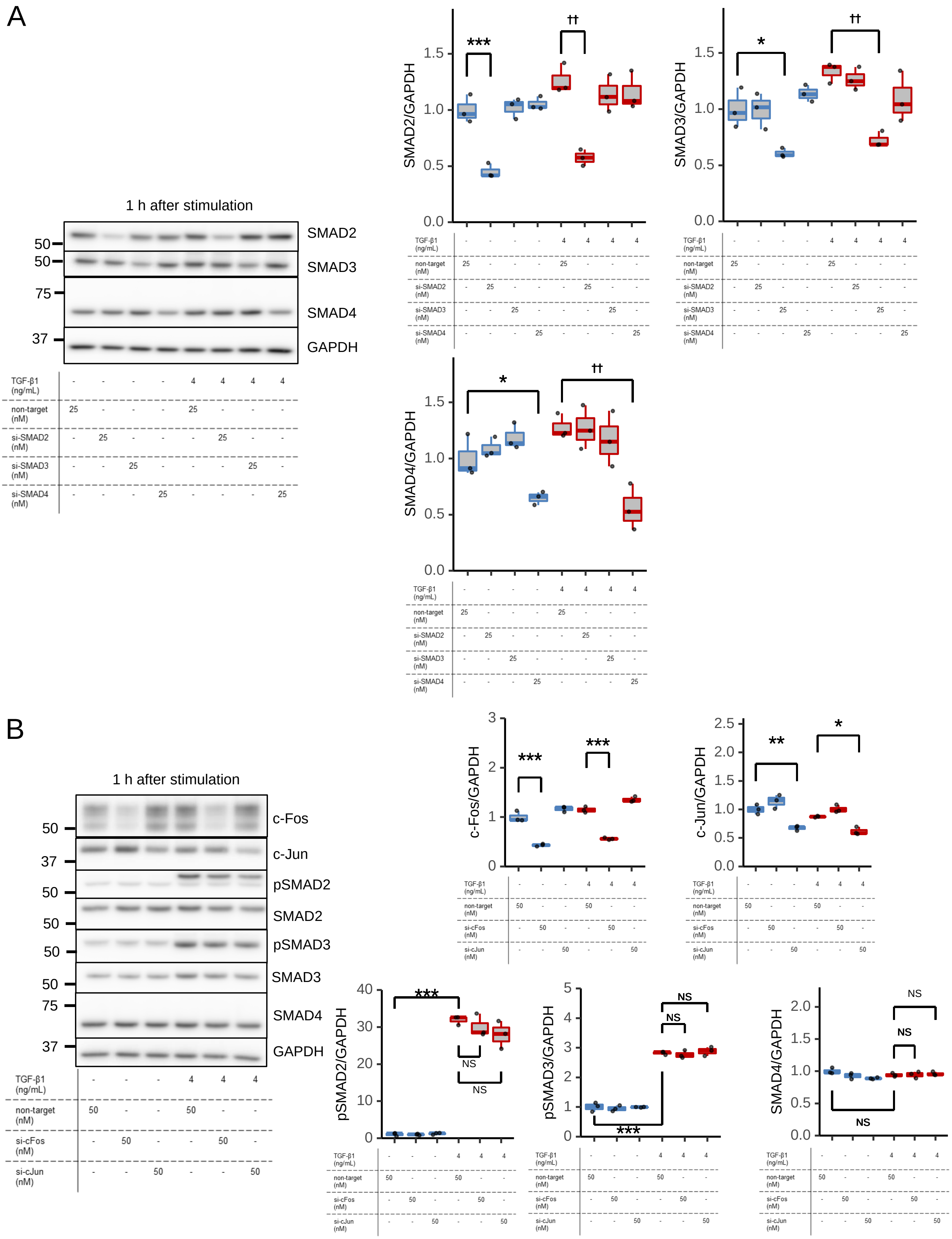

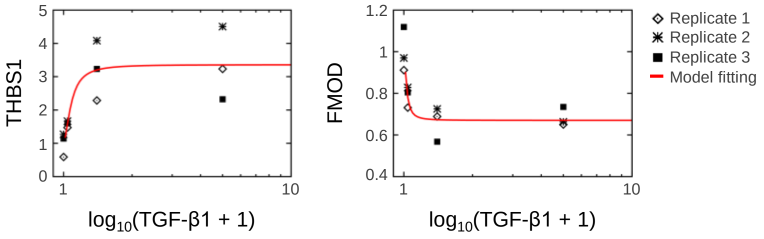

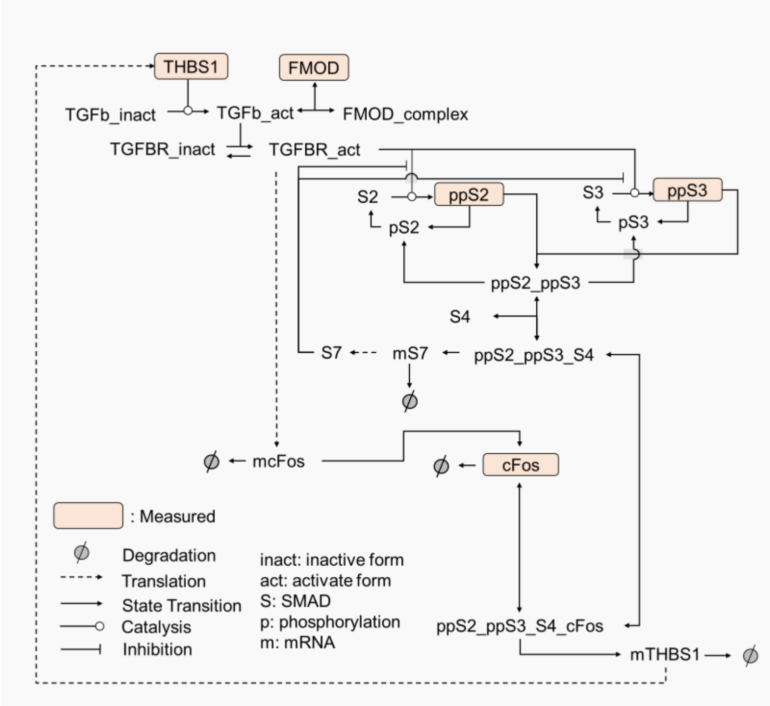

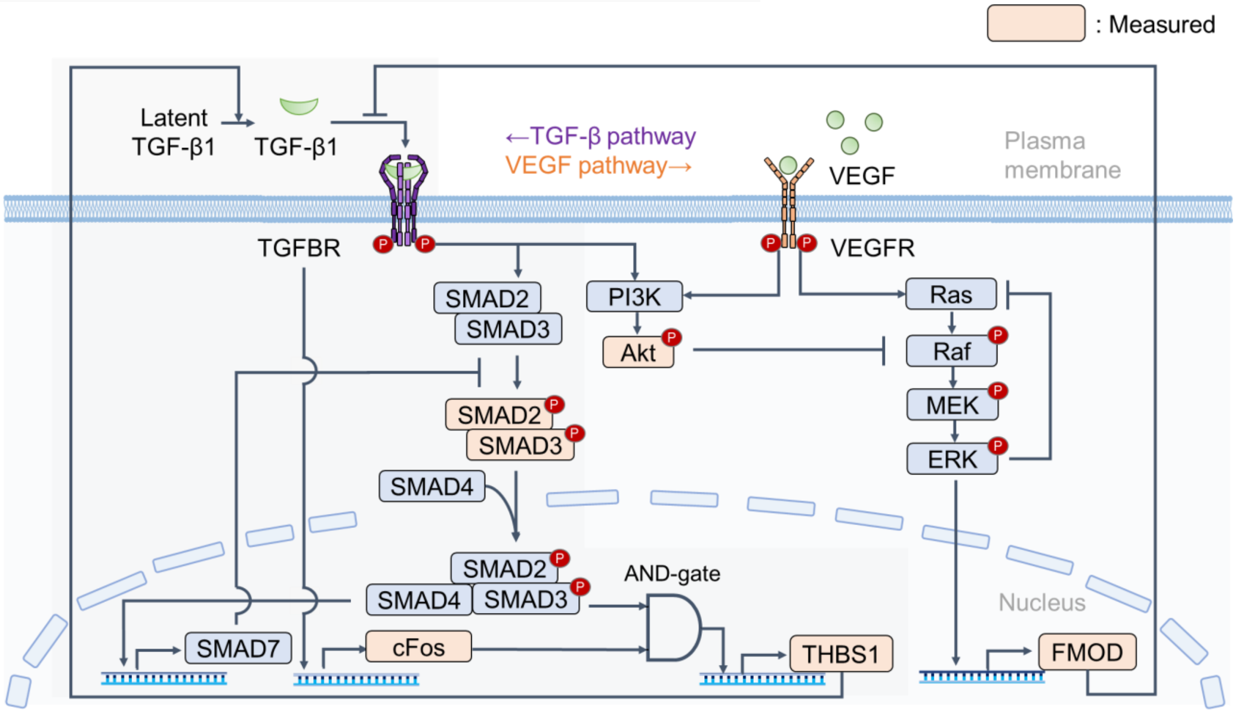

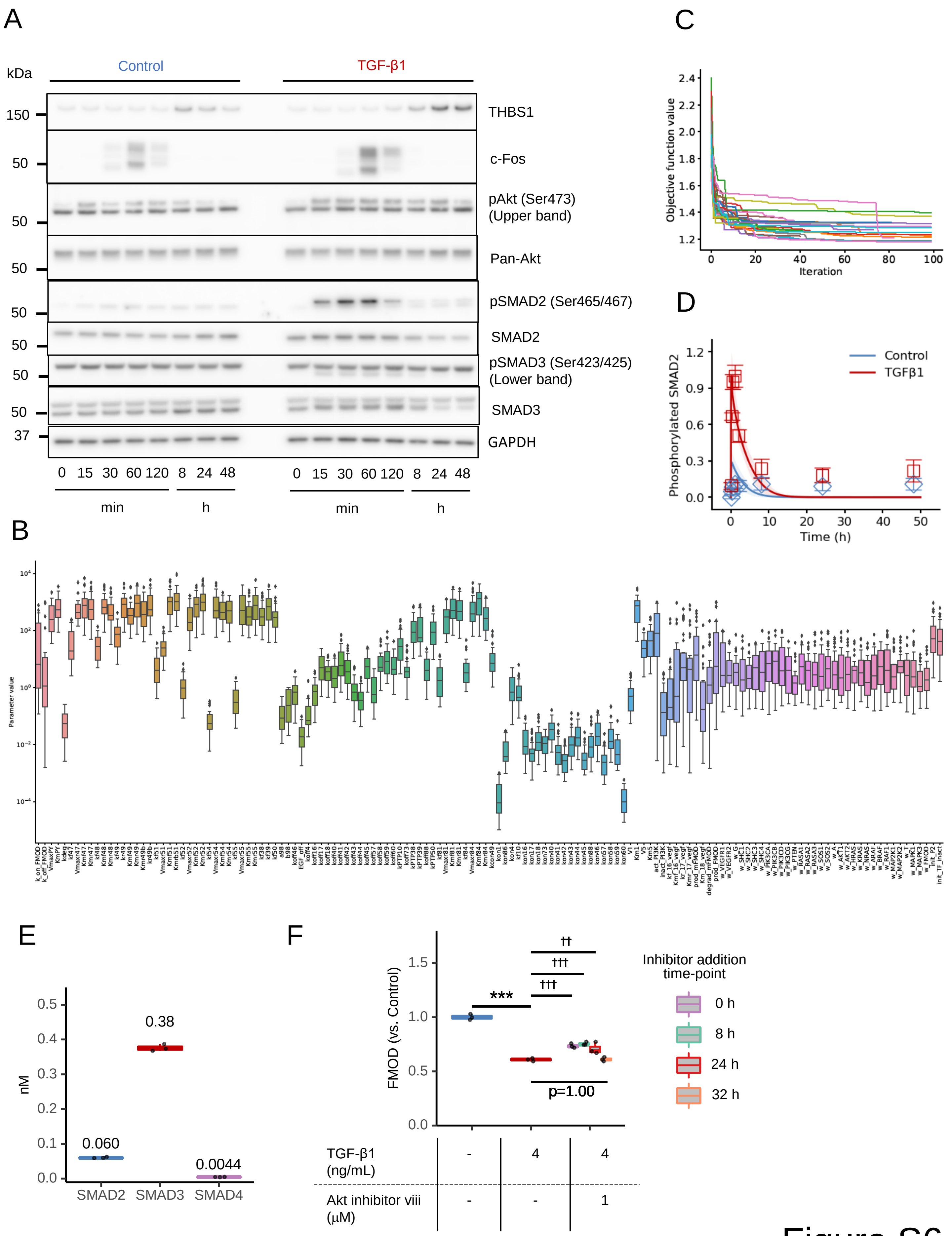
