## Supplement for "Positive and negative feedback regulation of the TGF-β1–SMAD4 axis explains two equilibrium states in human skin aging"

**SUPPLEMENTAL INFORMATION**

**Figure S1. Replication stress induces cellular senescence in BJ cells**

A) Western blot of p53 and p21 in replication-stress-induced BJ cells. (Left panel) Representative image. (Middle panel) Quantification of p53 expression. N = 3, * p < 0.05 (Dunnett’s test). (Right panel) Quantification of p21 expression. N = 3, ** p < 0.01 (Dunnett’s test).
B) (Left panel) SA-β-gal staining image of replication-stress-induced BJ cells. SA-β-gal-positive cells are indicated by an arrowhead (black). (Right panel) Quantification of SA-β-gal: SA-β-gal-positive rate (%) = number of SA-β-gal-positive cells / total number of cells × 100, N = 3 (4 points/well), *** p < 0.001 (Dunnett’s test).
C) FMOD ELISA in TGF-β1-stimulated BJ cells. Cells were treated with TGF-β1 (red) for 48 h and their supernatants were analyzed. N = 3, ** p < 0.01 (Student’s *t*-test).
D) Western blot of THBS1 and Lamin-B1 in TGF-β1-stimulated BJ cells. Cell lysates were collected 48 h after control (blue) or TGF-β1 (red) treatment. (Left panel) Representative image. (Middle panel) Quantification of THBS1. N = 3, *** p < 0.001 (Dunnett’s test). (Right panel) Quantification of Lamin-B1. N = 3, *** p < 0.001 (Dunnett’s test).
E) IL-6 and IL-8 ELISA in TGF-β1- or THBS1-stimulated BJ cells. Cell supernatants were collected at 48 h after control (blue), TGF-β1 (red), or THBS1 (purple) treatment. (First panel) Quantification of IL-6 ELISA with TGF-β1 treatment. N = 3, *** p < 0.001 (Dunnett’s test). (Second panel) Quantification of IL-6 ELISA with THBS1 treatment. N = 3, *** p < 0.01 (Dunnett’s test). (Third panel) Quantification of IL-8 ELISA with TGF-β1 treatment. N = 3, ** p < 0.01, *** p < 0.001 (Dunnett’s test). (Fourth panel) Quantification of IL-8 ELISA with THBS1 treatment. N = 3, ** p < 0.01, *** p < 0.001 (Dunnett’s test).
F) Effect of TGF-β1 simulation on SA-β-gal activity in BJ cells. (Left panel) Representative images. SA-β-gal-positive cells are shown with an arrowhead (black). Scale bar, 200 μm. (Right panel) Quantification of SA-β-gal: SA-β-gal-positive rate (%) = number of SA-β-gal-positive cells / total number of cells × 100, N = 3 (4 points/well), *** p < 0.001 (vs. control, Welch’s *t*-test).

**Figure S2. Analysis and validation of omics data on dermal senescence and aging**

A) (Left panel) Venn diagram showing RNA-seq upregulated DEGs (FC > 1.2, adj-p < 0.05, blue shade) and genes annotated from H3K27Ac differentially gained peaks (Log_2_FC > 0, adj-p < 0.05, red shade). The numbers of genes in each condition are shown. (Right panel) The top five upstream regulators determined using Ingenuity Pathway Analysis (IPA). A right-tailed Fisher’s exact test was used to calculate a p-value of overlap.
B) Clustering of public human skin fibroblast RNA-seq samples. Samples from human arm skin fibroblast (11–71 years of age, N = 45) were clustered into cluster 1 (N = 22), cluster 2 (N = 13), and cluster 3 (N = 10) using the “hclust” function of R with “method = ‘ward.D2.’”
C) Age distribution of clustered human skin fibroblast samples. Median (cluster 1: age 39.5; cluster 2: age 45.0; cluster 3: age 48.5) and average (cluster 1: age 38.9; cluster 2: age 44.8; cluster 3: age 54.5) of each cluster increase with cluster number. N = 3, * p < 0.05 (Wilcoxon rank sum test).
D) Public *in vivo* time-course expression data for *THBS1* and *FMOD*. Gene expression was normalized to Log_2_(transcripts per million [TPM] + 1). N = 45 (11–71 years of age, see donor list in Table S1). Linear regression lines were added using geom_smooth (method = “lm”).
E) Public *in vitro* time-course expression of *THBS1* and *FMOD*. Gene expression was indicated normalized to log_2_(TPM+1). N = 3, mean (SD).
F) qPCR analysis of replication-stress-induced HFF-1 cells. (Left panel) Quantification of *THBS1*. N = 3, ** p < 0.01 (Dunnett’s test). (Right panel) Quantification of *FMOD*. N = 3, ** p < 0.01, *** p < 0.001 (Dunnett’s test).
G) THBS1 and FMOD ELISA of replication-stress-induced HFF-1 (blue) and BJ (green) cell supernatants. (Upper left) Quantification of THBS1 in HFF-1 cells. N = 3, * p < 0.05 (Dunnett’s test). (Upper right) Quantification of FMOD in HFF-1 cells. N = 3, *** p < 0.001 (Dunnett’s test). (Bottom left) Quantification of THBS1 in BJ cells. N = 3, * p < 0.05, ** p < 0.01 (Dunnett’s test). (Bottom right) Quantification of FMOD in BJ cells. N = 3, *** p < 0.001 (Dunnett’s test).
H) *THBS1* expression between healthy and Hutchinson–Gilford progeria syndrome (HGPS) patients. Public RNA-seq data^26^ for healthy (age: 1–9 years, N = 12) and HGPS patients (age: 2–8 years, N = 10) derived from dermal fibroblasts. Gene expression was normalized to log_2_(TPM + 1). ** p < 0.01 (Welch’s *t*-test).
I) VEGF ELISA in replication-stress-induced HFF-1 cells. PDLs were determined from the supernatants after a 48-h culture. N = 3, * p < 0.05 (Dunnett’s test).
J) Cell viability analysis of TGF-β1- and FMOD-stimulated HFF-1 cells. After 24 h of treatment, cell viability was measured using the tetrazolium salt WST-8.

**Figure S3. Effect of TGF-βR and c-Fos/c-Jun inhibitors on SMAD activation and THBS1 expression**

A) Western blot analysis of TGF-β1-stimulated HFF-1 cells treated with TGF-βR inhibitors. Cells were treated with LY364947 and SB431542 with or without TGF-β1 for 48 h and the lysates were analyzed. (Left panel) Representative image. (Right panel) Quantification of THBS1, pSMAD2, and pSMAD3 expression following TGF-βR inhibitor treatment. N = 3, *** p < 0.001 (Tukey’s multiple comparisons).
B) Western blot analysis of TGF-β1-stimulated HFF-1 cells treated with T-5224. Cells were treated with T-5224 with or without TGF-β1 for 48 h and the lysates were analyzed. (Left panel) Representative image. (Right panel) Quantification of THBS1. N = 3, ** p < 0.01, *** p < 0.001 (Tukey’s multiple comparisons).
C) Western blot analysis of TGF-β1-stimulated HFF-1 cells treated with T-5224. Cells were treated with T-5224 with or without TGF-β1 for 1 h and the lysates were analyzed. (Left panel) Representative image. (Right panel) Quantification of pSMAD2, pSMAD3, and SMAD4. N = 3, NS: not significant, ** p < 0.01, *** p < 0.001 (Tukey’s multiple comparisons).

**Figure S4. Effect of SMADs and c-Fos/c-Jun KD on SMAD activation**

A) Western blot analysis of SMAD2, SMAD3, and SMAD4 with SMAD2–SMAD3–SMAD4 KD in HFF-1 cells. Cells were treated with siRNA for each SMAD with or without TGF-β1 for 1 h and the lysates were analyzed. (Left panel) Representative image. (Right panel) Quantification of SMAD2, SMAD3, and SMAD4. N = 3, * p < 0.05, *** p < 0.001 (Dunnett’s test), †† p < 0.01 (Dunnett’s test).
B) Western blot analysis of c-Fos/c-Jun KD in HFF-1 cells. Cell lysates were collected at 48 h after c-Fos/c-Jun KD and control (blue) or TGF-β1 (red) treatment. (Left panel) Representative image. (Right panel) Quantification of c-Fos, c-Jun, pSMAD2, pSMAD3, and SMAD4. N = 3, * p < 0.05, ** p < 0.01, *** p < 0.001 (Tukey’s multiple comparisons).

**Figure S5. Comprehensive skin aging ordinary differential equation (ODE) model**

A) The model parameters for the core transcription factor network consisting of TGF-β1, THBS1, and FMOD were trained on the experimental expression of THBS1 and FMOD upon treatment with different concentrations (0.04, 0.4, or 4 ng/mL) of TGF-β1 (Figure 3A). The x- and y-axes represent the steady state expressions of TGF-β1 and THBS1 (left panel) or FMOD (right panel). The points (◇: Replicate 1, $⋇$: Replicate 2, ■: Replicate 3) indicate experimental data, and red solid lines indicate model fitting results. The western blot images can be found in Figure 3A.
B) TGF-β and VEGF signaling pathways in an ODE model. (Upper panel) Diagram of molecular interactions in the TGF-β signaling model. A model of TGF-βR activation, SMAD phosphorylation, and SMAD complex formation was developed for the TGF-β pathway. (Lower panel) In addition to the TGF-β signaling model, the process of VEGFR activation to ERK phosphorylation was described using the Imoto model^47^ (see details in the Methods section).

**Figure S6. Mathematical modeling and validation of integrated TGF-β–VEGF signaling** **pathway**

A) Experimental time-course western blot image with/without TGF-β1 treatment in HFF-1 cells. HFF-1 was stimulated with/without TGF-β1 and the lysates were collected at 0 min, 15 min, 30 min, 60 min, 120 min, 8 h, 24 h, and 48 h. Representative image (N = 3).
B) Estimated parameter values of 30 parameter sets for integrated TGF-β–VEGF signaling model.
C) Objective function traces from 30 optimization runs for the integrated TGF-β–VEGF signaling model.
D) Validation of mechanistic model using phosphorylated SMAD2. The model reproduced phosphorylated SMAD2 dynamics. The points (Control: blue squares, TGF-β1: red squares) indicate experimental data, solid lines indicate the average simulation of 30 parameter sets, and shade areas indicate SD. The error bar represents SD. N = 3, mean (SD).
E) SMADs ELISA of the initial state of HFF-1 cells. After serum starvation, cell lysates were collected for quantification of initial SMAD expression. N = 3.
F) Quantification of FMOD in Akt inhibitor VIII-treated HFF-1 cells by ELISA. HFF-1 cells were initially treated with TGF-β1 and Akt inhibitor VIII was added at 0 h (purple), 8 h (green), 24 h (bright red), and 32 h (orange) and supernatants were collected at 48 h for quantification. N = 3, *** p < 0.001 (Student’s *t*-test), †† p < 0.01, ††† p < 0.001 (Dunnett’s test).

**TABLES**

**Table S1. Donor list used for public RNA-seq analysis**

| Run | Age (years) | Disease | Gender | Source | Ethnicity | Sample filtering |
| --- | --- | --- | --- | --- | --- | --- |
| SRR7093887 | 11 | Healthy | Female | Skin; Arm | Caucasian | Omitted |
| SRR7093888 | 11 | Healthy | Female | Skin; Arm | Caucasian | Used |
| SRR7093889 | 12 | Healthy | Male | Skin; Arm | Caucasian | Used |
| SRR7093897 | 19 | Healthy | Male | Skin; Arm | Caucasian | Used |
| SRR7093899 | 20 | Healthy | Female | Skin; Arm | Caucasian | Used |
| SRR7093811 | 24 | Healthy | Female | Skin; Arm | Caucasian | Used |
| SRR7093930 | 24 | Healthy | Male | Skin; Arm | Caucasian | Used |
| SRR7093812 | 25 | Healthy | Male | Skin; Arm | Caucasian | Used |
| SRR7093813 | 25 | Healthy | Female | Skin; Arm | Caucasian | Used |
| SRR7093928 | 25 | Healthy | Female | Skin; Arm | Caucasian | Used |
| SRR7093814 | 26 | Healthy | Male | Skin; Arm | Caucasian | Used |
| SRR7093815 | 26 | Healthy | Female | Skin; Arm | Caucasian | Used |
| SRR7093816 | 28 | Healthy | Male | Skin; Arm | Caucasian | Used |
| SRR7093817 | 29 | Healthy | Male | Skin; Arm | Caucasian | Used |
| SRR7093818 | 29 | Healthy | Male | Skin; Arm | Caucasian | Omitted |
| SRR7093819 | 30 | Healthy | Male | Skin; Arm | Caucasian | Used |
| SRR7093820 | 30 | Healthy | Female | Skin; Arm | Caucasian | Used |
| SRR7093821 | 30 | Healthy | Male | Skin; Arm | Caucasian | Omitted |
| SRR7093822 | 30 | Healthy | Male | Skin; Arm | Caucasian | Used |
| SRR7093908 | 37 | Healthy | Male | Skin; Arm | Caucasian | Used |
| SRR7093909 | 39 | Healthy | Male | Skin; Arm | Caucasian | Omitted |
| SRR7093825 | 41 | Healthy | Male | Skin; Arm | Caucasian | Used |
| SRR7093832 | 41 | Healthy | Female | Skin; Arm | Caucasian | Omitted |
| SRR7093830 | 42 | Healthy | Female | Skin; Arm | Caucasian | Used |
| SRR7093823 | 43 | Healthy | Female | Skin; Arm | Caucasian | Used |
| SRR7093833 | 43 | Healthy | Male | Skin; Arm | Caucasian | Used |
| SRR7093824 | 44 | Healthy | Male | Skin; Arm | Caucasian | Used |
| SRR7093834 | 44 | Healthy | Male | Skin; Arm | Caucasian | Used |
| SRR7093835 | 45 | Healthy | Male | Skin; Arm | Caucasian | Used |
| SRR7093828 | 46 | Healthy | Male | Skin; Arm | Caucasian | Used |
| SRR7093836 | 46 | Healthy | Male | Skin; Arm | Caucasian | Omitted |
| SRR7093837 | 46 | Healthy | Male | Skin; Arm | Caucasian | Used |
| SRR7093826 | 47 | Healthy | Male | Skin; Arm | Caucasian | Used |
| SRR7093831 | 47 | Healthy | Male | Skin; Arm | Caucasian | Used |
| SRR7093838 | 47 | Healthy | Male | Skin; Arm | Caucasian | Used |
| SRR7093827 | 50 | Healthy | Female | Skin; Arm | Caucasian | Used |
| SRR7093829 | 50 | Healthy | Male | Skin; Arm | Caucasian | Used |
| SRR7093910 | 51 | Healthy | Male | Skin; Arm | Caucasian | Omitted |
| SRR7093912 | 55 | Healthy | Male | Skin; Arm | Caucasian | Omitted |
| SRR7093839 | 61 | Healthy | Male | Skin; Arm | Caucasian | Used |
| SRR7093840 | 62 | Healthy | Female | Skin; Arm | Caucasian | Used |
| SRR7093841 | 62 | Healthy | Female | Skin; Arm | Caucasian | Used |
| SRR7093842 | 63 | Healthy | Male | Skin; Arm | Caucasian | Used |
| SRR7093843 | 64 | Healthy | Male | Skin; Arm | Caucasian | Used |
| SRR7093844 | 66 | Healthy | Male | Skin; Arm | Caucasian | Used |
| SRR7093845 | 67 | Healthy | Male | Skin; Arm | Caucasian | Used |
| SRR7093846 | 67 | Healthy | Male | Skin; Arm | Caucasian | Used |
| SRR7093847 | 68 | Healthy | Male | Skin; Arm | Caucasian | Omitted |
| SRR7093848 | 68 | Healthy | Male | Skin; Arm | Caucasian | Used |
| SRR7093939 | 68 | Healthy | Male | Skin; Arm | Caucasian | Omitted |
| SRR7093849 | 69 | Healthy | Male | Skin; Arm | Caucasian | Used |
| SRR7093850 | 69 | Healthy | Female | Skin; Arm | Caucasian | Used |
| SRR7093941 | 69 | Healthy | Male | Skin; Arm | Caucasian | Used |
| SRR7093940 | 70 | Healthy | Male | Skin; Arm | Caucasian | Used |
| SRR7093915 | 71 | Healthy | Female | Skin; Arm | Caucasian | Used |

**Table S2. Donor list of dermal tissue**

| Donor ID | Age (years) | Part | Gender | Race | Frozen time (Count) | Body mass index |
| --- | --- | --- | --- | --- | --- | --- |
| #1 | 23 | Breast | Female | Caucasian | 1 | 23 |
| #2 | 27 | Abdominal | Female | Caucasian | 1 | 24 |
| #3 | 31 | Breast | Female | Caucasian | 1 | 31 |
| #4 | 46 | Abdominal | Female | Caucasian | 1 | 21 |
| #5 | 61 | Breast | Female | Caucasian | 1 | 29 |
| #6 | 63 | Abdominal | Female | Caucasian | 1 | 28 |

**Table S3. Donor list used for public RNA-seq analysis (healthy and Hutchinson–Gilford progeria syndrome [HGPS] samples)**

| Run | Age (years) | Disease | Gender | Source | Ethnicity |
| --- | --- | --- | --- | --- | --- |
| SRR7093809 | 1 | Healthy | Male | Skin; Foreskin | Asian |
| SRR7093874 | 1 | Healthy | Male | Skin; Unspecified | Caucasian |
| SRR7093875 | 2 | Healthy | Female | Skin; Unspecified | Caucasian |
| SRR7093876 | 3 | Healthy | Male | Skin; Inguinal area | Latino/Hispanic |
| SRR7093877 | 3 | Healthy | Male | Skin; Unspecified | NA |
| SRR7093878 | 5 | Healthy | Male | Skin; Umbilical cord area | Black |
| SRR7093879 | 6 | Healthy | Male | Skin; Inguinal area | Black |
| SRR7093880 | 7 | Healthy | Male | Skin; Inguinal area | Black |
| SRR7093881 | 7 | Healthy | Male | Skin; Unspecified | Caucasian |
| SRR7093882 | 8 | Healthy | Male | Skin; Unspecified | Caucasian |
| SRR7093883 | 8 | Healthy | Male | Skin; Inguinal area | Caucasian |
| SRR7093884 | 9 | Healthy | Female | Skin; Unspecified | Black |
| SRR7093942 | 8 | HGPS | Female | Skin; Leg | Caucasian |
| SRR7093943 | 8 | HGPS | Male | NA | NA |
| SRR7093944 | 2 + 3 months | HGPS | Female | NA | NA |
| SRR7093945 | 3 + 9 months | HGPS | Female | NA | NA |
| SRR7093946 | 4 + 8 months | HGPS | Female | NA | NA |
| SRR7093947 | 8 + 6 months | HGPS | Male | NA | NA |
| SRR7093948 | 6 + 11 months | HGPS | Female | NA | NA |
| SRR7093949 | 5 | HGPS | Female | NA | NA |
| SRR7093950 | 8 + 10 months | HGPS | Male | NA | NA |
| SRR7093951 | 3 | HGPS | Female | NA | NA |

**Table S4. Gene list**

| Gene symbol | Transcripts per million (TPM, average) | TPM (SD) |
| --- | --- | --- |
| TGFBR1 | 64.76 | 2.24 |
| TGFBR2 | 95.81 | 4.21 |
| SMAD7 | 20.62 | 3.33 |
| FOS | 1.17 | 0.62 |
| THBS1 | 4522.39 | 40.42 |
| FMOD | 20.88 | 0.66 |
| FLT1 | 101.53 | 1.82 |
| KDR | 0.02 | 0.02 |
| GRB2 | 128.45 | 0.34 |
| SHC1 | 401.13 | 8.79 |
| SHC2 | 1.16 | 0.49 |
| SHC3 | 4.28 | 0.38 |
| SHC4 | 1.26 | 0.30 |
| PIK3CA | 16.36 | 0.80 |
| PIK3CB | 15.10 | 0.89 |
| PIK3CD | 32.27 | 1.57 |
| PIK3CG | 0.00 | 0.00 |
| PTEN | 51.20 | 1.38 |
| RASA1 | 85.47 | 4.19 |
| RASA2 | 26.93 | 0.49 |
| RASA3 | 92.99 | 3.57 |
| GAB1 | 7.09 | 0.54 |
| SOS1 | 25.34 | 1.14 |
| SOS2 | 20.01 | 0.81 |
| AKT1 | 84.62 | 2.25 |
| AKT2 | 41.27 | 0.82 |
| HRAS | 101.55 | 4.41 |
| KRAS | 42.43 | 1.21 |
| NRAS | 113.71 | 3.80 |
| ARAF | 85.60 | 1.46 |
| BRAF | 4.73 | 0.20 |
| RAF1 | 79.97 | 1.25 |
| MAP2K1 | 73.15 | 1.57 |
| MAP2K2 | 114.17 | 3.90 |
| PTPN1 | 84.13 | 3.12 |
| MAPK1 | 78.20 | 1.11 |
| MAPK3 | 75.96 | 5.75 |
