## Supplementary material for "Positive and negative feedback regulation of the TGF-β1–SMAD4 axis explains two equilibrium states in human skin aging": Key_Resource_Table

**KEY RESOURCES TABLE**

| **REAGENT or RESOURCE** | **SOURCE** | **IDENTIFIER** |
| --- | --- | --- |
| Antibodies |  |  |
| Anti-THBS1 | Cell Signaling Technology | Cat# 37879; RRID: AB_2799123 |
| Anti-FMOD | ProteinTech | Cat# 60108-1-Ig; RRID: AB_2105538 |
| Anti-Lamin B1 | ProteinTech | Cat# 12987-1-AP; RRID: AB_2136290 |
| Anti-p53 | Cell Signaling Technology | Cat# 2524; RRID: AB_331743 |
| Anti-p21 | Cell Signaling Technology | Cat# 2946; RRID: AB_2260325 |
| Anti-c-Fos | Cell Signaling Technology | Cat# 2250; RRID: AB_2247211 |
| Anti-c-Jun | Cell Signaling Technology | Cat# 9165; RRID: AB_2130165 |
| Anti-SMAD2 | Cell Signaling Technology | Cat# 5339; RRID: AB_10626777 |
| Anti-phosphorylated SMAD2 (Ser465/467) | Cell Signaling Technology | Cat# 3108; RRID: AB_490941 |
| Anti-SMAD3 | Cell Signaling Technology | Cat# 9523; RRID: AB_2193182 |
| Anti-phosphorylated SMAD3 (Ser423/425) | Cell Signaling Technology | Cat# 9520; RRID: AB_2193207 |
| Anti-SMAD4 | Cell Signaling Technology | Cat# 46535; RRID: AB_2736998 |
| Anti-phosphorylated Akt (Ser473) | Cell Signaling Technology | Cat# 9271; RRID: AB_329825 |
| Anti-GAPDH | Medical & Biological Laboratories | Cat# M171-3; RRID: AB_10597731 |
| Anti-GAPDH | ProteinTech | Cat# 10494-1-Ap; RRID: AB_2263076 |
| Anti-H3K27Ac | Abcam | Cat# ab177178; RRID: AB_2828007 |
| Chemicals, Peptides, and Recombinant Proteins |  |  |
| D-PBS (-) (1X) | Nacalai Tesque | Cat# 14249-24 |
| HBSS | Thermo Fisher Scientific | Cat# 14025092 |
| Dulbecco’s modified Eagle’s medium | ATCC | Cat# 30-2002 |
| Fetal bovine serum | Corning | Cat# 35-010-CV |
| Antibiotic–antimycotic | Thermo Fisher Scientific | Cat# 15240062 |
| Trypsin/EDTA | ATCC | Cat# 30-2101 |
| BAMBANKER^®^ | NIPPON Genetics | Cat# CS-04-001 |
| RIPA Buffer | Thermo Fisher Scientific | Cat# 89900 |
| Halt™ Protease and Phosphatase inhibitor | Thermo Fisher Scientific | Cat# 0078442 |
| Trypan blue | Thermo Fisher Scientific | Cat# 15250-061 |
| Cell counter | WakenBtech | Cat# WC2-100 |
| Hoechst^®^ 33342 | DOJINDO | Cat# 346-07951 |
| Opti-MEM™ I | Thermo Fisher Scientific | Cat# 31985-070 |
| Lipofectamine RNAiMax | Thermo Fisher Scientific | Cat# 13778150 |
| Precast gel 7.5–15% | Nacalai Tesque | Cat# 13066-44 |
| Wide Precast Gel 7.5–15% | Biocraft | Cat# MDG-287 |
| Tris/Glycine/SDS Buffer | Bio-Rad | Cat# 1610732 |
| 4× Laemmli sample buffer | Bio-Rad | Cat# 1610747 |
| 2-Mercaptoethanol | Bio-Rad | Cat# 1610710 |
| Precision Plus Protein WesternC blotting standard | Bio-Rad | Cat# 1610376 |
| Clarity Western ECL Substrate | Bio-Rad | Cat# 170-5060 |
| Clarity Max Western ECL Substrate | Bio-Rad | Cat# 1705062 |
| Blocking buffer | Bio-Rad | Cat# 12010020 |
| iBlot^®^2 PVDF | Thermo Fisher Scientific | Cat# IB24001 |
| Reconstitution buffer (0.1% BSA in 4 mM HCl PBS) | R&D Systems | Cat# RB04 |
| Recombinant human TGF-β1 | R&D Systems | Cat# 7754-BH-005 |
| Recombinant human THBS1 | R&D Systems | Cat# 3074-TH-050 |
| Recombinant human FMOD | Abcam | Cat# ab152392 |
| Recombinant human VEGF165 | ProteinTech | Cat# HZ-1038 |
| Dimethyl sulfoxide | Fujifilm | Cat# 041-29351 |
| LY364947 | Fujifilm | Cat# 123-05981 |
| Akt inhibitor VIII | Cayman Chemical | Cat# 14870 |
| LY294002 | Calbiochem | Cat# 440202 |
| T-5224 | Selleck Chemicals | Cat# S28966 |
| Tocriscreen Kinase Inhibitor Toolbox | Tocris Bioscience | Cat# 3514 |
| ZM306416HCl | Tocris Bioscience | Cat# 3514-2499 |
| Ki8751 | Tocris Bioscience | Cat# 3514-2542 |
| GW5074 | Tocris Bioscience | Cat# 3514-1381 |
| U0126 | Tocris Bioscience | Cat# 3514-1144 |
| SB 431542 | Tocris Bioscience | Cat# 3514-1614 |
| Formaldehyde | Thermo Fisher Scientific | Cat# 28908 |
| Proteinase K | Thermo Fisher Scientific | Cat# 26160 |
| Dispase^®^ II | Roche | Cat# 04942078001 |
| Critical Commercial Assays |  |  |
| Pierce^TM^ BCA Protein Assay Kit | Thermo Fisher Scientific | Cat# 23227 |
| NucleoSpin^®^ RNA kit | Macherey-Nagel GmbH & Co. | REF# 740955.50 |
| ReverTra Ace^®^ qPCR RT Master Mix | Toyobo Life Science | Cat# FSQ-201 |
| KOD SYBR^®^ qPCR kit | Toyobo Life Science | Cat# QKD-201 |
| SimpleChIP^®^ Enzymatic Chromatin IP kit | Cell Signaling Technology | Cat# 9003 |
| iDeal ChIP-seq kit for Histones | Diagenode | Cat# C01010171 |
| MinElute^®^ PCR Purification Kit | Qiagen | Cat# 28004 |
| Bioanalyzer Agilent High-Sensibility DNA Kit | Agilent | P/N: G2938-85004 |
| NEBNext^®^ Poly(A) mRNA Magnetic Isolation Module | New England Biolabs | Cat# E7490 |
| NEBNext^®^ Ultra™ ll Directional RNA Library Prep Kit | New England Biolabs | Cat# E7760 |
| ATAC-Seq Kit | Active Motif | Cat# 53150 |
| NEBNext^®^ Ultra II DNA Library Prep Kit for Illumina | New England Biolabs | Cat# 7645 |
| Cell Counting Kit-8 | DOJINDO | Cat# 343-07623 |
| SA-β-gal Detection Kit | BioVision | Cat# K320-250 |
| CycLex^®^ Cellular BrdU ELISA Kit Ver.2 | Medical & Biological Laboratories | Cat# CY-1142V2 |
| THBS1 ELISA Kit | R&D Systems | Cat# DTSP10 |
| FMOD ELISA Kit | Abcam | Cat# ab275895 |
| TGF-β1 ELISA Kit | R&D Systems | Cat# DB100B |
| VEGF ELISA Kit | R&D Systems | Cat# DVE00 |
| SMAD2 ELISA Kit | Abcam | Cat# ab260065 |
| SMAD3 ELISA Kit | Abcam | Cat# ab264624 |
| SMAD4 ELISA Kit | Abcam | Cat# ab253211 |
| IL-6 ELISA Kit | R&D Systems | Cat# D6050 |
| IL-8 ELISA Kit | Abcam | Cat# ab214030 |
| Deposited Data |  |  |
| RNA-seq: HFF-1 (PDL 24, PDL 36, PDL 47) | This Paper | DRA016119 |
| ChIP-seq: H3K27Ac HFF-1 (PDL 24, PDL 36, PDL 47); Input HFF-1 (PDL 24, PDL 36, PDL 47) | This Paper | DRA016119 |
| ATAC-seq: HFF-1 (PDL 24, PDL 36, PDL 47) | This Paper | DRA016119 |
| RNA-seq: HFF-1 (PDL 24) initial condition | This Paper | DRA016119 |
| Experimental models: Cell lines |  |  |
| Human dermal fibroblast HFF-1 | ATCC | SCRC-1041; RRID: CVCL_3285 |
| Human dermal fibroblast BJ | ATCC | CRL-2522; RRID: CVCL_3653 |
| Experimental models: Organisms/Strains |  |  |
| Frozen human full thickness skin | Biopredic International | Cat# TRA1FTR0, TRA1FTR2 |
| Oligonucleotides |  |  |
| qRT-PCR forward primer: *THBS1* 5′-TCCCCATCCAAAGCGTCTTC-3′ | This paper | N/A |
| qRT-PCR reverse primer: *THBS1* 5 ′-ACCACGTTGTTGTCAAGGGT-3′ | This paper | N/A |
| qRT-PCR forward primer: *FMOD* 5′-GGACGTGGTCACTCTCTGAA-3′ | This paper | N/A |
| qRT-PCR reverse primer: *FMOD* 5′-GGCTCGTAGGTCTCATACGG-3′ | This paper | N/A |
| qRT-PCR forward primer: *GAPDH* 5′-GTCTCCTCTGACTTCAACAGCG-3′ | OriGene | Cat# NM_002046 |
| qRT-PCR reverse primer: *GAPDH* 5′-ACCACCCTGTTGCTGTAGCCAA-3′ | OriGene | Cat# NM_002046 |
| ON-TARGET plus Non-targeting siRNA | Dharmacon | Cat# D-001810-02-20 |
| ON-TARGET plus Human c-Fos siRNA - SMARTpool | Dharmacon | Cat# L-003265-00-0010 |
| ON-TARGET plus Human c-Jun siRNA - SMARTpool | Dharmacon | Cat# L-003268-00-0010 |
| ON-TARGET plus Human SMAD2 siRNA - SMARTpool | Dharmacon | Cat# L-003561-00-0020 |
| ON-TARGET plus Human SMAD3 siRNA - SMARTpool | Dharmacon | Cat# L-020067-00-0020 |
| ON-TARGET plus Human SMAD4 siRNA - SMARTpool | Dharmacon | Cat# L-003902-00-0020 |
| Software and Algorithms |  |  |
| ImageJ Fiji version 1.52p | Schindelin et al.^75^ | http://fiji.sc  RRID: SCR_002285 |
| Biomass version 0.5.2 | Imoto et al.^47^ | https://github.com/biomass-dev/biomass  RRID: N/A |
| Gnuplot vesion 5.4 | Williams et al. | http://www.gnuplot.info/  RRID: SCR_008619 |
| R version 4.2.1 | The R Foundation | https://r-project.org  RRID: SCR_001905 |
| QIAGEN Ingenuity Pathway Analysis version 81348237 | Krämer et al.^76^ | https://digitalinsights.qiagen.com/products-overview/discovery-insights-portfolio/analysis-and-visualization/qiagen-ipa/  RRID: SCR_008653 |
| Nextflow version 21.10.6 | Tommaso et al.^77^ | https://www.nextflow.io/index.html  RRID: N/A |
| nfcore/chipseq version 1.2.2 | Ewels et al.^78^ | https://nf-co.re/chipseq/1.2.2  https://zenodo.org/record/7139814#.Y4lJDXbP1aQ  RRID: N/A |
| nfcore/atacseq version 1.2.1 | Ewels et al.^78^ | https://nf-co.re/atacseq/1.2.1  https://zenodo.org/record/7384115#.Y4lJv3bP1aQ  RRID: N/A |
| Trim Galore! version 0.6.6 | The Babraham Institute | http://www.bioinformatics.babraham.ac.uk/projects/trim_galore/  RRID: SCR_011847 |
| hisat2 version 2.2.1 | Kim et al.^79^ | http://ccb.jhu.edu/software/hisat2/index.shtml  RRID: SCR_015530 |
| Samtools version 1.9 | Danecek et al.^80^ | https://github.com/samtools/samtools  RRID: SCR_002105 |
| Subread version 2.0.1 | Liao et al.^81^ | https://subread.sourceforge.net/  RRID: SCR_009803 |
| BEDTools version 2.30.0 | Quinlan et al.^82^ | https://github.com/arq5x/bedtools2  RRID: SCR_006646 |
| HOMER version 4.11 | Heinz et al.^36^ | http://homer.ucsd.edu/homer/  RRID: SCR_010881 |
| DoRothEA version 1.8.0 | Garcia-Alonso et al.^32^ | https://saezlab.github.io/dorothea/  RRID: N/A |
| clusterProfiler version 4.4.4 | Wu et al.^83^ | https://bioconductor.org/packages/release/bioc/html/clusterProfiler.html  RRID: SCR_016884 |
| DESeq2 version 1.36.0 | Love et al.^84^ | https://bioconductor.org/packages/release/bioc/html/DESeq2.html  RRID: SCR_015687 |
| ChIPseeker version 1.32.1 | Yu et al.^85^ | https://bioconductor.org/packages/release/bioc/html/ChIPseeker.html  RRID: SCR_021322 |
| rrcov version 1.5.2 | Hubert et al.^86^ | https://cran.r-project.org/package=rrcov  RRID: N/A |
