## Supplementary material for "Positive and negative feedback regulation of the TGF-β1–SMAD4 axis explains two equilibrium states in human skin aging": MethodS1

```
TGF_b model

    #Abbreviation
    -----------------------------
    v: rate reaction
    t: time
    y: state variable
    x: constant parameter
    dydt: time derivative of y
    -----------------------------

    #Description of variable names
    -----------------------------
    - inact: inactive form
    - act: active form
    - S: SMAD
    - p: phosphorylation
    - m: mRNA
    -----------------------------

    # Rate reactions
        v[1] = x[C.kf_1_TGFbeta]*y[V.THBS1]*y[V.TGFb_inact]/(x[C.Kmf_1_TGFbeta] + y[V.TGFb_inact])
        v[2] = x[C.k_on_FMOD]*y[V.FMOD]*y[V.TGFb_act]
        v[3] = x[C.Rec_act]*y[V.TGFBR_inact]*y[V.TGFb_act]
        v[4] = x[C.pRec_debind]*y[V.TGFBR_act]
        v[5] = x[C.kf_2_TGFbeta]*y[V.S2]*y[V.TGFBR_act]/(x[C.Kmf_2_TGFbeta]*(1+y[V.S7]/x[C.k_inhibit_TGF])+y[V.S2])
        v[6] = x[C.S_dephosphos]*y[V.ppS2]
        v[7] = x[C.S_dephos]*y[V.pS2]
        v[8] = x[C.kf_3_TGFbeta]*y[V.S3]*y[V.TGFBR_act]/(x[C.Kmf_3_TGFbeta]*(1+y[V.S7]/x[C.k_inhibit_TGF])+y[V.S3])
        v[9] = x[C.S_dephosphos]*y[V.ppS3]
        v[10] = x[C.S_dephos]*y[V.pS3]
        v[11] = x[C.k_on_ppS2_ppS3]*y[V.ppS3]*y[V.ppS2]
        v[12] = x[C.S_dephosphos]*y[V.ppS2_ppS3]
        v[13] = x[C.k_on_ppS2_ppS3_S4]*y[V.ppS2_ppS3]*y[V.S4]
        v[14] = x[C.prod_mS7]*y[V.ppS2_ppS3_S4]**x[C.n1_TGF]/(x[C.Km_1_TGF]**x[C.n1_TGF]+y[V.ppS2_ppS3_S4]**x[C.n1_TGF])
        v[15] = x[C.mS7_turn]*y[V.mS7]
        v[16] = x[C.prod_S7]*y[V.mS7]
        v[17] = x[C.prod_mcFOS]*y[V.TGFBR_act]**x[C.n2_TGF]/(x[C.Km_2_TGF]**x[C.n2_TGF]+y[V.TGFBR_act]**x[C.n2_TGF])
        v[18] = x[C.mcFOS_turn]*y[V.mcFOS]
        v[19] = x[C.prod_cFOS]*y[V.mcFOS]
        v[20] = x[C.k_off_FMOD]*y[V.FMOD_complex]
        v[21] = x[C.k_off_ppS2_ppS3_S4]*y[V.ppS2_ppS3_S4]
        v[22] = x[C.k_off_ppS2_ppS3_S4_cFOS]*y[V.ppS2_ppS3_S4_cFOS]
        v[23] = x[C.k_on_ppS2_ppS3_S4_cFOS]*y[V.ppS2_ppS3_S4]*y[V.cFOS]
        v[24] = x[C.prod_mTHBS1]*y[V.ppS2_ppS3_S4_cFOS]**x[C.n4_TGF]/(x[C.Km_4_TGF]**x[C.n4_TGF]+y[V.ppS2_ppS3_S4_cFOS]**x[C.n4_TGF])
        v[25] = x[C.THBS1_turn]*y[V.mTHBS1]
        v[26] = x[C.prod_THBS1]*y[V.mTHBS1]
        v[27] = x[C.degrad_cFOS]*y[V.cFOS]

    # Equations of the model
        dydt[V.TGFb_inact] = - v[1]
        dydt[V.TGFb_act] = + v[1] - v[2] - v[3] + v[20]
        dydt[V.FMOD] = - v[2] + v[20]
        dydt[V.FMOD_complex] = + v[2] - v[20]
        dydt[V.TGFBR_inact] = - v[3] + v[4]
        dydt[V.TGFBR_act] = + v[3] - v[4]
        dydt[V.S2] = - v[5] + v[7]
        dydt[V.S3] = - v[8] + v[10]
        dydt[V.S4] = - v[13] + v[21]
        dydt[V.pS2] = + v[6] - v[7] + v[12]
        dydt[V.pS3] = + v[9] - v[10] + v[12]
        dydt[V.ppS2] = + v[5] - v[6] - v[11]
        dydt[V.ppS3] = + v[8] - v[9] - v[11]
        dydt[V.ppS2_ppS3] = + v[11] - v[12] - v[13] + v[21]
        dydt[V.ppS2_ppS3_S4] = + v[13] - v[23] - v[21] + v[22]
        dydt[V.mS7] = + v[14] - v[15]
        dydt[V.S7] = + v[16]
        dydt[V.mcFOS] = + v[17] - v[18]
        dydt[V.cFOS] = + v[19] - v[27] - v[23] + v[22]
        dydt[V.ppS2_ppS3_S4_cFOS] = + v[23] - v[22]
        dydt[V.mTHBS1] = + v[24] - v[25]
        dydt[V.THBS1] = + v[26]

    # Best fit parameter
        x[C.kf_1_TGFbeta] = 4.923e-01
        x[C.Kmf_1_TGFbeta] = 2.529e+00
        x[C.k_on_FMOD] = 1.000e+00
        x[C.k_off_FMOD] = 1.000e+00
        x[C.Rec_act] = 5.970e-01
        x[C.pRec_debind] = 8.082e-03
        x[C.S2tot] = 6.000e-02
        x[C.S3tot] = 3.800e-01
        x[C.S4tot] = 4.400e-03
        x[C.kf_2_TGFbeta] = 4.664e-01
        x[C.Kmf_2_TGFbeta] = 4.889e+00
        x[C.k_inhibit_TGF] = 1.569e+00
        x[C.S_dephosphos] = 1.001e-01
        x[C.S_dephos] = 1.947e+00
        x[C.kf_3_TGFbeta] = 9.264e-01
        x[C.Kmf_3_TGFbeta] = 1.004e-01
        x[C.k_on_ppS2_ppS3] = 8.995e-01
        x[C.k_on_ppS2_ppS3_S4] = 9.071e-01
        x[C.k_off_ppS2_ppS3_S4] = 3.579e-01
        x[C.prod_mS7] = 2.069e-01
        x[C.n1_TGF] = 1.000e+00
        x[C.Km_1_TGF] = 2.299e-01
        x[C.mS7_turn] = 8.564e+00
        x[C.prod_S7] = 2.258e-01
        x[C.prod_mcFOS] = 1.848e+01
        x[C.n2_TGF] = 1.000e+00
        x[C.Km_2_TGF] = 2.049e-02
        x[C.mcFOS_turn] = 2.056e-02
        x[C.prod_cFOS] = 1.134e-01
        x[C.k_on_ppS2_ppS3_S4_cFOS] = 6.391e-01
        x[C.k_off_ppS2_ppS3_S4_cFOS] = 2.785e-01
        x[C.prod_mTHBS1] = 2.157e+02
        x[C.n4_TGF] = 1.000e+00
        x[C.Km_4_TGF] = 1.102e-01
        x[C.THBS1_turn] = 5.464e-03
        x[C.prod_THBS1] = 3.524e+00
        x[C.w_THBS1] = 1.234e-01
        x[C.w_TGFBR1] = 2.282e-01
        x[C.w_TGFBR2] = 3.128e-01
        x[C.w_SMAD7] = 1.021e-01
        x[C.w_cFOS] = 1.047e-01
        x[C.degrad_cFOS] = 1.000e-01

    # None zero initial values for 'Control' condition
    # y0[V.TGFb_act] =0 for 'Control'
        y0[V.TGFb_inact] = 0.013
        y0[V.TGFBR_inact] = 4.475e+01
        y0[V.S2] = 6.000e-02
        y0[V.S3] = 3.800e-01
        y0[V.S4] = 4.400e-03
        y0[V.mS7] = 2.105e+00
        y0[V.mcFOS] = 1.225e-01
        y0[V.mTHBS1] = 5.581e+02
    # None zero initial values for 'TGFβ1' condition
    # y0[V.TGFb_act] = 0.0902 for 'TGFβ1'
        y0[V.TGFb_inact] = 0.013
        y0[V.TGFb_act] =0.0902
        y0[V.TGFBR_inact] = 4.475e+01
        y0[V.S2] = 6.000e-02
        y0[V.S3] = 3.800e-01
        y0[V.S4] = 4.400e-03
        y0[V.mS7] = 2.105e+00
        y0[V.mcFOS] = 1.225e-01
        y0[V.mTHBS1] = 5.581e+02
```
