## Supplementary material for "Positive and negative feedback regulation of the TGF-β1–SMAD4 axis explains two equilibrium states in human skin aging": MethodS2

```
TGFb_VEGF_model
    - TGF_b model
    - VEGF_model #Modified from Imoto et al., Cancers (2020)

    #Abbreviation
    -----------------------------
    v: rate reaction
    t: time
    y: state variable
    x: constant parameter
    dydt: time derivative of y
    -----------------------------

    #Description of variable names
    -----------------------------
    1. TGF_b model (v[1] ~ v[27])
        - inact: inactive form
        - act: active form
        - S: SMAD
        - p: phosphorylation
        - m: mRNA

    2. VEGF_model (v[28] ~ v[79])
        - E: VEGF
        - E1: VEGFR
        - G: Grb2.
        - sigmaG: Grb2-containing species in which the Grb2 SH2 domain is bound to
        tyrosine-phosphorylated receptor dimer (EijP) or to tyrosine-phosphorylated Shc (SP),
        and both Grb2 SH3 domains are unbound.
        - S: Shc.
        - sigmaS: Shc-containing species in which the Shc SH2 domain is bound to
        tyrosine-phosphorylated receptor dimer (EijP) or to membrane-localized,
        tyrosine-phosphorylated GAB1 (AP), and Shc is unphosphorylated.
        - I: PI-3K.
        - sigmaI: PI-3K-containing species in which PI-3K is bound to
        tyrosine-phosphorylated receptor dimer (EijP) or to membrane-localized,
        tyrosine-phosphorylated GAB1 (AP).
        - R: RasGAP.
        - sigmaR: RasGAP-containing species in which RasGAP is bound to
        tyrosine-phosphorylated receptor dimer (EijP) or to membrane-localized,
        tyrosine-phosphorylated GAB1 (AP), but is not phosphorylated.
        - T: PTP-1B.
        - sigmaT: PTP-1B-containing species in which PTP-1B is bound to
        tyrosine-phosphorylated receptor dimer (EijP) or to membrane-localized,
        tyrosine-phosphorylated GAB1 (AP).
        - A: GAB1.
        - sigmaA: Gab1-containing species in which the GAB1 PH domain is bound to
        PIP3 or the PRD is bound to Grb2, and GAB1 is unphosphorylated.
        - O: SOS.
        - sigmaO: SOS-containing species that are bound to
        a membrane- localized N-terminal SH3 domain of Grb2.
        - c: cytoplasmic
        - n: nuclear
    -----------------------------

    # extracellular volume to cytoplasmic volume ratio
    VeVc = 33.3

    # fraction definitions
        if y[V.sigmaS] + y[V.sigmaSP] + y[V.sigmaSP_G] > 0.0:
            fsigmaS = y[V.sigmaS]/(y[V.sigmaS] + y[V.sigmaSP] + y[V.sigmaSP_G])
        else:
            fsigmaS = 0.0
        if y[V.sigmaG] + y[V.sigmaG_A] + y[V.sigmaG_O] + y[V.A_sigmaG_O] > 0.0:
            fsigmaG = y[V.sigmaG]/(y[V.sigmaG] + y[V.sigmaG_A] + y[V.sigmaG_O] + y[V.A_sigmaG_O])
        else:
            fsigmaG = 0.0


        if y[V.sigmaA] + y[V.sigmaAP] + y[V.sigmaAP_S] + y[V.sigmaAP_R] + y[V.sigmaAP_I] + y[V.sigmaAP_T] > 0.0:
            fsigmaA = y[V.sigmaA]/(y[V.sigmaA] + y[V.sigmaAP] + y[V.sigmaAP_S] + y[V.sigmaAP_R] + y[V.sigmaAP_I] + y[V.sigmaAP_T])
        else:
            fsigmaA = 0.0


        if y[V.sigmaR] + y[V.sigmaRP] > 0.0:
            fsigmaR = y[V.sigmaR]/(y[V.sigmaR] + y[V.sigmaRP])
        else:
            fsigmaR = 0.0

        sigmaEP = y[V.E11P]
        if sigmaEP > 0.0:
            f11 = y[V.E11P]/sigmaEP
        else:
            f11 = 0.0
    v = {}

    # Rate reactions
    # TGFb_model
        v[1] = x[C.kf_1_TGFbeta]*y[V.THBS1]*y[V.TGFb_inact]/(x[C.Kmf_1_TGFbeta] + y[V.TGFb_inact])
        v[2] = x[C.k_on_FMOD]*y[V.FMOD]*y[V.TGFb_act]
        v[3] = x[C.Rec_act]*y[V.TGFBR_inact]*y[V.TGFb_act]
        v[4] = x[C.pRec_debind]*y[V.TGFBR_act]
        v[5] = x[C.kf_2_TGFbeta]*y[V.S2]*y[V.TGFBR_act]/(x[C.Kmf_2_TGFbeta]*(1+y[V.S7]/x[C.k_inhibit_TGF])+y[V.S2])
        v[6] = x[C.S_dephosphos]*y[V.ppS2]
        v[7] = x[C.S_dephos]*y[V.pS2]
        v[8] = x[C.kf_3_TGFbeta]*y[V.S3]*y[V.TGFBR_act]/(x[C.Kmf_3_TGFbeta]*(1+y[V.S7]/x[C.k_inhibit_TGF])+y[V.S3])
        v[9] = x[C.S_dephosphos]*y[V.ppS3]
        v[10] = x[C.S_dephos]*y[V.pS3]
        v[11] = x[C.k_on_ppS2_ppS3]*y[V.ppS3]*y[V.ppS2]
        v[12] = x[C.S_dephosphos]*y[V.ppS2_ppS3]
        v[13] = x[C.k_on_ppS2_ppS3_S4]*y[V.ppS2_ppS3]*y[V.S4]
        v[14] = x[C.prod_mS7]*y[V.ppS2_ppS3_S4]**x[C.n1_TGF]/(x[C.Km_1_TGF]**x[C.n1_TGF]+y[V.ppS2_ppS3_S4]**x[C.n1_TGF])
        v[15] = x[C.mS7_turn]*y[V.mS7]
        v[16] = x[C.prod_S7]*y[V.mS7]
        v[17] = x[C.prod_mcFOS]*y[V.TGFBR_act]**x[C.n2_TGF]/(x[C.Km_2_TGF]**x[C.n2_TGF]+y[V.TGFBR_act]**x[C.n2_TGF])
        v[18] = x[C.mcFOS_turn]*y[V.mcFOS]
        v[19] = x[C.prod_cFOS]*y[V.mcFOS]
        v[20] = x[C.k_off_FMOD]*y[V.FMOD_complex]
        v[21] = x[C.k_off_ppS2_ppS3_S4]*y[V.ppS2_ppS3_S4]
        v[22] = x[C.k_off_ppS2_ppS3_S4_cFOS]*y[V.ppS2_ppS3_S4_cFOS]
        v[23] = x[C.k_on_ppS2_ppS3_S4_cFOS]*y[V.ppS2_ppS3_S4]*y[V.cFOS]
        v[24] = x[C.prod_mTHBS1]*y[V.ppS2_ppS3_S4_cFOS]**x[C.n4_TGF]/(x[C.Km_4_TGF]**x[C.n4_TGF]+y[V.ppS2_ppS3_S4_cFOS]**x[C.n4_TGF])
        v[25] = x[C.THBS1_turn]*y[V.mTHBS1]
        v[26] = x[C.prod_THBS1]*y[V.mTHBS1]
        v[27] = x[C.degrad_cFOS]*y[V.cFOS]
    # VEGF_model
        v[28] = (x[C.kon1]*y[V.E]*y[V.E1] - x[C.EGF_off]*y[V.E_E1])
        v[29] = (x[C.kon4]*y[V.E_E1]*y[V.E_E1] - x[C.koff4]*y[V.E11])
        v[30] = (x[C.kf10]*y[V.E11] - x[C.VmaxPY]*y[V.E11P]/(x[C.KmPY] + y[V.E11P]) - x[C.kPTP10]*y[V.sigmaT]*y[V.E11P])
        v[31] = (4*x[C.kon16]*y[V.E11P]*y[V.G] - x[C.koff16]*fsigmaG*y[V.E11G])
        v[32] = (8*x[C.kon17]*y[V.E11P]*y[V.S] - x[C.koff17]*fsigmaS*y[V.E11S])
        v[33] = (2*x[C.kon18]*y[V.E11P]*y[V.R] - x[C.koff18]*fsigmaR*y[V.E11R])
        v[34] = (x[C.kf38]*y[V.sigmaS]*sigmaEP - x[C.VmaxPY]*y[V.sigmaSP]/(x[C.KmPY] + y[V.sigmaSP]) - x[C.kPTP38]*y[V.sigmaT]*y[V.sigmaSP])
        v[35] = (x[C.kf39]*y[V.sigmaA]*sigmaEP - x[C.VmaxPY]*y[V.sigmaAP]/(x[C.KmPY] + y[V.sigmaAP]) - x[C.kPTP39]*y[V.sigmaT]*y[V.sigmaAP])
        v[36] = (x[C.kon40]*y[V.sigmaG]*y[V.O] - x[C.koff40]*y[V.sigmaG_O])
        v[37] = (x[C.kon41]*y[V.sigmaG]*y[V.A] - x[C.koff41]*y[V.sigmaG_A]*fsigmaA)
        v[38] = (x[C.kon42]*y[V.sigmaSP]*y[V.G] - x[C.koff42]*y[V.sigmaSP_G]*fsigmaG)
        v[39] = (3*x[C.kon43]*y[V.sigmaAP]*y[V.S] - x[C.koff43]*y[V.sigmaAP_S]*fsigmaS)
        v[40] = (3*x[C.kon44]*y[V.sigmaAP]*y[V.I] - x[C.koff44]*y[V.sigmaAP_I])
        v[41] = (2*x[C.kon45]*y[V.sigmaAP]*y[V.R] - x[C.koff45]*y[V.sigmaAP_R]*fsigmaR)
        v[42] = (x[C.kon46]*y[V.P3]*y[V.A] - x[C.koff46]*y[V.P3_A]*fsigmaA)
        v[43] = (x[C.kf47]*y[V.P3]*y[V.Akt]/(x[C.Kmf47] + y[V.Akt]) - x[C.Vmaxr47]*y[V.Aktstar]/(x[C.Kmr47] + y[V.Aktstar]))
        v[44] = (x[C.kf48]*(1 - y[V.fint]*f11)*y[V.sigmaI]*y[V.P2]/(x[C.Kmf48] + y[V.P2]) - 3*x[C.PTEN]*y[V.P3]/(x[C.Kmr48] + y[V.P3]))
        v[45] = (x[C.kf49]*y[V.sigmaO]*y[V.RsD]/(x[C.Kmf49] + y[V.RsD]) - x[C.kr49]*y[V.sigmaR]*y[V.RsT]/(x[C.Kmr49] + y[V.RsT]) - x[C.kr49b]*y[V.sigmaRP]*y[V.RsT]/(x[C.Kmr49b] + y[V.RsT]) - x[C.kcon49]*y[V.RsT])
        v[46] = (x[C.kf50]*y[V.sigmaR]*sigmaEP - x[C.VmaxPY]*y[V.sigmaRP]/(x[C.KmPY] + y[V.sigmaRP]) - x[C.kPTP50]*y[V.sigmaT]*y[V.sigmaRP])
        v[47] = (x[C.kf51]*y[V.RsT]*y[V.Raf]/(x[C.Kmf51] + y[V.Raf]) - x[C.Vmaxr51]*y[V.Rafstar]*y[V.Aktstar]/(x[C.Kmrb51] + y[V.Rafstar]))
        v[48] = (x[C.kf52]*y[V.Rafstar]*y[V.MEK]/(x[C.Kmf52] + y[V.MEK]) - x[C.Vmaxr52]*y[V.ppMEKc]/(x[C.Kmr52] + y[V.ppMEKc]))
        v[49] = (x[C.kf54]*y[V.O]*y[V.ppERKc]/(x[C.Kmf54] + y[V.O]) - x[C.Vmaxr54]*y[V.OP]/(x[C.Kmr54] + y[V.OP]))
        v[50] = (x[C.kf55]*y[V.A]*y[V.ppERKc]/(x[C.Kmf55] + y[V.A]) - x[C.Vmaxr55]*y[V.AP]/(x[C.Kmr55] + y[V.AP]))
        v[51] = (x[C.kon57]*y[V.P3_A]*y[V.G] - x[C.koff57]*y[V.sigmaA_G])
        v[52] = (x[C.kon58]*y[V.sigmaA_G]*y[V.O] - x[C.koff58]*y[V.sigmaA_G_O])
        v[53] = (x[C.kon59]*y[V.sigmaG_O]*y[V.A] - x[C.koff59]*y[V.A_sigmaG_O]*fsigmaA)
        v[54] = (x[C.kon60]*y[V.sigmaG_A]*y[V.O] - x[C.koff60]*y[V.A_sigmaG_O])
        v[55] = (4*x[C.kon73]*y[V.E11P]*y[V.T] - x[C.koff73]*y[V.E11T])
        v[56] = (x[C.kf81]*y[V.E1]*y[V.ppERKc]/(x[C.Kmf81] + y[V.E1]) - x[C.Vmaxr81]*y[V.E1_PT]/(x[C.Kmr81] + y[V.E1_PT]))
        v[57] = (x[C.kf84]*y[V.E_E1]*y[V.ppERKc]/(x[C.Kmf84] + y[V.E_E1]) - x[C.Vmaxr84]*y[V.E_E1_PT]/(x[C.Kmr84] + y[V.E_E1_PT]))
        v[58] = (x[C.kon86]*y[V.E]*y[V.E1_PT] - x[C.EGF_off]*y[V.E_E1_PT])
        v[59] = (2*x[C.kon88]*y[V.sigmaAP]*y[V.T] - x[C.koff88]*y[V.sigmaAP_T])
        v[60] = x[C.kdeg]*y[V.E11P]
        v[61] = x[C.kdeg]*y[V.E11G]
        v[62] = x[C.kdeg]*y[V.E11S]
        v[63] = x[C.kdeg]*y[V.E11R]
        v[64] = x[C.kdeg]*y[V.E11T]
        v[65] = x[C.V1] * y[V.ppMEKc] * y[V.ERKc] /  ( x[C.Km1] * (1 + y[V.pERKc] / x[C.Km2]) + y[V.ERKc] )
        v[66] = x[C.V2] * y[V.ppMEKc] * y[V.pERKc] /  ( x[C.Km2] * (1 + y[V.ERKc] / x[C.Km1]) + y[V.pERKc] )
        v[67] = x[C.V3] * y[V.pERKc] /  ( x[C.Km3] * (1 + y[V.ppERKc] / x[C.Km4]) + y[V.pERKc] )
        v[68] = x[C.V4] * y[V.ppERKc] /  ( x[C.Km4]* (1 + y[V.pERKc] / x[C.Km3]) + y[V.ppERKc] )
        v[69] = x[C.V5] * y[V.pERKn] /  ( x[C.Km5] * (1 + y[V.ppERKn] / x[C.Km6]) + y[V.pERKn] )
        v[70] = x[C.V6] * y[V.ppERKn] /  ( x[C.Km6] * (1 + y[V.pERKn] / x[C.Km5]) + y[V.ppERKn] )
        v[71] = x[C.KimERK] * y[V.ERKc] - x[C.KexERK] * (x[C.Vn]/x[C.Vc]) * y[V.ERKn]
        v[72] = x[C.KimpERK] * y[V.pERKc] - x[C.KexpERK] * (x[C.Vn]/x[C.Vc]) * y[V.pERKn]
        v[73] = x[C.KimppERK] * y[V.ppERKc] - x[C.KexppERK] * (x[C.Vn]/x[C.Vc]) * y[V.ppERKn]
        v[74] = x[C.kf_16_vegf]*y[V.TF_inact]*y[V.ppERKn]/(x[C.Kmf_16_vegf]+y[V.TF_inact])
        v[75] = x[C.kr_17_vegf]*y[V.TF_act]/(x[C.Kmr_17_vegf]+y[V.TF_act])
        v[76] = x[C.prod_mFMOD]*(y[V.TF_act])**x[C.n1_vegf]/((x[C.Km_18_vegf])**x[C.n1_vegf]+(y[V.TF_act])**x[C.n1_vegf])
        v[77] = x[C.degrad_mFMOD]*y[V.mFMOD]
        v[78] = x[C.prod_FMOD]*y[V.mFMOD]
        v[79] = x[C.act_PI3K]*y[V.TGFBR_act]*y[V.I] - x[C.inact_PI3K]*y[V.sigmaI]

    # Equations of the model
    # TGFb_model
        dydt[V.TGFb_inact] = - v[1]
        dydt[V.TGFb_act] = + v[1] - v[2] - v[3] + v[20]
        dydt[V.FMOD] = - v[2] + v[20]
        dydt[V.FMOD_complex] = + v[2] - v[20]
        dydt[V.TGFBR_inact] = - v[3] + v[4]
        dydt[V.TGFBR_act] = + v[3] - v[4]
        dydt[V.S2] = - v[5] + v[7]
        dydt[V.S3] = - v[8] + v[10]
        dydt[V.S4] = - v[13] + v[21]
        dydt[V.pS2] = + v[6] - v[7] + v[12]
        dydt[V.pS3] = + v[9] - v[10] + v[12]
        dydt[V.ppS2] = + v[5] - v[6] - v[11]
        dydt[V.ppS3] = + v[8] - v[9] - v[11]
        dydt[V.ppS2_ppS3] = + v[11] - v[12] - v[13] + v[21]
        dydt[V.ppS2_ppS3_S4] = + v[13] - v[23] - v[21] + v[22]
        dydt[V.mS7] = + v[14] - v[15]
        dydt[V.S7] = + v[16]
        dydt[V.mcFOS] = + v[17] - v[18]
        dydt[V.cFOS] = + v[19] - v[27] - v[23] + v[22]
        dydt[V.ppS2_ppS3_S4_cFOS] = + v[23] - v[22]
        dydt[V.mTHBS1] = + v[24] - v[25]
        dydt[V.THBS1] = + v[26]
    # VEGF_model
        dydt[V.E] = (-v[28] - v[58])/VeVc
        dydt[V.E1] = -v[28] - v[56]
        dydt[V.E_E1] = v[28] - v[29] - v[29] - v[57] #VEGF_VEGFR
        dydt[V.E11] = v[29] - v[30] #VEGF_VEGFR_dimer
        dydt[V.E11P] = v[30] - v[31] - v[32] - v[33] - v[55] - v[60] #VEGF_pVEGFR_dimer
        dydt[V.G] = -v[31] - v[38] - v[51] + v[61] #Grb2
        dydt[V.S] = -v[32]  - v[39] + v[62] #Shc
        dydt[V.I] = - v[40] - v[79] #PI3K
        dydt[V.R] = -v[33]  - v[41] + v[63] #RasGAP
        dydt[V.O] = -v[36] - v[49] - v[52] - v[54] #SOS
        dydt[V.A] = -v[37] - v[42] - v[50] - v[53] #GAB1
        dydt[V.E11G] = v[31] - v[61]
        dydt[V.E11S] = v[32] - v[62]
        dydt[V.E11R] = v[33] - v[63]
        dydt[V.sigmaG] = v[31]  - v[36] - v[37] + v[38] - v[61]
        dydt[V.sigmaS] = v[32]  - v[34] + v[39] - v[62]
        dydt[V.sigmaI] = v[40] + v[79]
        dydt[V.sigmaR] = v[33]  + v[41] - v[46] - v[63]
        dydt[V.sigmaA] = -v[35] + v[37] + v[42] + v[53]
        dydt[V.sigmaSP] = v[34] - v[38]
        dydt[V.sigmaAP] = v[35] - v[39] - v[40] - v[41] - v[59]
        dydt[V.sigmaG_O] = v[36] - v[53]
        dydt[V.sigmaG_A] = v[37] - v[54]
        dydt[V.sigmaSP_G] = v[38]
        dydt[V.sigmaAP_S] = v[39]
        dydt[V.sigmaAP_I] = v[40]
        dydt[V.sigmaAP_R] = v[41]
        dydt[V.P3_A] = v[42] - v[51]
        dydt[V.P2] = -v[44]
        dydt[V.P3] = -v[42] + v[44]
        dydt[V.Akt] = -v[43]
        dydt[V.Aktstar] = v[43]
        dydt[V.RsD] = -v[45]
        dydt[V.RsT] = v[45]
        dydt[V.sigmaRP] = v[46]
        dydt[V.Raf] = -v[47]
        dydt[V.Rafstar] = v[47]
        dydt[V.MEK] = -v[48]
        dydt[V.ppMEKc] = v[48]
        dydt[V.OP] = v[49]
        dydt[V.AP] = v[50]
        dydt[V.A_sigmaG_O] = v[53] + v[54]
        dydt[V.sigmaA_G] = v[51] - v[52]
        dydt[V.sigmaA_G_O] = v[52]
        dydt[V.sigmaO] = v[36] + v[52] + v[54]
        dydt[V.T] = -v[55]  - v[59] + v[64]
        dydt[V.E11T] = v[55] - v[64]
        dydt[V.sigmaT] = v[55]  + v[59] - v[64]
        dydt[V.E1_PT] = v[56] - v[58]
        dydt[V.E_E1_PT] = v[57] + v[58]
        dydt[V.sigmaAP_T] = v[59]
        dydt[V.fint] = x[C.a98]*(-y[V.fint] + x[C.b98])
        dydt[V.ERKc] = -v[65] + v[67] - v[71]
        dydt[V.ERKn] = v[69] + v[71]*(x[C.Vc]/x[C.Vn])
        dydt[V.pERKc] = v[65] - v[66] -v[67] +v[68]-v[72]
        dydt[V.pERKn] = -v[69] + v[70] + v[72]*(x[C.Vc]/x[C.Vn])
        dydt[V.ppERKc] =  v[66] - v[68] - v[73]
        dydt[V.ppERKn] = -v[70] + v[73]*(x[C.Vc]/x[C.Vn])
        dydt[V.TF_inact]     = + v[75] - v[74]
        dydt[V.TF_act]     = + v[74] - v[75]
        dydt[V.mFMOD]     = + v[76] - v[77]
        dydt[V.FMOD]     = - v[2] + v[20] + v[78]

    # Best fit parameter
        x[C.kf_1_TGFbeta] = 4.923e-01
        x[C.Kmf_1_TGFbeta] = 2.529e+00
        x[C.k_on_FMOD] = 2.168e+02
        x[C.k_off_FMOD] = 2.045e+00
        x[C.Rec_act] = 5.970e-01
        x[C.pRec_debind] = 8.082e-03
        x[C.S2tot] = 6.000e-02
        x[C.S3tot] = 3.800e-01
        x[C.S4tot] = 4.400e-03
        x[C.kf_2_TGFbeta] = 4.664e-01
        x[C.Kmf_2_TGFbeta] = 4.889e+00
        x[C.k_inhibit_TGF] = 1.569e+00
        x[C.S_dephosphos] = 1.001e-01
        x[C.S_dephos] = 1.947e+00
        x[C.kf_3_TGFbeta] = 9.264e-01
        x[C.Kmf_3_TGFbeta] = 1.004e-01
        x[C.k_on_ppS2_ppS3] = 8.995e-01
        x[C.k_on_ppS2_ppS3_S4] = 9.071e-01
        x[C.k_off_ppS2_ppS3_S4] = 3.579e-01
        x[C.prod_mS7] = 2.069e-01
        x[C.n1_TGF] = 1.000e+00
        x[C.Km_1_TGF] = 2.299e-01
        x[C.mS7_turn] = 8.564e+00
        x[C.prod_S7] = 2.258e-01
        x[C.prod_mcFOS] = 1.848e+01
        x[C.n2_TGF] = 1.000e+00
        x[C.Km_2_TGF] = 2.049e-02
        x[C.mcFOS_turn] = 2.056e-02
        x[C.prod_cFOS] = 1.134e-01
        x[C.k_on_ppS2_ppS3_S4_cFOS] = 6.391e-01
        x[C.k_off_ppS2_ppS3_S4_cFOS] = 2.785e-01
        x[C.prod_mTHBS1] = 2.157e+02
        x[C.n4_TGF] = 1.000e+00
        x[C.Km_4_TGF] = 1.102e-01
        x[C.THBS1_turn] = 5.464e-03
        x[C.prod_THBS1] = 3.524e+00
        x[C.w_THBS1] = 1.234e-01
        x[C.w_TGFBR1] = 2.282e-01
        x[C.w_TGFBR2] = 3.128e-01
        x[C.w_SMAD7] = 1.021e-01
        x[C.w_cFOS] = 1.047e-01
        x[C.degrad_cFOS] = 1.000e-01
        x[C.VmaxPY] = 2.209e+03
        x[C.KmPY] = 4.197e+02
        x[C.kdeg] = 2.220e-02
        x[C.kf47] = 2.441e+02
        x[C.Vmaxr47] = 3.040e+02
        x[C.Kmf47] = 6.986e+01
        x[C.Kmr47] = 1.218e+03
        x[C.kf48] = 7.758e+00
        x[C.Kmf48] = 1.543e+03
        x[C.Kmr48] = 3.083e+03
        x[C.PTEN] = 7.689e+01
        x[C.kf49] = 1.340e+02
        x[C.kr49] = 4.429e+03
        x[C.Kmf49] = 1.923e+02
        x[C.Kmr49] = 8.206e+02
        x[C.Kmr49b] = 5.209e+02
        x[C.kr49b] = 5.056e+02
        x[C.kf51] = 8.832e-01
        x[C.Vmaxr51] = 2.239e+01
        x[C.Kmf51] = 5.166e+03
        x[C.Kmrb51] = 8.959e+03
        x[C.kf52] = 3.306e+00
        x[C.Vmaxr52] = 1.320e+02
        x[C.Kmf52] = 5.670e+01
        x[C.Kmr52] = 1.965e+03
        x[C.kf54] = 2.758e-02
        x[C.Vmaxr54] = 2.994e+02
        x[C.Kmf54] = 1.760e+03
        x[C.Kmr54] = 1.338e+02
        x[C.kf55] = 3.059e-01
        x[C.Vmaxr55] = 1.347e+02
        x[C.Kmf55] = 8.731e+01
        x[C.Kmr55] = 1.127e+03
        x[C.kf38] = 6.409e+01
        x[C.kf39] = 1.249e+02
        x[C.kf50] = 1.029e+02
        x[C.a98] = 1.685e-02
        x[C.b98] = 7.300e-01
        x[C.koff46] = 3.845e-01
        x[C.EGF_off] = 8.144e-03
        x[C.koff4] = 2.320e-01
        x[C.koff16] = 2.271e+00
        x[C.koff17] = 8.975e+00
        x[C.koff18] = 6.361e+00
        x[C.koff40] = 7.349e+00
        x[C.koff41] = 1.709e+00
        x[C.koff42] = 1.779e+00
        x[C.koff43] = 7.319e-01
        x[C.koff44] = 1.339e-01
        x[C.koff45] = 1.650e+01
        x[C.koff57] = 4.455e+00
        x[C.koff58] = 4.199e+00
        x[C.koff59] = 1.170e+01
        x[C.koff60] = 8.973e-01
        x[C.kPTP10] = 1.457e+02
        x[C.koff73] = 4.650e-01
        x[C.kPTP38] = 1.861e+02
        x[C.kPTP39] = 5.194e+01
        x[C.koff88] = 2.285e+01
        x[C.kPTP50] = 2.738e+02
        x[C.kf81] = 1.262e+01
        x[C.Vmaxr81] = 7.175e+01
        x[C.Kmf81] = 1.533e+03
        x[C.Kmr81] = 1.676e+03
        x[C.kf84] = 1.951e+00
        x[C.Vmaxr84] = 5.420e+02
        x[C.Kmf84] = 1.027e+03
        x[C.Kmr84] = 5.849e+02
        x[C.kcon49] = 2.571e+01
        x[C.kon1] = 2.199e-05
        x[C.kon86] = 1.320e-02
        x[C.kon4] = 2.444e+00
        x[C.kf10] = 1.122e+00
        x[C.kon16] = 3.510e-03
        x[C.kon17] = 3.378e-02
        x[C.kon18] = 1.022e-02
        x[C.kon73] = 3.401e-02
        x[C.kon40] = 3.385e-02
        x[C.kon41] = 2.294e-03
        x[C.kon42] = 3.881e-03
        x[C.kon43] = 5.142e-03
        x[C.kon44] = 2.135e-02
        x[C.kon45] = 1.039e-03
        x[C.kon88] = 1.085e-03
        x[C.kon46] = 1.315e-01
        x[C.kon57] = 4.061e-03
        x[C.kon58] = 2.149e-01
        x[C.kon59] = 5.092e-03
        x[C.kon60] = 3.622e-04
        x[C.V1] = 5.407e-01
        x[C.Km1] = 4.461e+02
        x[C.V2] = 2.200e-01
        x[C.Km2] = 3.500e+02
        x[C.V3] = 7.200e-01
        x[C.Km3] = 1.600e+02
        x[C.V4] = 6.480e-01
        x[C.Km4] = 6.000e+01
        x[C.V5] = 9.121e+00
        x[C.Km5] = 3.254e+00
        x[C.V6] = 9.121e+00
        x[C.Km6] = 3.254e+00
        x[C.KimERK] = 1.200e-02
        x[C.KexERK] = 1.800e-02
        x[C.KimpERK] = 1.200e-02
        x[C.KexpERK] = 1.800e-02
        x[C.KimppERK] = 1.100e-02
        x[C.KexppERK] = 1.300e-02
        x[C.Vn] = 2.200e-01
        x[C.Vc] = 9.400e-01
        x[C.act_PI3K] = 5.185e+00
        x[C.inact_PI3K] = 2.077e-01
        x[C.kf_16_vegf] = 1.792e-01
        x[C.Kmf_16_vegf] = 1.639e-03
        x[C.kr_17_vegf] = 4.213e+00
        x[C.Kmr_17_vegf] = 8.677e+00
        x[C.prod_mFMOD] = 2.758e+01
        x[C.Km_18_vegf] = 1.533e+02
        x[C.n1_vegf] = 1.000e+00
        x[C.degrad_mFMOD] = 7.070e-01
        x[C.prod_FMOD] = 3.175e-01
        x[C.w_VEGFR1] = 6.361e-01
        x[C.w_VEGFR2] = 1.185e+01
        x[C.w_G] = 6.101e+00
        x[C.w_SHC1] = 1.996e+00
        x[C.w_SHC2] = 9.208e-01
        x[C.w_SHC3] = 1.451e+01
        x[C.w_SHC4] = 4.376e+00
        x[C.w_PIK3CA] = 3.317e+01
        x[C.w_PIK3CB] = 8.983e+00
        x[C.w_PIK3CD] = 4.935e-01
        x[C.w_PIK3CG] = 4.971e-01
        x[C.w_PTEN] = 1.502e+00
        x[C.w_RASA1] = 3.196e+01
        x[C.w_RASA2] = 8.671e+00
        x[C.w_RASA3] = 6.741e+00
        x[C.w_SOS1] = 1.192e+01
        x[C.w_SOS2] = 1.055e+01
        x[C.w_A] = 7.601e+00
        x[C.w_AKT1] = 2.547e+01
        x[C.w_AKT2] = 1.378e+00
        x[C.w_HRAS] = 8.886e-01
        x[C.w_KRAS] = 7.941e-01
        x[C.w_NRAS] = 1.985e+00
        x[C.w_ARAF] = 1.422e+01
        x[C.w_BRAF] = 3.225e+00
        x[C.w_RAF1] = 9.078e+00
        x[C.w_MAP2K1] = 2.270e+01
        x[C.w_MAP2K2] = 3.834e+00
        x[C.w_T] = 8.234e+00
        x[C.w_MAPK1] = 1.341e+00
        x[C.w_MAPK3] = 7.190e-01
        x[C.w_FMOD] = 1.214e+00

    # None zero initial values for 'Control' condition
    # y0[V.TGFb_act] =0 for 'Control'
        y0[V.TGFb_inact] = 0.013
        y0[V.TGFBR_inact] = 4.475e+01
        y0[V.S2] = 6.000e-02
        y0[V.S3] = 3.800e-01
        y0[V.S4] = 4.400e-03
        y0[V.mS7] = 2.105e+00
        y0[V.mcFOS] = 1.225e-01
        y0[V.mTHBS1] = 5.581e+02
        y0[V.E] = 1.000e+01
        y0[V.E1] = 6.482e+01
        y0[V.G] = 7.836e+02
        y0[V.S] = 8.693e+02
        y0[V.I] = 6.943e+02
        y0[V.R] = 3.592e+03
        y0[V.T] = 6.927e+02
        y0[V.O] = 5.132e+02
        y0[V.A] = 5.389e+01
        y0[V.P2] = 4.811e+01
        y0[V.Akt] = 2.212e+03
        y0[V.RsD] = 3.497e+02
        y0[V.Raf] = 1.958e+03
        y0[V.MEK] = 2.098e+03
        y0[V.ERKc] = 1.595e+02
        y0[V.TF_inact] = 6.616e+01
        y0[V.mFMOD] = 2.536e+01
    # None zero initial values for 'TGFβ1' condition
    # y0[V.TGFb_act] = 0.0902 for 'TGFβ1'
        y0[V.TGFb_inact] = 0.013
        y0[V.TGFb_act] =0.0902
        y0[V.TGFBR_inact] = 4.475e+01
        y0[V.S2] = 6.000e-02
        y0[V.S3] = 3.800e-01
        y0[V.S4] = 4.400e-03
        y0[V.mS7] = 2.105e+00
        y0[V.mcFOS] = 1.225e-01
        y0[V.mTHBS1] = 5.581e+02
        y0[V.E] = 1.000e+01
        y0[V.E1] = 6.482e+01
        y0[V.G] = 7.836e+02
        y0[V.S] = 8.693e+02
        y0[V.I] = 6.943e+02
        y0[V.R] = 3.592e+03
        y0[V.T] = 6.927e+02
        y0[V.O] = 5.132e+02
        y0[V.A] = 5.389e+01
        y0[V.P2] = 4.811e+01
        y0[V.Akt] = 2.212e+03
        y0[V.RsD] = 3.497e+02
        y0[V.Raf] = 1.958e+03
        y0[V.MEK] = 2.098e+03
        y0[V.ERKc] = 1.595e+02
        y0[V.TF_inact] = 6.616e+01
        y0[V.mFMOD] = 2.536e+01
```
